# Cell membranes sustain phospholipid imbalance via cholesterol asymmetry

**DOI:** 10.1101/2023.07.30.551157

**Authors:** Milka Doktorova, Jessica L. Symons, Xiaoxuan Zhang, Hong-Yin Wang, Jan Schlegel, Joseph H. Lorent, Frederick A. Heberle, Erdinc Sezgin, Edward Lyman, Kandice R. Levental, Ilya Levental

## Abstract

Membranes are molecular interfaces that compartmentalize cells to control the flow of nutrients and information. These functions are facilitated by diverse collections of lipids, nearly all of which are distributed asymmetrically between the two bilayer leaflets. Most models of biomembrane structure and function often include the implicit assumption that these leaflets have similar abundances of phospholipids. Here, we show that this assumption is generally invalid and investigate the consequences of lipid abundance imbalances in mammalian plasma membranes (PM). Using quantitative lipidomics, we discovered that cytoplasmic leaflets of human erythrocyte membranes have >50% overabundance of phospholipids compared to exoplasmic leaflets. This imbalance is enabled by an asymmetric interleaflet distribution of cholesterol, which regulates cellular cholesterol homeostasis. These features produce unique functional characteristics, including low PM permeability and resting tension in the cytoplasmic leaflet that regulates protein localization. These largely overlooked aspects of membrane asymmetry represent an evolution of classic paradigms of biomembrane structure and physiology.

## Introduction

The plethora of molecular processes at the cellular plasma membrane (PM) is enabled by a diverse set of membrane proteins, but also by the membrane’s lipid matrix. One characteristic feature of eukaryotic PMs is their compositional asymmetry, i.e. drastically different phospholipid (PL) compositions of the two bilayer leaflets (Bretscher, 1972; Devaux, 1991; Verkleij et al., 1973). This asymmetry is established and maintained by ATP-driven enzymes that directionally flip PLs across the bilayer (Nagata et al., 2020; Sakuragi and Nagata, 2023). Regulation of these processes produces leaflets with distinct compositions and physical properties, including relatively tightly packed exoplasmic leaflets and more charged and fluid cytoplasmic leaflets (Lorent et al., 2020). The interleaflet distribution of cholesterol, a major PM component, remains elusive amid conflicting reports (Steck and Lange, 2018). While specific lipid compositions differ between organisms and cell types, structural features of protein transmembrane domains shared across diverse organisms suggest that most eukaryote PMs are asymmetric (Lorent et al., 2020).

PM asymmetry has been associated with many cellular functions, most notably apoptosis, wherein cell surface exposure of the cytoplasmic leaflet lipid phosphatidylserine (PS) is a central hallmark (Nagata et al., 2020). PS exposure on platelets is also required for activation of blood coagulation enzymes (Sakuragi and Nagata, 2023). This release of lipid asymmetry is often referred to as ‘scrambling’ and is mediated by ATP-independent lipid channels (Sakuragi and Nagata, 2023). PM asymmetry has also been implicated in cell-cell fusion (van den Eijnde et al., 2001), viral entry (Mercer and Helenius, 2008), and intracellular signaling (Elliott et al., 2005); however, in most cases, it is not lipid asymmetry *per se*, but rather its loss that is associated with cellular responses (Bevers and Williamson, 2016; Doktorova et al., 2020a). Thus, despite extensive evidence for the prominent chemical and physical asymmetries of PM bilayers, it is yet unknown why most cells expend significant metabolic resources to maintain membrane lipid distributions far away from their symmetric equilibrium.

Here, integrating cellular, biochemical, and computational approaches, we report the discovery of a largely overlooked structural and functional aspect of membrane asymmetry in cells: namely, a substantial steady-state phospholipid abundance imbalance between PM leaflets. This imbalance is enabled by highly asymmetric distributions of cholesterol, which can rapidly redistribute to buffer leaflet stresses. These unexpected phospholipid and cholesterol distributions produce unique membrane properties that illuminate the functional roles of PM asymmetry and its regulated release.

## Results

### Lipid imbalance in plasma membranes

Recently, we combined enzymatic digestion of exoplasmic leaflet lipids with shotgun mass spectrometry to quantify the molecularly detailed compositional asymmetry of nearly all PL classes in the human erythrocyte PM (Lorent et al., 2020). Here, we completed the characterization of the known erythrocyte PM lipidome by quantifying glycosphingolipids (GSLs) and cholesterol abundances. We found that GSLs constitute ∼15 mol% of exoplasmic leaflet lipids (Figures 1A, S1B) while cholesterol is present at 40 (± 1) mol% of PM lipids (Figure 1C). Due to its rapid transbilayer diffusion, cholesterol’s interleaflet distribution is not accessible by enzymatic approaches (Steck and Lange, 2018).

**Fig. 1.**
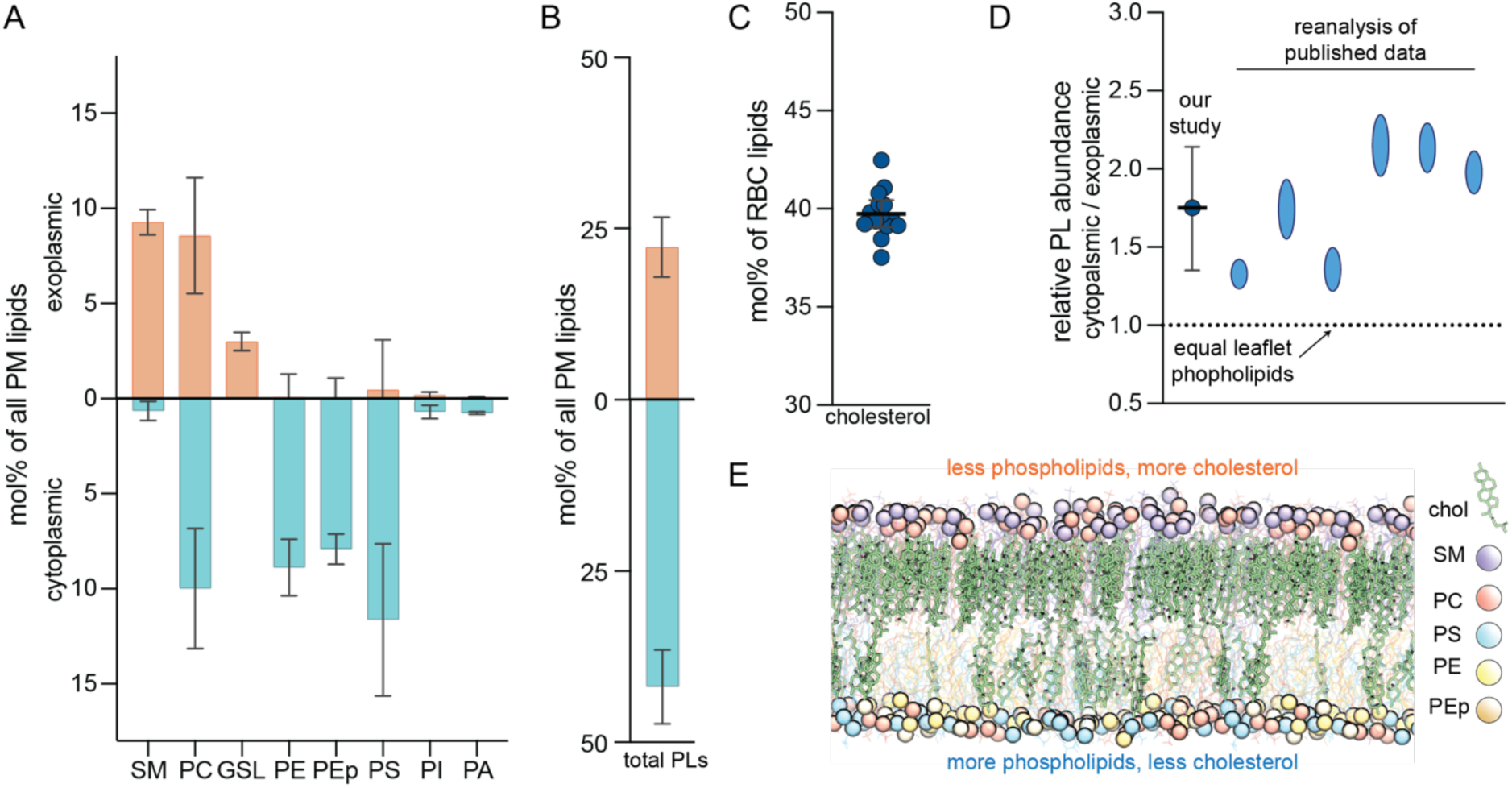
The exoplasmic leaflet of erythrocyte PMs has fewer phospholipids. (**A**) Shotgun mass spectrometry quantifications of asymmetric distribution of major lipid classes in the erythrocyte PM. Exoplasmic and cytoplasmic lipidomes are determined by phospholipase treatment of live cells and include species with a variety of chain compositions. Controls and validations detailed in (Lorent et al., 2020). Values represent mol% of lipid class as a percentage of PM lipids, with interleaflet distribution shown in orange/blue bars. Average ± SD from >3 independent experiments. SM sphingomyelin, PC phosphatidylcholine, GSL glycosphingolipids, PE phosphatidylethanolamine, PEp PE plasmalogen, PS phosphatidylserine, PI phosphatidylinositol, PA phosphatidic acid. Minor classes (<1% not shown). (**B**) Total phospholipid (including GSLs) abundances of the two PM leaflets obtained by summing the corresponding leaflet components in (A). (**C**) Cholesterol measured by lipidomics as a mol% of all erythrocyte lipids. The mean is 40 (± 1) mol% of PM lipids, however its transbilayer distribution cannot be resolved by this analysis. (**D**) Relative total phospholipid (PL) abundances between the two PM leaflets of erythrocytes from this work and from re-analysis of previous reports (Table S1). (**E**) Snapshot of an equilibrated atomistic computational model of the erythrocyte PM with overabundance of PLs in the cytoplasmic leaflet (Cyto+ model, Table S5). The membrane contains 40 mol% cholesterol, which converges to an asymmetric distribution with ∼3-fold enrichment in the exoplasmic leaflet. Lipid classes are color-coded (phosphorous atoms shown as spheres; cholesterol in green). Structures of PLs used in simulations shown in Figure S2.

Internal standards in mass spectrometry measurements allow quantification of individual lipid species, which can then be used to compare total PL abundances between the two PM leaflets. Surprisingly, we found that cytoplasmic leaflet phospholipids are 1.5–2.3-fold more abundant than those resident in the exoplasmic leaflet (Figure 1B) (for simplicity, ‘PLs’ here includes GSLs despite their not having a phosphate group). Such large differences in PL abundances between leaflets were unexpected because monolayer areas in a bilayer must match. This imbalance cannot be explained by membrane components not included in this analysis, such as GPI-anchored proteins, phosphoinositides, or protein transmembrane domains (see SI text, section 1.5 for discussion). Notably, differences in interleaflet PL abundances are not unprecedented, with similar imbalances evident when the foundational data that originally reported lipid asymmetry (Rawyler et al., 1985; Renooij et al., 1974; Van der Schaft et al., 1987; Verkleij et al., 1973; Zwaal et al., 1975) is reanalyzed (Figure 1D and Table S1).

### Phospholipid imbalance is sustained by cholesterol

To investigate the previously dismissed (Rawyler et al., 1985) feasibility of a membrane with extensive PL abundance imbalance, we built an atomically detailed computational PM model with lipid compositions of the two leaflets guided by lipidomics data (Table S5). The model was built with more phospholipids in the cytoplasmic leaflet (relative to the exoplasmic one) and was termed Cyto+ (Figure 1E). The simulated Cyto+ membrane also contained 40 mol% cholesterol, whose initial interleaflet distribution was adjusted to ensure matching leaflet surface areas. Having the unique ability to quickly flip-flop between leaflets (Bruckner et al., 2009; Lange et al., 1981), cholesterol equilibrated its distribution during the 17 *μ*s-long trajectory, with 77% of cholesterol molecules accumulating in the PL-poor exoplasmic leaflet (Figure 1E). Thus, we hypothesized that cholesterol can enrich in PL-poor leaflets, enabling large interleaflet phospholipid imbalances.

To test this hypothesis, we simulated a series of coarse-grained asymmetric bilayers with fixed PL leaflet imbalance (cytoplasmic leaflet with 35% more PLs than exoplasmic) and cholesterol abundances ranging from 10 to 50 mol% (Figure 2A, Table S2). In all these simulations, cholesterol equilibrated to an asymmetric distribution, accumulating in the PL-poor exoplasmic leaflet (Figure S3). Notably, low cholesterol concentrations were unable to sustain stable, flat bilayer morphologies; instead, the systems adopted unusual, nonlamellar configurations (Figure 2A). In contrast, membranes with ≥30 mol% cholesterol produced stable, flat bilayers, confirming that cholesterol (in sufficient abundance) enables tolerance for large PL imbalances via its asymmetric interleaflet distribution.

**Fig. 2.**
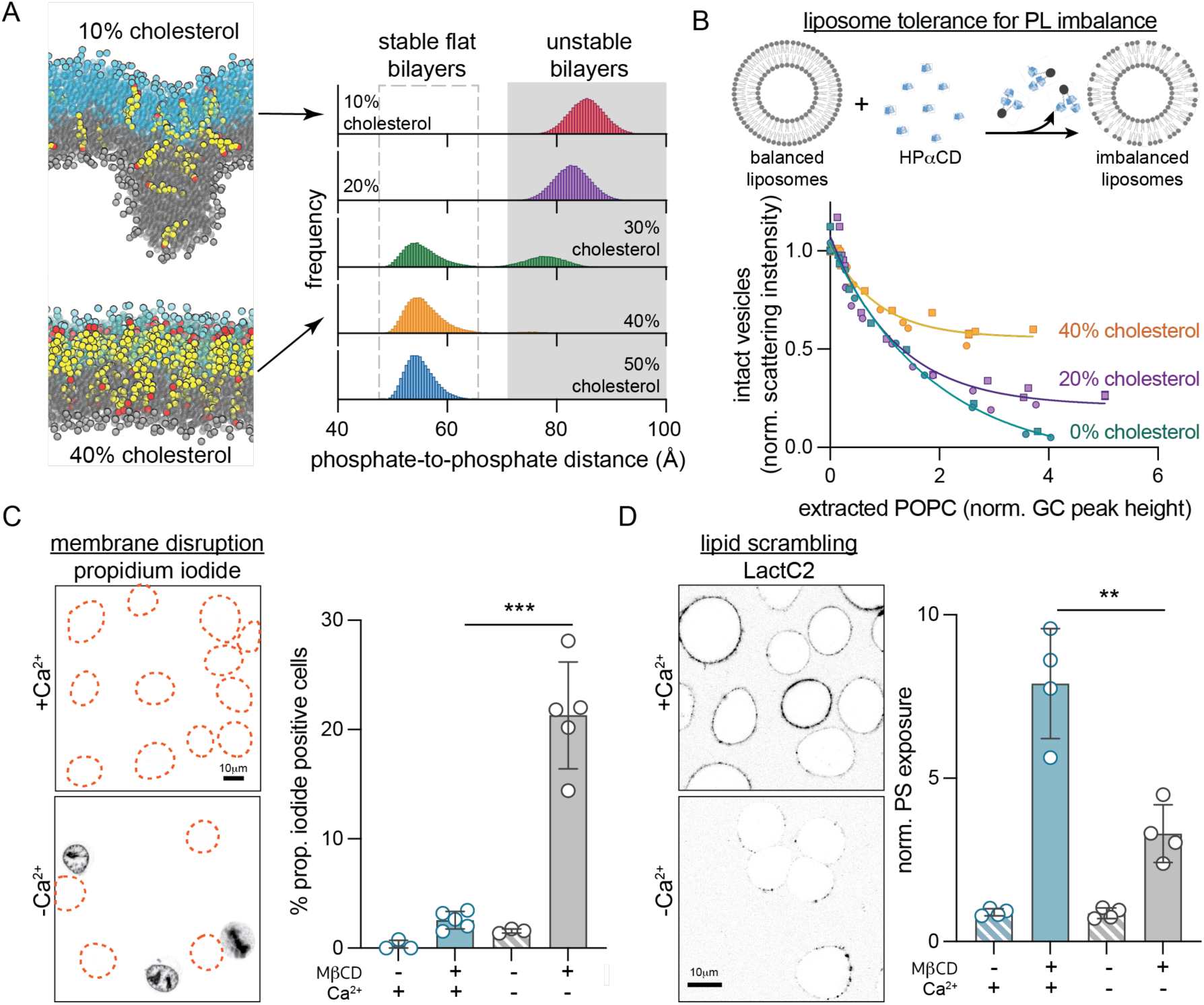
Cholesterol imparts tolerance for phospholipid abundance imbalance between membrane leaflets. (**A**) Coarse-grained simulations of a PM model with 35% more PLs in the cytoplasmic leaflet and varying cholesterol abundance (10-50%, Table S2). Histograms show the distributions of maximal distances between lipid phosphate groups in the two leaflets. Values >70 Å indicate unstable bilayers with non-lamellar morphology. Representative snapshots from simulated bilayers with 10 and 40% cholesterol shown for reference (exoplasmic leaflet PL phosphate groups shown in cyan, cytoplasmic in gray, cholesterol is yellow with hydroxyl headgroup in red). (**B**) Extraction of external leaflet PLs from extruded liposomes with 0, 20 and 40 mol% cholesterol. POPC is extracted from the external leaflet by HP*α*CD, which induces PL imbalance between leaflets causing membrane instability and liposome destruction. Amount of extracted lipid is quantified by GC/MS, while the corresponding fraction of intact vesicles is measured by light scattering (data from two independent experiments). (**C**) Extraction of cholesterol compromises PM integrity, evidenced by propidium iodide (PI) staining of nuclei, if lipid scrambling is suppressed by chelating Ca^2+^. Outlines represent PI-negative cells. (**D**) Extraction of cholesterol from the PM of RBL cells with M*β*CD leads to scrambling of PM lipids evidenced by exposure of PS on the cell surface (binding of external PS-marker, LactC2-mClover), while suppressing scrambling reduces PS exposure. Average intensity of LactC2-mClover on the PM (PS exposure) normalized to -M*β*CD is shown on the right. Data shown in (C) and (D) is from average ± SD from 3-5 independent experiments. **p<0.01, ***p<0.001.

To experimentally evaluate this inference, we measured the tolerance of liposomes to PL imbalance via single-sided lipid extraction (Figure 2B). Large unilamellar vesicles (composed of POPC with 0-40 mol% cholesterol) were titrated with 2-hydroxypropyl-*α*-cyclodextrin (HP*α*CD), a cyclic polysaccharide that avidly extracts PLs but not cholesterol from the outward-facing membrane leaflet (Huang and London, 2013). More HP*α*CD extracts more PL, inducing larger leaflet imbalances, which ultimately destroy the liposomes, evidenced by decreased light scattering of the suspensions (Figure S4). By directly quantifying the amount of extracted POPC (i.e. induced interleaflet PL abundance imbalance) by gas chromatography, we observed that cholesterol protects vesicles from destruction in a dose-dependent fashion, consistent with simulations (Figure 2B).

Our lipidomics measurements indicate that cholesterol comprises ∼40 mol% of PM lipids in erythrocytes, consistent with previous estimates in mammalian PMs (Steck and Lange, 2018). Our simulations and synthetic membrane experiments suggest that such high cholesterol levels are required to sustain large PL imbalances (Figure 2A–B). Thus, we hypothesized that extracting cholesterol from cell PMs would destabilize the bilayer and compromise membrane integrity. To facilitate imaging and genetic manipulations, cellular experiments were conducted with cultured rat basophil leukemia (RBL) cells. We selectively extracted cholesterol from RBL PMs with methyl-*β*-cyclodextrin (M*β*CD), which depletes both PM leaflets through cholesterol’s rapid transbilayer movement (Zidovetzki and Levitan, 2007), but does not extract PLs (McGraw et al., 2019). Contrary to our prediction, membrane integrity was not affected by M*β*CD, as revealed by lack of propidium iodide (PI) staining (Figure 2C) or permeability to dextran polymers (Figure S5). Rather, as previously reported (Arashiki et al., 2016), cholesterol extraction induced rapid PL scrambling evidenced by externalization of inner-leaflet PS (Figure 2D, S5A), which likely serves to eliminate any pre-existing PL imbalance between leaflets. To suppress scrambling and thereby isolate the effects of PL imbalance, we relied on the calcium dependence of the most widely characterized scramblase TMEM16F (Sakuragi and Nagata, 2023). Indeed, treatment of RBL cells with M*β*CD under Ca^2+^-free conditions significantly reduced PS exposure (Figure 2D, S5A). Strikingly, this treatment also disrupted membrane integrity, revealed by robust staining of nuclei with PI (Figure 2C, S5B) and leakage of dextran into the cytoplasm (Figure S5C). These results reveal that in response to cholesterol extraction, PLs are redistributed between PM leaflets to preserve membrane integrity. If this mechanism is inhibited, membrane integrity is lost, supporting the notion that cholesterol imparts tolerance for interleaflet PL abundance imbalance in living cells.

### Cholesterol distribution in the PM

Our Cyto+ model indicates that cholesterol can balance excess PLs in the cytoplasmic leaflet by accumulating in the exoplasmic leaflet (Figure 1E). PL imbalance in the PM can be established by the activity of enzymes termed flippases, which use ATP to translocate aminophospholipids (PS and PE) from the exoplasmic to the cytosolic leaflet against their concentration gradient (Sakuragi and Nagata, 2023). In the absence of compensatory lipid movement, active lipid flipping from the outer to the inner leaflet would under-populate the former and over-populate the latter, creating tension in the under-populated leaflet and compression in the over-populated one due to the requirement that leaflet areas match despite different lipid abundances. This effect was confirmed by a series of simulations in which increasing numbers of PS lipids were flipped into the ‘inner’ leaflet (Figure 3A). We also found that the resulting stresses could be alleviated by cholesterol spontaneously ‘flopping’ to the outer leaflet, consistent with previous reports of rapidly flipping lipids buffering leaflet tension (Bruckner et al., 2009; Miettinen and Lipowsky, 2019) (Figure 3A).

**Fig. 3.**
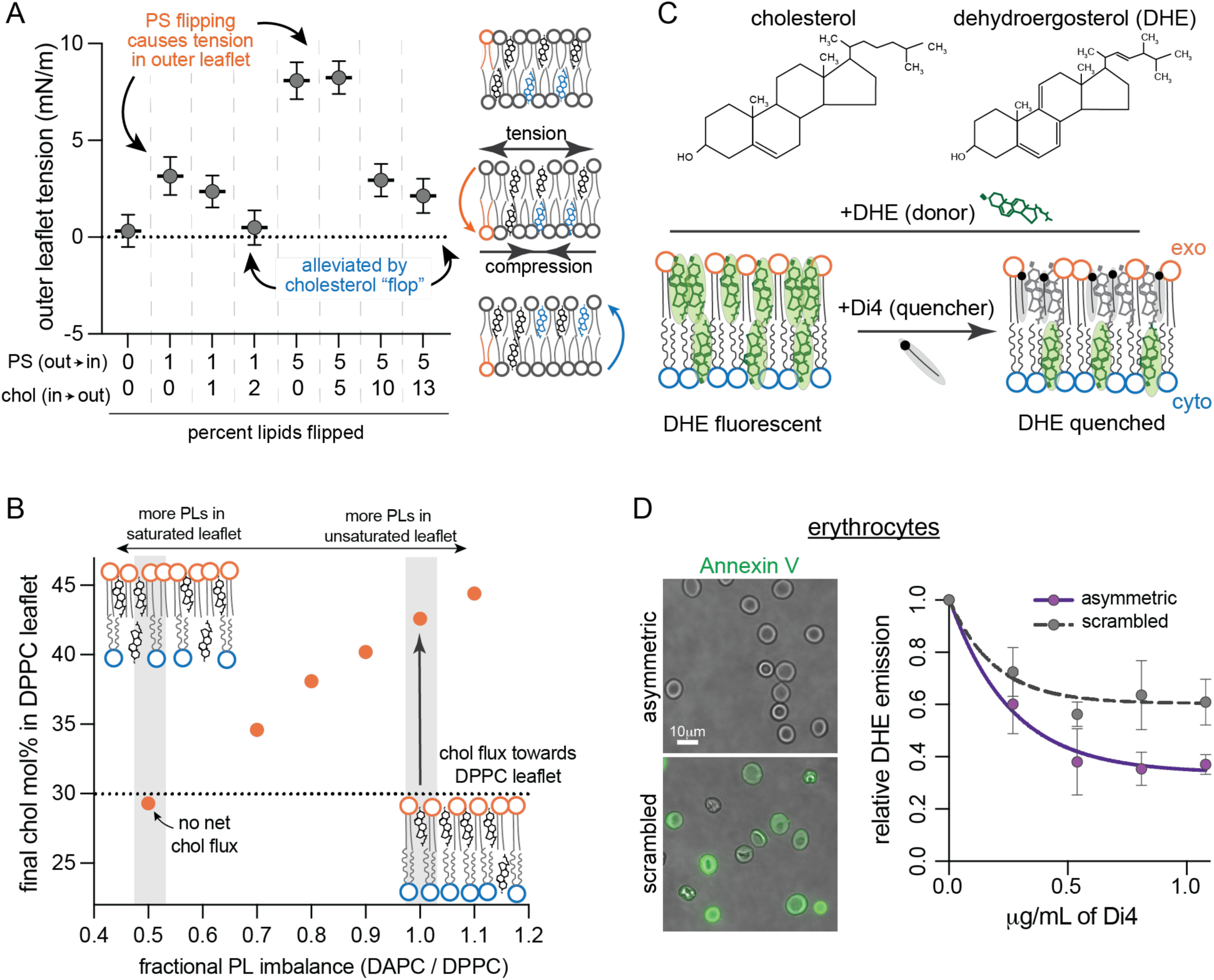
Cholesterol interleaflet distribution in model and cell membranes. (**A**) Leaflet tension (differential stress) in the outer leaflet of simulated bilayers composed of POPS (PS), POPC and cholesterol (Chol) modeling various extents of PS and Chol flipping. Atomistic bilayers were constructed with the indicated percentage of PS lipids effectively fl**i**pped from the outer leaflet to the inner leaflet, and Chol molecules fl**<u>o</u>**pped from the inner to the outer leaflet (Table S3). Outer leaflet tension calculated from equilibrated lateral pressure distributions; inner leaflet is under an equal magnitude compression. (**B**) Cholesterol distribution in coarse-grained asymmetric bilayers composed of a fully saturated outer leaflet (DPPC) with fixed number of PLs opposing a highly unsaturated inner leaflet (DAPC) with varying PL abundance (Table S4). The fractional imbalance of DAPC-to-DPPC lipids goes from underpopulated DAPC leaflet (left) to underpopulated DPPC leaflet (right). Chol was initiated at 30 mol% in each leaflet. Simulations were run for 10 µs allowing Chol to equilibrate between the two leaflets; the equilibrated cholesterol concentrations in the outer leaflet are shown. Schematics of the equilibrated relative lipid distributions are shown for comparison. (**C**) Schematic illustration of experimental approach for measuring Chol interleaflet distribution in erythrocytes. Minor fraction (<10%) of cholesterol in the erythrocyte PM is exchanged with DHE. A quencher, Di4, is added externally leading to its insertion into the outer leaflet. The fraction of DHE fluorescence quenched by Di4 provides a readout of relative DHE residence in the exoplasmic leaflet. (**D**) DHE fluorescence in erythrocyte membranes as a function of Di4 concentration. Data shown for untreated cells and cells whose PM lipids were scrambled with 100 μM PMA. Representative images show binding of PS-marker Annexin V before (top) and after (bottom) PMA treatment. Average ± SD for 3 independent experiments.

In addition to this stress-buffering role, cholesterol preferentially interacts with saturated versus unsaturated lipids, especially saturated sphingomyelins (Almeida, 2009). To incorporate this effect, we simulated a series of coarse-grained asymmetric membranes with one fully saturated (outer) and one highly unsaturated (inner) leaflet. Interleaflet PL abundances were varied, while cholesterol was initiated at 30 mol% in both leaflets (Figure 3B). In nearly all cases, we observed net cholesterol flux towards the outer leaflet (Figure 3B); e.g. when PLs were initially balanced, cholesterol equilibrates at >40 mol% in the outer leaflet (∼4-fold enriched relative to unsaturated leaflet) producing an over-populated outer leaflet even at the cost of generating, rather than alleviating, membrane stress (Figure S6). Net cholesterol flux could only be eliminated when the saturated lipid-rich ‘outer’ leaflet had 2-fold more PLs than the inner.

These findings reveal that cholesterol’s preference for certain lipid classes (saturated lipids, sphingolipids) combines with its tendency to populate under-packed leaflets to determine its ultimate transbilayer distribution, as discussed in previous theoretical and computational work (Allender et al., 2019; Hossein and Deserno, 2020; Varma and Deserno, 2022). In our lipidomics measurements, erythrocyte PM exoplasmic leaflets are more saturated (Figure S1A), rich in sphingolipids (Figure 1A), and PL-underpopulated (Figure 1B), making a strong prediction for cholesterol enrichment therein.

To test this prediction, we developed an assay to measure cholesterol distribution in a living cell PM. Erythrocytes were isolated from healthy donors and a small fraction of cholesterol was exchanged for the fluorescent cholesterol analog, dehydroergosterol (DHE, Figure 3C), which has similar chemical and biophysical properties (McIntosh et al., 2008; Pourmousa et al., 2014). We assayed the asymmetric distribution of DHE by fluorescence resonance energy transfer (FRET) to a quencher present only in the outer leaflet (Figure 3C). To that end, we added the fluorophore Di4 (which is charged and cannot flip between leaflets (Lorent et al., 2020)) into the erythrocyte outer leaflet. DHE and Di4 form a FRET pair with a Förster distance that is approximately half the bilayer thickness, and therefore exchange energy much more efficiently when they are in the same leaflet (section 5.2 in SI). Thus, Di4-induced quenching of DHE fluorescence provides a readout of the relative outer leaflet enrichment of DHE, as a proxy for cholesterol’s transbilayer distribution. Labeling conditions were tuned to ensure that overall sterol levels and membrane physical properties were unaffected (Figure S7) and that ∼50% of DHE is quenched in symmetric liposomes (Figure S8). Titration of Di4 into erythrocytes quenched DHE fluorescence, plateauing at ∼64% reduction of DHE emission (Figure 3D). Quenching of the majority of DHE fluorescence by an externally applied quencher is consistent with the prediction that cholesterol is enriched in the exoplasmic leaflet of a living cell PM. This enrichment was eliminated by releasing lipid asymmetry via treatment with phorbol myristate acetate (PMA) (Figure 3D), which scrambles erythrocyte PM phospholipids (Wesseling et al., 2016). Thus, computational predictions and experimental measurements suggest that cholesterol is enriched in the exoplasmic leaflet of human erythrocyte PMs. We discuss caveats and other previous studies of cholesterol asymmetry in the caption of Figure S8.

### Validation of the Cyto+ model

Our results suggest that the exoplasmic leaflet of human erythrocyte PMs is relatively saturated, PL-depleted, and cholesterol-rich, a model of PM structure which we termed Cyto+. A large, atomistic simulation of this model was stable over >17 µsec with cholesterol continuing to partition away from the cytoplasmic leaflet, producing an exoplasmic leaflet with high density, lipid packing, and acyl chain order (Figure S9). To validate this emerging model of lipid organization and membrane structure, we developed methods to quantitatively compare its structural and dynamical properties with those of living cell membranes. Previous measurements used the leaflet-specific fluorescence lifetime of Di4 (Lorent et al., 2020) to infer that the exoplasmic PM leaflet of mammalian cells is more tightly packed than the cytoplasmic. To examine how leaflet imbalances contribute to this observation, we calibrated experimental Di4 lifetimes to simulated lipid packing (i.e. area-per-lipid) in simple synthetic bilayers spanning a large range of lipid densities (Figure 4A). These two measures were highly correlated, allowing us to translate experimental Di4 lifetimes into a specific membrane structural parameter (area/lipid). We found that the exoplasmic PM leaflet of NIH 3T3 fibroblasts is ∼20% more tightly packed (40.9 ± 2.31 Å^2^/lipid) than the cytoplasmic leaflet (51.2 ± 2.22 Å^2^/lipid) (Figure 4C−D, hatched bars). We directly compared these measurements to three distinct models of PM organization: in addition to the Cyto+ model (Figure 1E) derived from lipidomics and supported by DHE FRET measurements (Figure 3C−D), we simulated two alternative models with identical leaflet lipid *compositions* but either equal (Equal) or greater (Exo+) PL *abundance* in the exoplasmic monolayer (Figure 4B, Table S5). The lipid packing of the exoplasmic leaflet of the Cyto+ model matched nearly perfectly with the experimental measurement, while Exo+ was least accurate (Figure 4C, left). Further, the large difference in area/lipid between the two leaflets was successfully captured only by the Cyto+ model (Figures 4C, S10). We also simulated a scrambled version of Cyto+ (i.e. same lipid composition, but all lipids equally distributed between leaflets) and observed an intermediate lipid density between the exoplasmic and cytoplasmic leaflets, which matched very closely to experimental measurements of the fibroblast PM scrambled by treatment with the calcium ionophore A23187 (Figure 4D). Thus, leaflet structural differences in living mammalian cells are most consistent with an overabundance of PLs in the cytoplasmic leaflet (Cyto+).

**Fig. 4.**
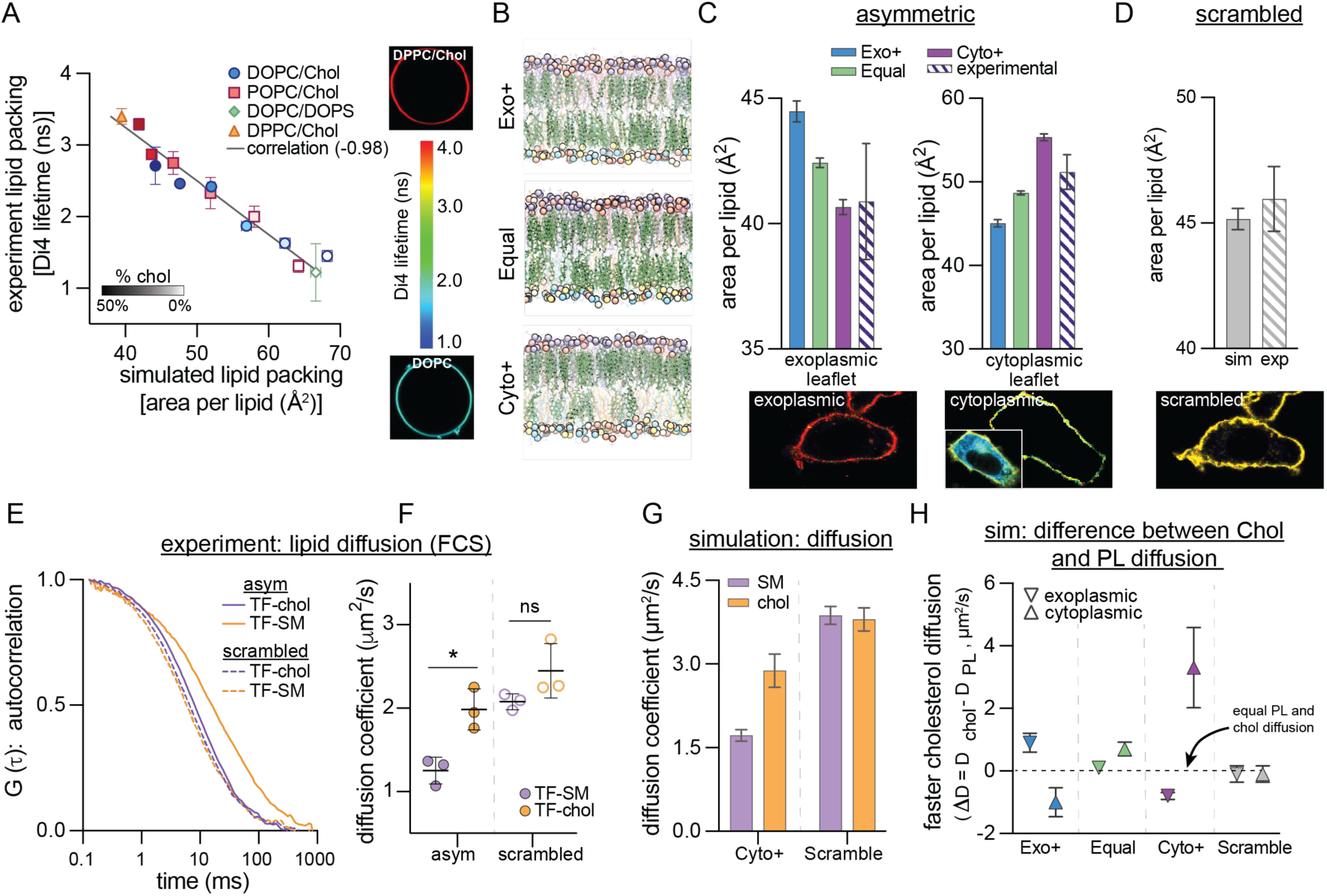
Experimental validation of Cyto+ model. (**A**) Calibration of Di4 lifetime, an experimental reporter of lipid packing measured in giant unilamellar vesicles (GUVs), with simulated area-per-lipid (APL) across a set of synthetic symmetric bilayers. Shown is average ± SD of individual experiments. Representative images of the least and most tightly packed GUVs are shown on the right. Color bar indicates Di4 lifetime. Light-to dark-filled symbols represent increasing mol% of cholesterol. (**B**) Snapshots of three simulated PM models having more PLs in their exoplasmic (top) or cytoplasmic (bottom) leaflets, or similar numbers of PLs in both leaflets (middle). Lipid representation is the same as in Figure 1E. (**C**) Leaflet packing densities (area per lipid) from the three simulated PM models with different interleaflet PL abundances from (B) compared to experimental measurements in PMs of live fibroblasts. (**D**) APL of the scrambled Cyto+ model compared to the experimentally scrambled fibroblast PM (via ionophore treatment). Representative images at bottom of C-D are of Di4 lifetime in the PM of fibroblast cells (color scale for lifetime shown in panel A). ‘Cytoplasmic’ panel shows the masked PM of a microinjected cell with full image as an inset. (**E**) Representative FCS curves for TF-SM and TF-chol in the PM of live fibroblasts before (asym) and after (scrambled) ionophore treatment. (**F**) Corresponding diffusion coefficients of TF-SM and TF-chol calculated from FCS measurements. Data points represent means of three biologically independent replicates, each with >5 cells and >5 measurements per cell; thus, each point is the average of >25 FCS curves. (**G**) Diffusion coefficients of SM and cholesterol (chol) in the simulated Cyto+ membrane (left) and its scrambled counterpart (right). Cholesterol diffusion in Cyto+ represents the asymmetry-weighted average of the slowly diffusing population in the exoplasmic leaflet and the rapidly diffusing population in the cytoplasmic leaflet (Figure S13A). (**H**) Difference between cholesterol and phospholipid diffusion (ΔD) in each leaflet of the simulated asymmetric PM models and the Cyto+ scrambled membrane. *p<0.01. All simulations shown in this figure are all-atom.

Next, we investigated the correspondence in lipid *dynamics* between live cells and our Cyto+ model by comparing lipid diffusion in simulated membranes to measurements in cells (Figure 4E−H). Recently, it was reported that a pool of cellular cholesterol diffuses much faster than PLs in both cultured cells and chordate embryos (Pinkwart et al., 2019), which was notably different from synthetic, cell-derived, and computational model membranes, where the two lipid types diffused similarly. Thus, this rapidly diffusing pool of cholesterol may be a hallmark of the lipid organization of many living cell PMs (Pinkwart et al., 2019). We confirmed these findings in fibroblast PMs by measuring diffusion of fluorescently tagged sphingomyelin (TF-SM (Sezgin et al., 2012)) and cholesterol (TF-Chol (Holtta-Vuori et al., 2008; Wustner et al., 2016)) via fluorescence correlation spectroscopy (FCS) (Figure 4E−F). TF-Chol diffused 1.6-fold faster than TF-SM and this difference was eliminated by scrambling PM lipids. Similar trends were observable in the Cyto+ simulation: overall, cholesterol diffused 1.7-fold faster than SM and this difference was eliminated by lipid scrambling (Figure 4G). Further comparison between cholesterol and PL diffusion in each leaflet revealed that these differences were attributable to the much faster diffusion of cholesterol (relative to PLs) in the cytoplasmic PM leaflet (Figure 4H). In contrast, large differences between cholesterol and PL diffusion were not observed in any of the leaflets of the Exo+ and Equal models (Figure 4H). Consistently, cholesterol has been reported to diffuse faster than phospholipids only in simulations of highly unsaturated, loosely packed membranes with low cholesterol concentration (Javanainen and Martinez-Seara, 2019). Thus, the previously reported unexpected dynamics of cholesterol in mammalian PMs appears to be consistent with the unusual lipid configurations proposed by the Cyto+ model.

### Functional consequences of lipid imbalances

Having validated the compositional, structural, and dynamical features of the Cyto+ PM model against cell measurements, we examined the consequences of this unconventional lipid organization on membrane biophysical properties. The large lipid asymmetries suggest the presence of stress in the membrane, and we first calculated the lateral pressure distribution in each leaflet (Figure 5A). While the bilayer has no net stress, the exoplasmic leaflet is compressed to 8.2 ± 0.8 mN/m while the cytoplasmic leaflet is under tension of the same magnitude. Such balanced compressive/tensile forces in an overall-tensionless membrane are termed ‘differential stress’ (Doktorova and Levental, 2022; Varma and Deserno, 2022). In the Cyto+ model, differential stress arises both from differences in leaflet PL abundances and from cholesterol’s preference for the outer leaflet. Lipid scrambling eliminates lipid asymmetries and alleviates differential stress (Figure S11A).

**Fig. 5.**
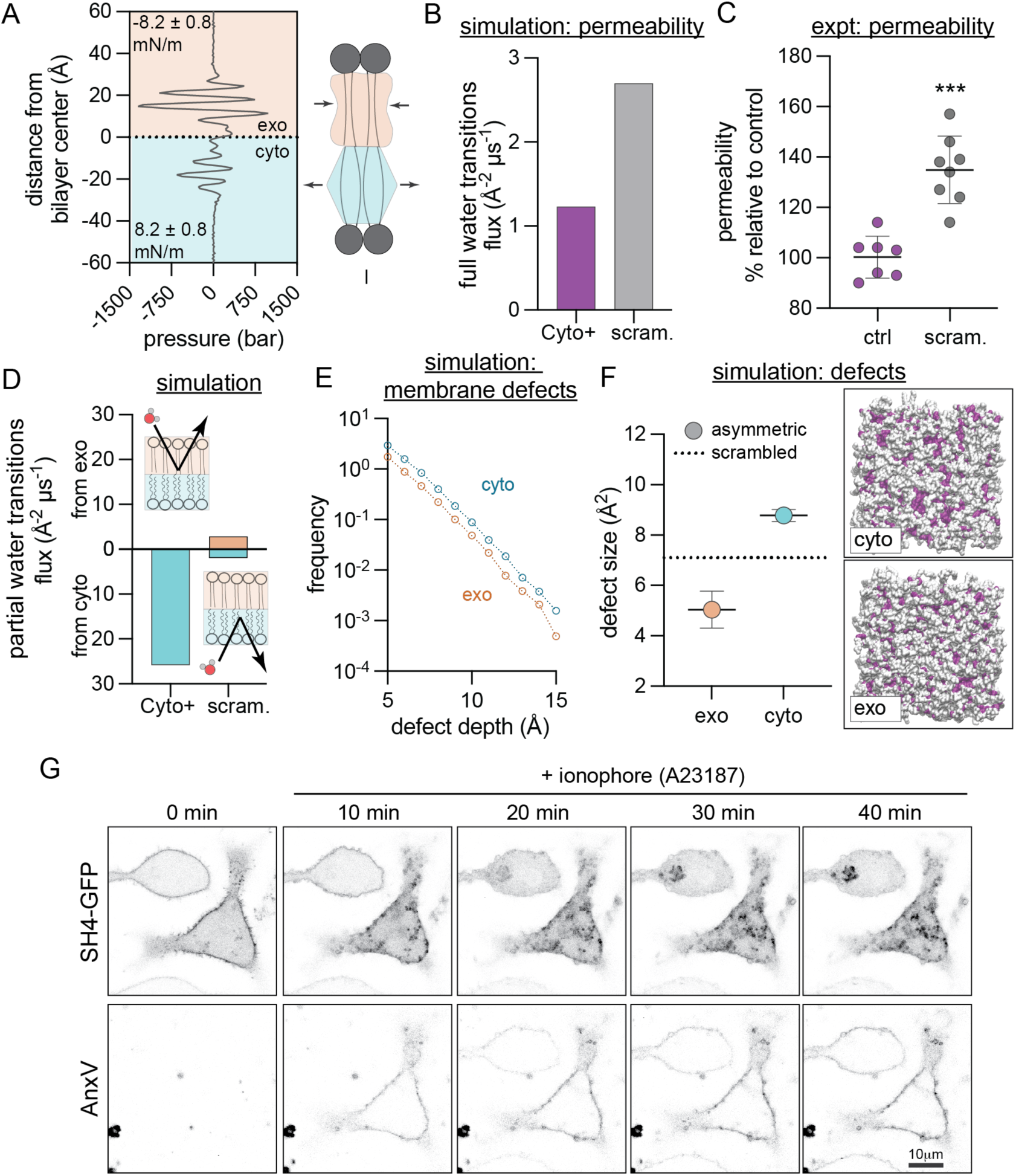
Biophysical features of the Cyto+ model. (**A**) Transbilayer lateral pressure distribution in the simulated Cyto+ PM shows that the bilayer is differentially stressed, i.e. the exoplasmic leaflet (orange) is compressed while the cytoplasmic leaflet (cyan) is under tension of the same magnitude. (**B**) Simulated water flux through Cyto+ compared to scrambled PM. (**C**) Experimental permeability of erythrocyte membranes to FDA. PMA-scrambled erythrocytes shown relative to untreated asymmetric controls. Shown are average ± SD of independent experiments. (**D**) Flux of water molecules partially permeating the leaflets in simulations, i.e. entering the bilayer from the exoplasmic (top) or cytoplasmic (bottom) leaflet and exiting from the same leaflet. (**E–F**) Hydrophobic defects in leaflets of simulated Cyto+ membrane and its scrambled counterpart. (**E**) Histogram comparing defect sizes in the two Cyto+ leaflets. (**F**) Defect size constants of deep defects in the Cyto+ leaflets compared to the scrambled bilayer (dashed line). Illustrative simulation snapshots of the Cyto+ bilayer viewed from the cyto or exo leaflet show shallow (in white) and deep (in color) solvent-exposed areas (defects) in surface representation. Opposite leaflet is shown in gray. (**G**) Redistribution of lipidated peptide, the SH4 domain of Lyn (SH4-GFP), in RBL cells induced by PM scrambling with A23187, evidenced by concomitant exposure of PS monitored by the PS-marker AnxV-647. ***p<0.001

Based on its significant lipid compression, we hypothesized that the cholesterol-enriched outer leaflet may constitute a robust barrier to transbilayer permeability. This hypothesis was tested first *in silico* by calculating the flux of water molecules through the bilayer (i.e. entering from one side and exiting from the other). Comparing the Cyto+ model to its scrambled counterpart revealed that the scrambled PM was ∼80% more permeable to water than the asymmetric PM (Figure 5B), highlighting a functional PM property arising from asymmetric lipid distribution, rather than overall lipid composition. To experimentally validate this observation, we investigated asymmetry-dependent PM permeability in erythrocytes. We measured transbilayer permeation of a small molecule, fluorescein diacetate (FDA), which becomes fluorescent only after encountering intracellular esterases. Because transbilayer flux is the rate-limiting step in this assay, the rate of fluorescence increase is directly proportional to membrane permeability (Levental et al., 2020). Scrambled RBCs were 35% more permeable to FDA than asymmetric RBCs (Figure 5C), in qualitative agreement with the computational observation (Figure 5B). Simulations further suggested that this difference is likely due to the tightly packed outer leaflet, which presents the main barrier to transbilayer permeation in the Cyto+ model (Figure 5D). We observed many partial transitions of water molecules entering and exiting the cytosolic leaflet, but none from the exoplasmic leaflet (Figure 5D), illustrating the bipolar nature of the asymmetric membrane environment.

The low lipid packing, high resting tension, and high permeability of the Cyto+ cytoplasmic leaflet suggest the likelihood of frequent hydrophobic defects, i.e. areas where the hydrophobic bilayer core is transiently exposed to water. Consistent with this prediction, simulations revealed that the Cyto+ cytoplasmic leaflet has more and larger deep hydrophobic defects than either the exoplasmic or scrambled leaflets (Figure 5E−F). Such defects have been previously implicated as “binding sites” for peripheral proteins that interact with membranes through hydrophobic moieties like lipid anchors (Larsen et al., 2015) or amphipathic helices (Cui et al., 2011; Vanni et al., 2013). Our analysis predicts that the structure of the Cyto+ membrane facilitates peripheral protein binding to the cytoplasmic leaflet by promoting the insertion of hydrophobic anchors (Larsen et al., 2015).

To test this hypothesis in live cells, we examined the asymmetry-dependent PM binding of a short lipidated peptide, the SH4 domain of the Src-family kinase Lyn, which associates with membranes via two lipid anchors (Figure 5G). To release PM asymmetry, cells were treated with A23187, a calcium ionophore that induces scrambling (evidenced by binding of the PS-marker Annexin V (AnxV) to the cell surface). In untreated cells, AnxV did not bind, and the peptide localized largely at the PM. Ionophore treatment produced the predicted AnxV binding, but also a concomitant detachment of the peptide from the PM and relocation to the cytosol and intracellular organelles (Figure 5G). The same behavior was observed for two other peptides representative of the membrane anchoring domain of an unrelated peripheral protein, H-Ras (Figure S12). These peptides bear no overall charge, suggesting that their behavior was not dependent on changes in inner leaflet electrostatics (Yeung et al., 2008). Rather, the general desorption of lipidated peptides from the PM after scrambling suggests that their PM association is likely mediated by the deep hydrophobic defects that arise in the tensed, cholesterol-poor cytosolic leaflet.

### Transbilayer lipid imbalance regulates cellular cholesterol sensing

To explore the physiological consequences of asymmetric lipid distributions, we inferred that since most cellular cholesterol sensing and transfer machinery is in the cytoplasm, only cytoplasmic leaflet cholesterol would be relevant to these processes. Therefore, manipulating the determinants of cholesterol’s transbilayer distribution would be expected to affect cellular cholesterol handling. To test this hypothesis, we manipulated two of these determinants: outer leaflet sphingomyelin (Fig 6) and transbilayer phospholipid abundance (Figure 7).

**Fig. 6.**
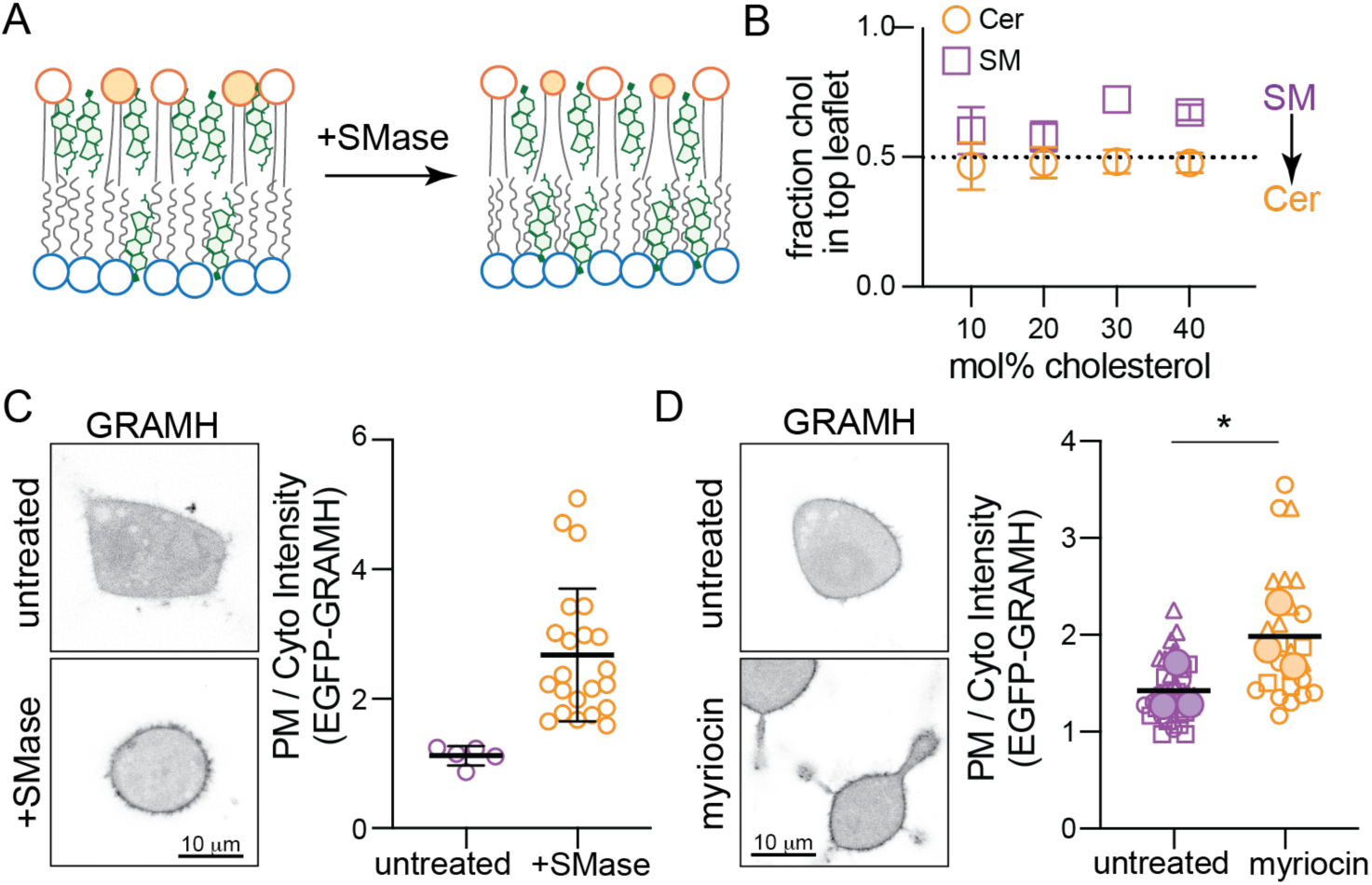
Sphingomyelin drives cholesterol asymmetry. (A) Schematic of proposed Chol redistribution induced by SMase. (B) Interleaflet Chol distributions in CG simulations of the simplified CG PM model from Fig. 2A before (purple) and after (orange) all SM has been converted to ceramide. (C) EGFP-GRAMH localization in RBL cells following SMase treatment; representative images on left, quantification on right. Each data point is a cell. (D) EGFP-GRAMH localization in RBL cells treated with 25 μM myriocin for 24 h; representative images on left, quantification on right. Small symbols represent individual cells, with symbol shapes denoting independent experiments. Filled larger symbols are means of independent experiments. Paired t-test on means of independent experiments; *p< 0.05.

**Fig. 7.**
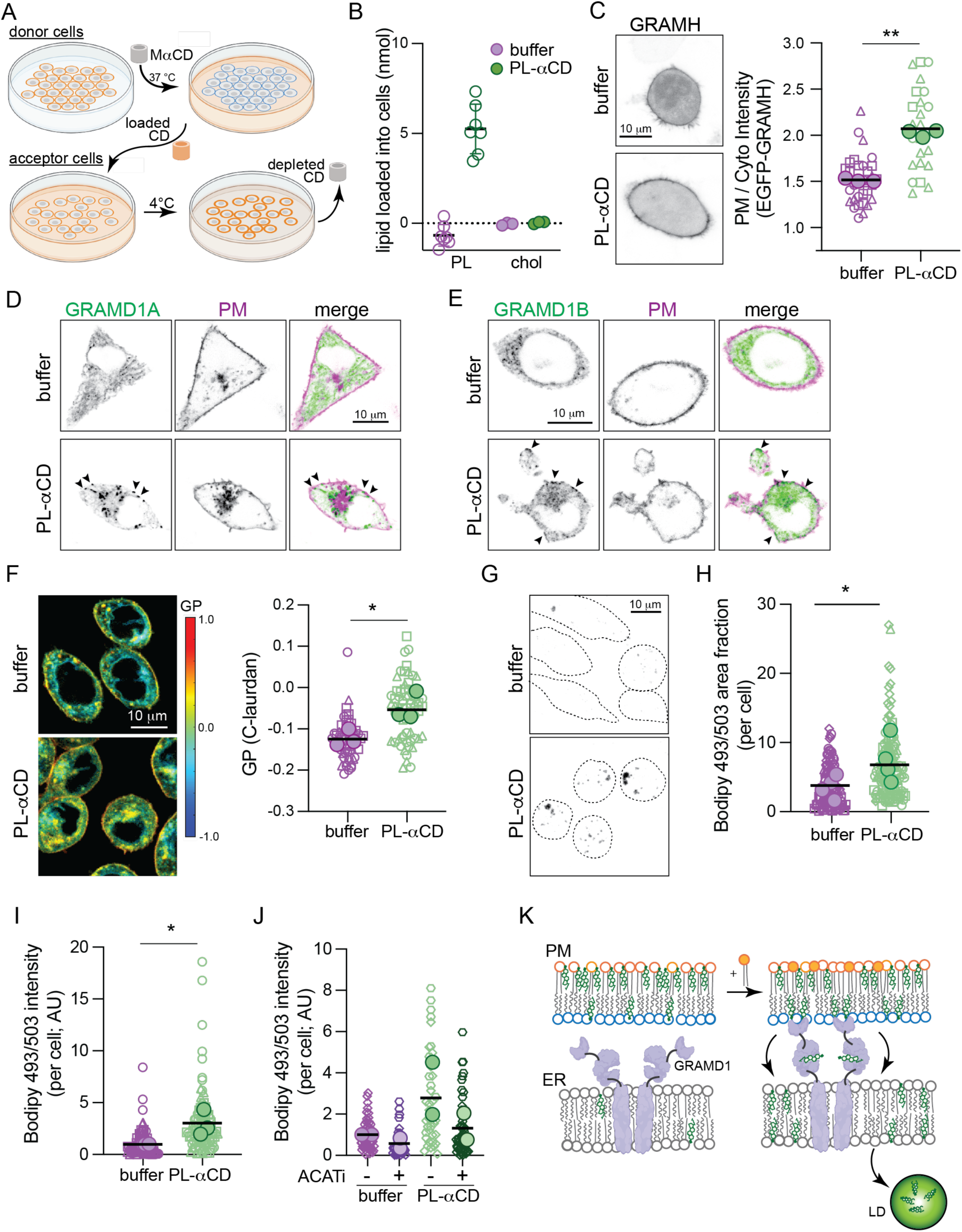
Cholesterol homeostasis is regulated by PM transbilayer phospholipid distribution. (**A**) Schematic of PL loading into the PM outer leaflet of target cells via MαCD extraction of PL and loading onto fresh cells. (**B**) Quantification of PL and Chol loading into ∼3 x 10^5^ RBLs via PL-MαCD. (**C**) EGFP-GRAMH localization in cells treated with PL-MαCD; representative images on left, quantification on right. (**D**) EGFP-GRAMD1A and (**E**) EGFP-GRAMD1B localization in cells treated with PL-MαCD (co-expressing a PM marker, magenta). Arrowheads denote GRAM protein puncta at the PM induced by PL-MαCD. (**F**) Left, C-laurdan GP maps of cells treated with PL-MαCD. Warm colors represent higher GP (i.e. tighter lipid packing). Right, quantification of GP of internal membranes. (**G**) Bodipy 493/503 staining of cells treated with PL-MαCD. Representative confocal max projection images. (**H**) Quantification of area fraction with Bodipy staining per cell. (**I**) Quantification of total Bodipy intensity per cell. (**J**) Quantification of Bodipy 493/503 intensity in cells treated with PL-MαCD with and without ACAT inhibition (1 ug/mL Sandoz 58-035). (**K**) Schematic of cholesterol translocation from the outer leafler of the PM to lipid droplets when phospholipids are loaded to the outer leaflet. In C, F, H, I, and J small symbols represent individual cells, with symbol shapes denoting independent experiments. Filled larger symbols are means of independent experiments. Paired t-test on means of independent experiments; p< 0.05, p<0.01.

Chol accumulates in the exoplasmic PM leaflet in part due to its preferred interactions with saturated sphingolipids. Thus, we reasoned that reducing SM levels would lower cholesterol’s affinity for the outer leaflet, leading to its redistribution to the inner leaflet (Figure 6A). Simulations supported this inference, as replacing SM with Ceramide (Cer) in simulated asymmetric membranes (see Fig 2A) notably reduced Chol asymmetry and increased its concentration in the cytoplasmic leaflet (Figure 6B). To explore this effect experimentally in living cells, we used a cytoplasmic Chol-binding probe based on the cholesterol-sensing element of the cholesterol transport protein GRAMD1b (Aster-B). The isolated GRAM domain of this protein has been engineered into a biosensor (GRAM-H), which translocates to the PM when ‘accessible cholesterol’ is increased (Koh et al., 2023). We observed a robust and significant increase in PM recruitment induced by treating cells with SMase to convert SM into Cer (Figure 6C) or by inhibiting SM synthesis with myriocin (Figure 6D). Thus, reducing outer leaflet SM increased inner leaflet cholesterol. These results are quantitatively consistent with previous observations of SMase treatment (Koh et al., 2023) and provide a mechanistic explanation: cholesterol ‘accessibility’ is determined, at least in part, by its transbilayer asymmetry.

To manipulate transbilayer phospholipid imbalance, we developed a method to increase the phospholipid abundance of the PM outer leaflet without changing its composition (Figure 7A). We extracted outer leaflet PLs from a confluent dish of ‘donor’ RBL cells at 37°C with MαCD. We then transferred the PL-saturated-MαCD to a new dish of ‘acceptor’ RBL cells, which had lower confluence and incubated at 4°C (Figure 7A). We reasoned that lower temperature reduces the solubility of PL in MαCD and leads to its deposition into the ‘acceptor’ cells, as previously shown in synthetic systems (Reagle et al., 2024). To test this hypothesis, we measured the lipid abundance in the PL-MαCD solution before and after incubation with acceptor cells and found that 4 nmol of PLs could be delivered to ∼3 x 10^5^ cells (Figure 7B), increasing outer leaflet lipid abundance in acceptor cells by ∼40% (see SI Sec 5.5 for details). This manipulation did not induce PM lipid scrambling (no binding of PS sensors to cell surface, Fig S18B) nor extensive PM deformations (Bettache et al., 2003).

According to our hypothesis, increasing PL abundance in the outer leaflet should redistribute cholesterol to the inner leaflet (compare Cyto+ versus Exo+ in Fig 4B), and indeed we observed significant recruitment of GRAM-H to the PM upon outer leaflet lipid loading (Figure 7C). Similar effects were observable when the PM was loaded with pure synthetic lipids (Figure S18D-E). We then hypothesized that the increase in inner leaflet cholesterol would activate cellular cholesterol homeostasis machinery, leading to cholesterol transport to internal organelles. The machinery mediating such PM-to-ER cholesterol transport has been recently identified as the GRAMD1/Aster proteins, which are ER residents that translocate to PM-ER contact sites upon cholesterol loading and use their STAR domains to shuttle excess PM cholesterol to the ER (Besprozvannaya et al., 2018; Naito et al., 2019; Sandhu et al., 2018). We observed translocation of GRAMD1A (Figure 7D) and GRAMD1B (Figure 7E) to puncta at the PM induced by loading either outer leaflet or synthetic PLs (Figure S18F) into the PM outer leaflet, presumably due to cholesterol redistribution to the inner leaflet. Consistent with our inference that GRAMD1b transports excess cytosolic leaflet cholesterol from the PM to the ER, the membrane packing sensor C-laurdan reported a significant increase in lipid packing of internal cell membranes induced by loading the outer leaflet with PLs (Figure 7F).

During Laurdan imaging, we noted spherical structures with unusually high GP values (Figure 7F, Figure S18C), which have been previously associated with cholesterol ester-rich lipid droplets (LDs) (Ventura et al., 2023). We hypothesized that these may represent newly accumulating LDs induced by the transfer of cholesterol to the ER, which activates cellular cholesterol storage via its esterification and deposition into lipid droplets (Chang et al., 2006). To assay LD accumulation, we stained cells with the neutral lipid dye Bodipy 493/503, which accumulates in LDs, and imaged cell volumes by confocal microscopy (Figure 7G). We observed a robust increase in the number, size, and fluorescence intensity of LDs induced by PL loading (Figure 7G-I, S18G-H), similar to loading cells directly with cholesterol (Figure S18K-L). We observed the same effect after SMase treatment (Figure S18I-J), consistent with SMase inducing redistribution of cholesterol to the cytoplasmic leaflet (Figure 6). To confirm that these LDs were due to increased cholesterol esterification, we inhibited the enzyme responsible for cholesterol esterification, acyl-coenzymeA:cholesterol acyltransferase (ACAT). Indeed, ACAT inhibition blocked accumulation of LDs induced by outer leaflet PL loading (and SMase), confirming their dependence on cholesterol esters (Figure 7J, S18G-L).

These observations reveal that cellular cholesterol sensing, homeostasis, and storage can be activated without changes in cholesterol levels (Fig S18A), but rather by its transbilayer distribution in the PM which can be regulated by the PL composition and abundance of PM leaflets.

## Discussion and implications

We report a previously overlooked feature of cell PMs: large differences in steady-state phospholipid abundance between bilayer leaflets, facilitated by asymmetric cholesterol distribution. High cholesterol concentrations support bilayer tolerance for the large PL imbalances revealed by our lipidomics measurements (Figures 1-2), and cholesterol extraction compromises membrane integrity (Figure 2C–E). While the relative paucity of PLs in exoplasmic leaflets was unexpected and had not been previously reported, re-analysis of previous measurements of lipid asymmetry confirms our observations (Table S1, Figure 1D). This result was perceived as physically unrealistic due to the usually implicit assumption that leaflets with equal areas must also contain similar PL abundances (Rawyler et al., 1985). This assumption is fundamental to most current conceptions of biomembrane lipid distributions, underlying the interpretations and design of most experiments and models. However, theory (Varma and Deserno, 2022), simulations, and our measurements reveal that this assumption is generally invalid and is incorrect for mammalian PMs.

Cholesterol sustains interleaflet PL imbalances by accumulating in PL-poor leaflets. This capability is enabled by cholesterol’s uniquely rapid passive diffusion between leaflets. Cholesterol’s equilibrium distribution is determined by a balance of its tendency to “fill gaps” in the underpopulated leaflet, its preference for saturated and sphingo-lipids, and entropic effects favoring symmetric arrangement (Figure 3A–B). The balance between these inputs is described in an elegant theory by Deserno and coworkers (Foley et al., 2023; Varma and Deserno, 2022). As exoplasmic leaflets in our measurements are not only more saturated but also relatively PL-poor, cholesterol is predicted to enrich in the exoplasmic leaflet. Consistent with this reasoning and theoretical predictions (Varma and Deserno, 2022), we observe enrichment of cholesterol in the erythrocyte outer leaflet (Figure 3D). This result is consistent with a study that used engineered sensors to infer much higher cholesterol concentrations in the outer PM leaflet (Liu et al., 2017). Notably, in two recently reported structures of membrane proteins determined in native cell membranes (Coupland et al., 2024; Tao et al., 2023), cholesterol was associated to the exoplasmic side of the transmembrane domains, consistent with its enrichment in the exoplasmic leaflet. The possibility of significant cholesterol enrichment in one leaflet was questioned on the theoretical grounds that rapid cholesterol flip-flop would appear to preclude significant cholesterol disparity between leaflets (Steck and Lange, 2018). Our findings reveal a simple mechanism that not only allows, but demands, interleaflet cholesterol imbalances. While other studies have reported on cholesterol’s distribution in PMs (Figure S8 caption), our model provides a mechanistic explanation for cholesterol’s ultimate transbilayer distribution.

Experimental measurements of leaflet lipid packing support our model of a PM with a PL-rich and cholesterol-poor cytoplasmic leaflet (Figures 4A−D, S10). Similarly, experimentally observed fast cholesterol diffusion (Figure 4E−F) appears to be a signature of a differentially stressed, unsaturated lipid-rich, cholesterol-poor leaflet (Figures 4G−H, S13), consistent only with the Cyto+ model. The inference that the cytoplasmic PM leaflet is under tension may seem contradictory with its relative over-abundance of PLs (Figure 1). However, in the presence of cholesterol, such stresses are not solely the result of lipid imbalances, but rather several independent drivers of cholesterol distribution (Figure 3), which can induce, rather than alleviate, leaflet tension (Figure S6). It may be surprising that a membrane with one compressed and one tensed leaflet would not spontaneously deform (i.e. bend) to alleviate these stresses. However, differential stress can be balanced by curvature stresses arising from differences in the preferred (spontaneous) curvatures of leaflet lipids (Hossein and Deserno, 2020), as is the case for the Cyto+ PM model (Figure S11B). Further, other mechanical constrains (most prominently the cytoskeleton) may prevent spontaneous bending in living cells.

The asymmetric distributions of PLs and cholesterol in the two PM leaflets revealed by our analyses have far-reaching implications for biomembrane structure and function (Figures 5-7). Lipid abundance asymmetries between the two PM leaflets provides cells with novel regulatory modes, namely the possibility of tuning leaflet packing, tension, and dynamics independent of lipid chemistry. These arrangements may underlie PM properties such as low permeability and regulated binding of lipidated proteins and be relevant for cellular responses to membrane lipid perturbations, e.g. in genetic diseases of (sphingo)lipid metabolism (Pekkinen et al., 2019; Simons and Gruenberg, 2000) or from dietary lipids (Levental et al., 2020). Crucially, abundance asymmetry also regulates the mechanisms of cellular cholesterol homeostasis (e.g. trafficking, efflux, storage), which underlie sterol uptake (Ferrari et al., 2023), bile acid production (Cortes and Eckel, 2022), steroid hormone synthesis (Sandhu et al., 2018), and the levels of circulating lipoproteins that are strongly predictive of cardiovascular disease (Xiao et al., 2023). We therefore posit that abundance asymmetry of membrane lipids presents a previously unexplored, yet broadly consequential dimension of PM organization and function.

## Limitations of the Study

The most significant limitation of our study is that direct analysis of lipidomic asymmetry remains limited to erythrocyte plasma membranes. While the general features of this asymmetry (e.g. charged lipids confined to inner leaflet) have been confirmed in many contexts (Doktorova et al., 2020b), neither the detailed lipid compositions of PM leaflets nor their abundance imbalances can be directly measured in nucleated cells using currently available methods. The same limitation persists for organellar membranes. Similarly, directly measuring cholesterol asymmetry remains a major technical challenge: our experiments reveal relative enrichments but not absolute concentrations. In the context of these limitations, it is important to emphasize that both parameters (leaflet composition and interleaflet abundance imbalance) are likely context dependent, being determined by a complex interplay between cell state (e.g. lipid metabolism, membrane trafficking) and its environment (e.g. raw materials for lipid synthesis). Thus, there is unlikely to be a single, universal description of living membrane asymmetry; rather, our study emphasizes that phospholipid and cholesterol abundance imbalances are an independent, regulated, physiologically important aspect of the membrane phenotype (Bigay and Antonny, 2012). Finally, our study focuses solely on computational, synthetic, and cultured cell models, thus its implications for physiological lipid homeostasis remain hypothetical. While the epidemiological connections between diet, health, and disease are among the strongest correlations in biomedicine, underlying mechanistic understanding remains incomplete. Whether and how transbilayer lipid asymmetries affect these mechanisms of physiological lipid handling is an important future direction of research.

## Acknowledgments

We thank Madhusmita Tripathy and Anand Srivastava for providing their code for calculating packing defects in three dimensions, and Anne Kenworthy, Robert Ernst, and Itay Budin for critical reading of the manuscript. M.D. was supported by NIH F32GM134704 and SciLifeLab & Wallenberg Data Driven Life Science Program grant: KAW 2024.0159. J.S. is funded by the Marie Skłodowska-Curie Actions Postdoctoral Fellowship (Grant 101059180). E.S. is funded by Karolinska Institutet, SciLifeLab, Swedish Research Council Starting Grant 2020-02682 and Human Frontier Science Program (Grant RGP0025/2022). F.A.H. is supported by NIH/NIGMS R01GM138887. E.L. was supported by R01 GM120351. J.L.S was supported by GM008280. K.R.L. and I.L. are supported by NIH (GM120351, GM134949), the Volkswagen Foundation (grant 93091), and the Human Frontiers Science Program (RGP0059/2019).

## Author contributions

Conceptualization: MD, KRL, IL

Methodology: MD, JLS, KRL, JHL, EL, FAH, ES

Investigation: MD, JLS, XZ, HW, JS, KRL

Visualization: MD, KRL, IL, JLS

Formal analysis: MD, KRL, JLS, JS

Resources: EL, ES

Data curation: MD, KRL

Funding acquisition: MD, JS, ES, FAH, EL, IL

Supervision: ES, EL, FAH, KRL, IL

Writing – original draft: MD, KRL, IL

Writing – review & editing: MD, JLS, XZ, HW, JS, JHL, FAH, ES, EL, KRL, IL

## Data and materials availability

All data and code used for analysis will be uploaded to a publicly accessible archive.

## Methods

### Lipidomics

Human erythrocytes (freshly isolated and intact) were treated with two enzymes, phospholipase A2 (PLA2, Sigma) and sphingomyelinase (SMase, Sigma) to digest exclusively the lipid species on the exoplasmic PM leaflet as detailed in ^1^ and summarized in SI. Enzyme-treated cells were analyzed with shotgun electron spray ionization with tandem MS-MS (ESI-MS/MS) by Lipotype and compared to untreated controls to determine the abundance of ∼400 unique phospholipid species, including glycosphingolipids, on the exoplasmic leaflet via the extent of digestion. MS data acquisition and analysis were performed as in ^1^ and summarized in Supplemental Information.

### Phospholipid extraction from synthetic liposomes

Large unilamellar vesicles (LUVs) composed of POPC (Avanti) with 0, 20 and 40% Chol (Avanti) were prepared at a concentration of 24 mM as described in ^2^. Hydroxypropyl-alpha-cyclodextrin (HP*α*CD, Sigma Aldrich) was dissolved in water at a nominal concentration of 500 mM and spun through a pre-washed centricon filter device (Ultra-15, 100,000 Da molecular weight cutoff; Amicon; MilliporeSigma) to remove any aggregates as detailed in SI. POPC extraction from the liposomes was achieved by mixing 170 *μ*M LUVs with increasing amounts of HP*α*CD at room temperature. Light scattering was measured on an Anton Paar Litesizer 100 at 22°C. Subsequently, each sample was spun through the same Centricon filter device as above; the filtrate was transferred to a glass culture tube and spiked with 2 *μ*l of DMPC (Avanti, 25.27 mg/mL in chloroform); the sample was frozen overnight and lyophilized the next day. The lyophilized powder was subjected to acid-catalyzed methanolysis to extract the fatty acid methyl esters (FAMEs) and their analysis was done on an Agilent 5890A gas chromatograph (Santa Clara, CA) with a 5975C mass-sensitive detector operating in electron-impact mode and an HP-5MS capillary column (30 m x 0.25 mm, 0.25-mm film thickness) as described in ^2^ and summarized in SI. The respective peak areas of the FAMEs corresponding to the myristoyl chain of DMPC and the palmitoyl chain of POPC were recorded and used to calculate the normalized amount of POPC in the sample.

### Cell culture

RBL-2H3 cells were purchased from ATCC and grown in Eagle’s Minimum Essential Medium (EMEM) media (Gibco) with 10% fetal bovine serum (FBS, Genessee) and 1% penicillin-streptomycin (Gibco). 3T3 cells were grown in Dulbecco’s Modified Eagle Medium (DMEM) supplemented with 10% FBS.

### Cholesterol extraction from cells

RBL-2H3 cells were washed twice with CaR buffer (145 mM NaCl, 5 mM KCl, 10 mM HEPES at pH 7.4), then incubated with CaR buffer containing either 2 mM CaCl_2_ or 2 mM EDTA in the absence or presence of 6 mM M*β*CD (Thermo Fisher). For monitoring PS exposure and dextran permeability, the buffer also contained LactC2-mClover and Rhodamine-dextran (Sigma-Aldrich) as detailed in SI. For PI permeability measurements, the buffer contained propidium iodide (Alfa Aesar). Cells and buffer were incubated at room temperature (20−22°C) and imaged at the indicated time points (30, 60, and 90 minutes). Confocal fluorescence imaging of cells was performed at 21°C using a Nikon C2+ point scanning system attached to a Nikon Eclipse Ti2-E microscope equipped with a Plan Apo Lambda 60X oil immersion objective and a Bioptech objective cooling collar. Imaging data was analyzed with FIJI as described in SI.

### DHE distribution in erythrocytes

Human erythrocytes were isolated from healthy donors with informed consent and suspended in Ringer’s solution (125 nM NaCl; 5 mM KCl; 3 mM CaCl2; 1 mM MgSO4; 30 mM HEPES; 5 mM glucose; pH 7.4). DHE (Sigma Aldrich) was dissolved in M*β*CD (160 mM in Ringer’s solution) at 1:8 ratio at room temperature and centrifuged briefly to remove any aggregates. 3.5×10^8^ cells/mL were suspended in a mixture of the DHE/M*β*CD solution and free M*β*CD in Ringer’s solution for final concentrations of 50 *μ*M DHE and 1.2 mM M*β*CD, unless otherwise specified, and nutated for 10 minutes at room temperature. The sample was washed 3 times with Ringer’s solution with 2,000 rcf spins for 3 minutes at room temperature. Di4 (Di-4-ANEPPDHQ, Thermo Fisher, dissolved in ethanol) was diluted 10-fold in Ringer’s solution to 100 µg/mL and added to cells (1.5×107 cells/mL) at room temperature. Di4 addition was directly followed by measurement of DHE emission spectrum on a PTI fluorimeter at 37 °C. To minimize inner filter effect (IFE), measurements were taken using a 10 × 2 mm pathlength cuvette oriented with 2 mm pathlength facing detector and minimal sample volume. Excitation and emission slits were open to 5 nm, the gain was optimized for increased signal, and measurements were taken at a scan rate of 0.1 s/nm to minimize photobleaching. DHE quenching was monitored by measuring the decrease in emission at 385 nm upon addition of Di4 until a plateau was reached. The relative amount of quenched DHE fluorescence at this plateau was used to quantify the fraction of DHE (and therefore, cholesterol) residing in the outer leaflet of the membrane.

### Quantification of sterol content in erythrocytes

To quantify total sterol content in the erythrocyte membrane, lipids were first extracted using a two-step extraction protocol ^3^. Amplex Red cholesterol assay was performed (according to manufacturer instructions; Invitrogen) to determine the abundance of cholesterol and DHE. Each reading was normalized to cell count. Technical triplicates were measured for each sample. All values are normalized to untreated cells and are the average and standard deviations from at least two independent biological replicates.

### DHE distribution in synthetic liposomes

Symmetric large unilamellar vesicles (LUVs) at 25 mg/mL lipid were prepared as in ^2^. Unsaturated lipids were extruded at room temperature, and multicomponent mixtures were extruded at 45°C. For DHE-containing vesicles, DHE was incorporated into the lipid mixture in organic solvent (prior to drying and hydration). DHE titrations were performed by replacing cholesterol for DHE and keeping total sterol concentration constant. Measurements were done on a PTI fluorimeter as described above. 1 *μ*L of 2.5 mg/mL LUVs was diluted into 165 *μ*L of Ringer’s solution, yielding ∼15 μg/mL LUVs in the cuvette. If Di4 was not already incorporated into the liposomes, Di4 (stock dissolved in ethanol) was first diluted 10x in Ringer’s solution, then 1.5 μL from the diluted solution were added to the LUV suspension and pipette mixed. Samples were checked for risk of photobleaching via time-based emission scans. DHE emission spectra was obtained by excitation at 310 nm and emission between 340−500 nm at 37 °C. Measurements with externally added Di4 were taken immediately after addition with each sample of LUVs discarded after each measurement and the cuvette thoroughly washed.

### DHE self-quenching in liposomes

Vesicles were prepared following the rapid solvent exchange (RSE) protocol as described previously ^4^ and summarized in the SI. The fluorescence intensity was measured on a Horiba Fluorolog 3 spectrofluorometer by exciting the sample at 327.8 nm and collecting the emission in the range 340−640 nm. The excitation and emission slits were set to 2.5 and 3 nm, respectively and the integration time was 0.1 s. The 2 mL sample was continuously stirred during the measurement via a flea stir bar placed in the cuvette.

To compare the DHE fluorescence across samples, the full spectrum of each sample was first normalized by subtracting the spectrum of the respective lipid-only (without DHE) sample. The normalized spectra were then integrated to obtain total DHE fluorescence, and the value was corrected for the volume difference measured during sample preparation. Integration of the DHE spectra, as opposed to using the fluorescence at a fixed wavelength, was necessary due to changes in the shape of the spectrum as a result of the formation of various DHE configurational states at higher DHE concentrations ^5^ (Figure S15).

### Scrambling of erythrocyte membranes

To scramble erythrocyte membranes, live cells (∼3.4×10^8^ cells/mL) were incubated with 100 *μ*M phorbol-12-myristate acetate (PMA, Cayman Chemicals) in Ringer’s solution for 10 minutes at room temperature. PMA was used to scramble instead of A23187 because the latter is fluorescent and therefore interferes with DHE and Di4 measurements.

### Permeability measurements in erythrocytes

2×10^6^ cells/mL were resuspended in Ringer’s solution, 1 μg/mL fluorescein diacetate (Cayman Chemicals, 1 mg/mL in acetone) was added and mixed by pipetting, and the fluorescence intensity was read at 450 nm excitation and 510 nm emission over a period of 1 min (at 1 Hz) with 5 nm excitation and emission slits. In each sample, fluorescence increased linearly, thus the slope of fluorescence over time remained constant over the 60 s, consistent with a constant flux (Q) (Figure S16). The permeability coefficient (*P*) is defined by Fick’s Law as *Q* = *PA*(C_out_ − C_in_) where *P* is permeability, *A* is the area of the membrane, and C_out_ and C_in_ are the concentrations of the solute outside and inside the cells, respectively. In our experiments, C_in_ = 0 because FDA is quickly converted into fluorescein upon entering the cell. With equal *A* (due to same number of cells and cellular membranes) and C_out_ across samples, permeability can be directly compared between conditions by normalizing to control samples.

### Peptide binding to cell plasma membranes

RBL cells at 70-80% confluency were harvested with Trypsin (Gibco) and transfected with the respective plasmids via electroporation (see SI for details). After 24 hours, cells were washed with PBS buffer, then maintained in Annexin V buffer (150 mM NaCl, 10 mM HEPES, 2 mM CaCl2, pH 7.4, with 100x diluted Annexin V AF647 from ThermoFisher, #A23204). Ionophore A23187 (Cayman, #11016) diluted in Annexin V buffer was added to the cells with a final concentration of 6 µM. Cells were imaged immediately after the addition of A23187 with a 63X water immersion objective on a Leica SP8 confocal microscope. Images were taken with 10 second intervals for up to 40 minutes.

### Measurement of cholesterol and lipid diffusion in nucleated cells

3T3 cells were seeded on glass-bottom dishes two days before the diffusion experiments. For fluorescent labeling, the fluorescent lipid analogs (Topfluor SM and Topfluor Chol, Avanti Polar Lipids) were first dissolved in DMSO at a final concentration of 1 mg/ml. Before labelling, the cells were washed twice with L-15 medium to remove the full media. Later the fluorescent analogs were mixed with L-15 medium at a 1:1000 ratio (final concentration of 1 µg/ml). The cells were incubated with this suspension for three to five minutes at room temperature, then washed with L-15 twice followed by fluorescence correlation spectroscopy (FCS) measurements. To measure diffusion of scrambled plasma membranes, 3T3 cells were treated with 3 µM A23187 ionophore for eight to ten minutes at 37°C before lipid labelling. Scrambling was confirmed by AnnexinV-AF647 labelling (ThermoFisher A23204). FCS was carried out using a Zeiss LSM 780 microscope with a 40X water immersion objective (numerical aperture 1.2) as described previously ^6^.

### Measurement of Di4 lifetime in GUVs and cells

Giant unilamellar vesicles (GUVs) were prepared by electroformation, and Di4 was added to them externally as described previously ^1^. The samples were imaged at room temperature on a Leica SP8 FLIM confocal microscope with excitation at 485 nm with an 80 MHz pulsed laser, and the lifetime collected in an emission window of 550−800nm. The data was fit using a 2-component reconvolution fit, and the intensity-weighted average lifetime was reported.

To measure Di4 lifetime in the outer leaflet of 3T3 cells, cells were incubated with 1 μg/mL Di4 at 4 °C for 8 min in CaR buffer (145 mM NaCl, 5 mM KCl, 10 mM HEPES) supplemented with 2 mM CaCl_2_ and 2 mM MgCl_2_ at pH 8.0. Cells were briefly washed twice in the same buffer at ambient temperature before imaging at room temperature. To prevent contaminating signal from internalization of the dye, all images were acquired within 20 minutes. To measure Di4 lifetime in scrambled plasma membranes, 3 mM A23187 was added to cells in the same CaR buffer for 15 minutes at 37 °C. Following this treatment, the cells were washed with CaR buffer, with Di4 incubation and imaging as above.

### Extraction and loading of plasma membrane phospholipids

Phospholipids were extracted from a pool of ‘donor’ RBL cells by incubating the cells with Hexakis-(2,3,6-tri-O-methyl)-alpha-Cyclodextrin, MaCD (Cyclodextrin-Shop) dissolved in CaR buffer, at 37°C for 5 min. The supernatant was removed and added to a fresh batch of donor RBL cells for a second incubation. Following that, the supernatant containing MaCD in complex with native PM lipids was filtered to discard any cell debris. The resulting PL-aCD stock solution was incubated with a new batch of RBL cells at 4°C for 15 min to load the extracted phospholipids into the cells PMs. The amounts of phospholipids in the PL-aCD solution before and after the 4°C incubation were quantified with a colorimetric inorganic phosphate assay, and the amount of lipid loaded into the PM was calculated from their difference.

### Quantification of lipid droplets

After manipulation of PM lipid composition, cells were incubated in Tyrode’s buffer for 45 min at 37 °C. Following that 2 *μ*M Bodipy 493/503 (Thermo Fisher) in Tyrode’s buffer was added to the cells and incubated at 37 °C for another 15 min. The cells were then washed and imaged at room temperature. For every cell, Z-stacks of images were taken every 1 *μ*m to capture the entire cell volume, then converted to a maximum projection. ROIs were drawn around the cell perimeter, background was subtracted and the total pixel intensity and percent area covered by non-zero pixels were calculated. 15-25 cells were imaged for each condition per experiment.

### Molecular Dynamics simulations

All bilayer simulations were initially constructed and equilibrated with CHARMM-GUI ^7^. Coarse-grained (CG) simulations were carried out in Gromacs ^8^ with the Martini 2.2 force-field ^9^ and all-atom simulations were run with NAMD ^10^ and the CHARMM36 force-field ^11^ unless otherwise noted. All simulation parameters and solvation details are described in the SI.

#### Tolerance for PL imbalance, CG simulations

A series of asymmetric membranes were constructed with a lipid composition mimicking that of the erythrocyte plasma membrane but with fixed PL imbalance (35% more PLs in the cytoplasmic leaflet) and varying amounts of Chol (ranging between 10 and 50 mol% of all membrane lipids). Table S2 summarizes the initial lipid make-up of the membranes.

After equilibration all cholesterol molecules were replaced with a recently optimized cholesterol model ^12^ which circumvents a previously reported problem with cholesterol in Martini ^13^, and then simulated for 10 *μ*s. Cholesterol interleaflet distribution was determined from the orientation of the Chol molecules. Phospholipids did not change their leaflet residence during the simulations. The maximal phosphate-to-phosphate distance was calculated from the difference between the minimum and maximum z positions of all phosphate (PO4) beads in each frame of the trajectory.

#### Effects of PS flip-flop on tension, AA simulations

To evaluate the effect of flipping PS lipids to the opposite leaflet, we performed a series of AA simulations in which the total number of lipids in the bilayer remained fixed and we mimicked PS flipping by generating different starting configurations with respect to PS interleaflet distribution. The effects of Chol flipping were examined in the same way. The lipid compositions, system sizes and total trajectory lengths of the simulations are listed in Table S3. The tension in each leaflet was calculated as described previously ^14^ and summarized in the SI. Errors on the tension were estimated with a bootstrapping approach in which a dataset with the same total number of frames was reconstructed by picking frames from the original dataset at random and allowing repetitions. The pressure profile and leaflet tensions were then obtained from the resampled data. This procedure was repeated multiple times to obtain the error (i.e. standard deviation) of the leaflet tension.

#### Cholesterol distribution in simpler asymmetric membranes

To examine the effects of phospholipid imbalance and tension on cholesterol distribution in asymmetric membranes, we performed a series of simulations first CG and then in AA representations.

Initially, 8 asymmetric bilayers with one saturated leaflet (composed of 100 DPPC lipids) and one unsaturated leaflet (composed of a varying number of DAPC lipids), each containing 30 mol% cholesterol (Chol), were constructed and simulated (Table S4). To count the number of lipids and Chol in each leaflet, the bilayer was first centered so that the bilayer midplane was in the center of the simulation box in every frame. Then, the DAPC and DPPC lipids whose phosphate (PO4) beads were below/above the bilayer center were counted as belonging to the bottom/top leaflets respectively. Chol was treated the same way except that its headgroup OH bead was used instead. Chol distribution equilibrated within the first few *μ*s in all systems and all reported quantities are averaged over the last 5 *μ*s of the trajectories.

The equilibrated lipid distributions from the CG simulations were used to construct and simulate corresponding all-atom bilayers (Table S4) for detailed analysis of leaflet tension.

#### PM models: AA simulations

To examine the effects of PL imbalance on cholesterol interleaflet distribution and bilayer properties we built 3 asymmetric models of the plasma membrane. Each had ∼40 mol% overall cholesterol and the same leaflet phospholipid compositions modeled after the lipidomics results, but differed in the relative total abundances of PLs in their two leaflets (see Table S5 for the exact numbers and types of lipids in each leaflet). The bilayers were constructed and simulated as described in the SI.

Prior to analysis each system was centered so that the bilayer was in the middle of the simulation box and the average (x,y,z) of the bilayer midplane was at (0,0,0). The last 1.2 *μ*s of the simulations (with output frequency of 120 ps) were used for all analysis except for lipid diffusion which was calculated over the trajectory lengths indicated in Table S6 with output frequency of 20 ps.

To analyze the permeability of a membrane we first identified all water molecules that at some point in time were found 10 Å or more below the instantaneous average z position of the lipid phosphate groups in either leaflet. We then recorded the time evolution of the z coordinates of each of these water molecules throughout the trajectory frames. This data was then used to count the number of times water molecules entered the bilayer from one side, reached within 10 Å of the bilayer midplane, i.e. with −10 ≤ 𝑧 ≤ 10 Å, and either moved through the opposite leaflet and exited from the other side (full transitions) or turned back and exited from the same side (partial transitions). The number of full and partial transitions was normalized by the lateral area of the bilayer and the total simulation time of the analyzed portion of the trajectory.

Defects were calculated with the PACKMEM tool ^15^ in 2 dimensions and with the algorithm from^16^ in 3 dimensions as detailed in the SI.

To analyze the lipid and Chol diffusion from each trajectory, we first took the raw coordinates of the lipid phosphate atoms (P in CHARMM36 notation) or Chol’s oxygen atom (O3 in CHARMM36 notation) from every trajectory frame output at a frequency of 20 ps, and unwrapped them (that is, removed any jumps in the coordinates between frames due to wrapping around the periodic boundaries). We then used the time evolution of these atoms’ (𝑥, 𝑦) positions to calculate the 2-dimensional mean squared displacement, 𝑚𝑠𝑑(𝑡), and average it over all lipids (or all Chol molecules) in each leaflet. A small region of the 𝑚𝑠𝑑(𝑡) function was then fit to 𝑓(𝑡) = 4𝐷𝑡 where 𝐷 is the diffusion coefficient. This small region was chosen with a heuristic approach where consecutive 100-ns-long blocks of the 𝑚𝑠𝑑(𝑡) function were fit to 𝑔(𝑡) = *ax^b^* and the block with the largest 𝑏 parameter was the one used for obtaining 𝐷. Representative plots of the data and fits is shown in Figure S14. Errors on the diffusion coefficients for each leaflet were calculated with a bootstrapping approach in which a dataset with the same total number of lipids was constructed by picking lipids at random, then the respective 𝑚𝑠𝑑(𝑡) profiles were averaged and analyzed as above. The procedure was repeated multiple times and the standard deviation of the resulting diffusion coefficients was reported.

Further details can be found in Supplements.

## Supplementary Materials

Supplementary Text

Figs. S1 to S18

Tables S1 to S6

References

