## Supplemental Material for "Cell membranes sustain phospholipid imbalance via cholesterol asymmetry"

**The PDF file includes:**

Supplementary Text
Figs. S1 to S18
Tables S1 to S6
References

#### Supplementary Text


1. **Lipidomics**
   1.1. Protocol overview
   1.2. Sample preparation
   1.3. Data analysis
   1.4. Hemolysis measurements
   1.5. Estimation of the abundances of other unreported membrane components

2. **MD simulations**
   2.1. Tolerance for PL imbalance: CG simulations
   2.2. Effect of PS flip-flop on tension: AA simulations
   2.3. Cholesterol distribution in asymmetric membranes: CG/AA simulations
   2.4. PM models: AA simulations
   2.5. Calibration of APL and Di4 lifetime: AA simulations

3. **In vitro experiments with liposomes**
   3.1. Cholesterol and membrane tolerance for PL imbalance: LUVs
   3.2. DHE self-quenching in liposomes
   3.3. FRET experiments with DHE and Di4: LUVs
   3.4. Calibration of APL and Di4 lifetime: GUVs

4. **In vivo experiments with erythrocytes**
   4.1. DHE exchange
   4.2. DHE quenching
   4.3. Scrambling
   4.4. Permeability measurements

5. **Experiments with nucleated cells**
   5.1. Chol extraction and PM scrambling/permeability
   5.2. Di4 lifetime measurements in untreated and scrambled 3T3 cells
   5.3. Cholesterol and lipid diffusion in 3T3 cells
   5.4. Peptide binding to cell PM
   5.5. Manipulation of plasma membrane lipids in RBL cells
   5.6. Imaging of cholesterol-binding proteins, C-laurdan and lipid droplets

### 1. Lipidomics

#### 1.1 Protocol overview

Human erythrocytes (freshly isolated and intact) were treated with two enzymes, phospholipase 2 (PLA2) and sphingomyelinase (SMase) to digest exclusively the lipid species on the exoplasmic PM leaflet. Enzyme-treated cells were compared to untreated controls to determine the abundance of ~600 unique phospholipid species on the exoplasmic leaflet via the extent of digestion. The analysis included all major phospholipid species in erythrocytes together with glycosphingolipids (GSLs). Hemolysis was monitored to ensure that the integrity of the erythrocytes was preserved, and enzyme treatment of sonically disrupted cells verified the optimality of experimental conditions (i.e., the enzymes successfully degraded all available target lipids). Additional controls including lack of spontaneous phospholipid flipping on the experimental timescales can be found in (Lorent et al., 2020).

The lipidome of the inner leaflet was inferred from the lipid species remaining after digestion of intact cells, while the outer leaflet was calculated by subtracting the abundance of each lipid species remaining after digestion from the untreated controls for each target of the enzyme used. These data were highly consistent between multiple healthy adult donors (N = 3) and repeated samples from the same donor. These measurements produced raw lipidomes of the inner and outer leaflets. The total lipid amounts per leaflet were determined by adding the mol% of each individual species and averaging across experiments (n > 3).

Quantitative accuracy of these approaches is supported by the fact that the products of the lipolysis reactions (lysolipids and ceramide) are quantitatively equivalent to the abundances of the parent lipids (PL and SM) consumed by the reaction (see supplement of (Lorent et al., 2020)).

#### 1.2 Sample preparation

Samples (300 µl) of packed, freshly isolated (see Section 4.1), intact human erythrocytes from healthy donors with informed consent were treated with 10 IU PLA2 (Apis mellifera) or 0.5 IU SMase (Bacillus cereus) in 5 ml isotonic saline solution (50 mM Tris HCl, 0.25 mM CaCl2, 0.25 mM MgCl2, 150 mM NaCl, pH 7.4) for 30 min at 30 °C to specifically digest only the lipid species present on the exoplasmic leaflet of the PM. After treatment for the indicated time, the cells were fast-frozen in liquid nitrogen, and their detailed lipid compositions were analyzed by shotgun electron spray ionization with tandem MS-MS (ESI-MS/MS) by Lipotype and detailed below. Lipidomes were prepared from at least three independent human donors for all experiments, using the following procedures.

**Lipid extraction for mass spectrometry lipidomics**

Mass spectrometry-based lipid analysis was performed by Lipotype GmbH (Dresden, Germany) as described (Surma et al. 2021). Lipids were extracted using a two-step chloroform/methanol procedure (Ejsing et al. 2009). Gangliosides were extracted from the water phase of the preceding chloroform/methanol extraction with a solid phase extraction protocol (Porter et al., 2021). Samples were spiked with internal lipid standard mixture containing: cardiolipin

14:0/14:0/14:0/14:0 (CL), ceramide 18:1;2/17:0 (Cer), diacylglycerol 17:0/17:0 (DAG), hexosylceramide 18:1;2/12:0 (HexCer), dihexosylceramide 18:1;2/12:0 (DiHexCer), Globoside 3 18:1;2/17:0 (Gb3), GM3-D3 18:1;2/18:0 (GM3), GM1-D3 18:1;2/18:0 (GM1), lyso-phosphatidate 17:0 (LPA), lyso-phosphatidylcholine 12:0 (LPC), lyso-phosphatidylethanolamine 17:1 (LPE), lyso-phosphatidylglycerol 17:1 (LPG), lyso-phosphatidylinositol 17:1 (LPI), lyso-phosphatidylserine 17:1 (LPS), phosphatidate 17:0/17:0 (PA), phosphatidylcholine 15:0/18:1 D7 (PC), phosphatidylethanolamine 17:0/17:0 (PE), phosphatidylglycerol 17:0/17:0 (PG), phosphatidylinositol 16:0/16:0 (PI), phosphatidylserine 17:0/17:0 (PS), cholesterol ester 16:0 D7 (CE), sphingomyelin 18:1;2/12:0;0 (SM), sulfatide d18:1;2/12:0;0 (Sulf), triacylglycerol 17:0/17:0/17:0 (TAG) and cholesterol D6 (Chol). Gb4 was estimated semi-quantitatively, using the Gb3 standard. After extraction, the organic phase was transferred to an infusion plate and dried in a speed vacuum concentrator. 1st step dry extract was re-suspended in 7.5 mM ammonium acetate in chloroform/methanol/propanol (1:2:4; V:V:V) and 2nd step dry extract in 33% ethanol solution of methylamine in chloroform/methanol (0.003:5:1; V:V:V). All liquid handling steps were performed using Hamilton Robotics STARlet robotic platform with the Anti Droplet Control feature for organic solvents pipetting.

**MS data acquisition**

Samples were analyzed by direct infusion on a QExactive mass spectrometer (Thermo Scientific) equipped with a TriVersa NanoMate ion source (Advion Biosciences). Samples were analyzed in both positive and negative ion modes with a resolution of $R_{m/z=200}$=280000 for MS and $R_{m/z=200}$=17500 for MSMS experiments, in a single acquisition. MSMS was triggered by an inclusion list encompassing corresponding MS mass ranges scanned in 1 Da increments (Surma et al., 2015). Both MS and MSMS data were combined to monitor CE, Chol, DAG and TAG ions as ammonium adducts; LPC, LPC O-, PC, PC O-, as formiate adducts; and CL, LPS, PA, PE, PE O-, PG, PI and PS as deprotonated anions. MS only was used to monitor LPA, LPE, LPE O-, LPG and LPI as deprotonated anions; Cer, HexCer, and SM as formiate adducts and cholesterol as ammonium adduct of an acetylated derivative (Liebisch et al., 2006).

**Data analysis and post-processing**

Data were analyzed with in-house developed lipid identification software based on LipidXplorer (Herzog et al., 2012; Herzog et al., 2011). Data post-processing and normalization were performed using an in-house developed data management system. Only lipid identifications with a signal-to-noise ratio >5 and a signal intensity 5-fold higher than in corresponding blank samples were considered for further data analysis.

##### 1.3 Data analysis

Lipidomic analysis yielded a list of >600 individual lipid species and their picomolar abundances. These were processed by first removing the TAG and sterol esters from the analysis, and expressing the remaining lipids as mol% of membrane lipids. Within a class (depending on which enzyme was used per experiment) each lipid species was then compared separately for each individual biological replicate (that is, the enzymatically treated samples were directly compared to the untreated parallel sample for each individual experiment to control for variance across erythrocyte isolation or human donor). The outer leaflet contents of lipid $L$ ($\chi_L^{\mathrm{out}}$) were obtained by subtracting the relative abundance of $L$ in the enzyme-treated samples ($\chi_L^{\text{enz}}$) from that in the untreated samples ($\chi_L$) with each being expressed as mol% of total membrane lipids:

$$\chi_L^{\text{out}} = \chi_L - \chi_L^{\text{enz}}.$$

Each species calculated to be on the inside (that is, $\chi_L^{\text{enz}}$) was summed to give the total inner leaflet composition, and each species calculated to be on the outside was summed to give the total outer leaflet composition. All measured glycosphingolipids were assumed to be on the outer leaflet. These values then report on the total contents of lipids in each leaflet. The ranges reported here for each lipid class are determined from the mean and standard deviations for the individual lipid species within the lipid class. To obtain the range of total lipid abundances for each leaflet, the means of the lipid classes from that leaflet were summed, and their errors propagated accordingly.

##### 1.4 Hemolysis measurements

For all enzyme treatments, hemolysis was monitored to ensure that the erythrocytes remained intact, and only conditions that yielded no hemolysis were considered for asymmetry studies. Hemolysis was quantified by measuring the absorbance of the supernatant of the enzyme-treated cells at 540 nm on a Tecan plate reader. As an additional test, erythrocytes were incubated with FITC-dextran (MW = 3 kDa) during treatment with SMase and imaged on a Nikon A1R laser scanning confocal microscope immediately after treatment to ensure no leakage of dye into the erythrocyte cytoplasm.

##### 1.5 Estimation of the abundances of other unreported membrane components

To estimate the proportion of GPI-anchored proteins as a mole fraction of erythrocyte lipids, we used published values to estimate the abundance of both. From (Kuchel et al., 2021), we estimated the surface area of an erythrocyte as ~150 $\mu m^2$, subtracted 20% of this surface area for transmembrane domain-containing proteins (Dupuy and Engelman, 2008) and another 20% for cholesterol, giving a total erythrocyte phospholipid surface area of 90 $\mu m^2$ ($9\times10^9$ Å$^2$). Assuming a conservative average phospholipid surface area of 60 Å$^2$ (Lorent et al., 2020), the approximate number of phospholipids to cover the outer leaflet erythrocyte surface area is thus on the order of $1.5\times10^8$.

The four abundantly expressed GPI-APs in erythrocytes are CD59, DAF, Semaphorin-7a, and AChE (Bryk and Wisniewski, 2017; Pasini et al., 2006; Ravenhill et al., 2019). The expression levels (molecules/erythrocyte) of CD59 and DAF have been estimated using biochemical approaches as being $2\times10^4$ (Kooyman et al., 1995) and $3.3\times10^3$ (Kinoshita et al., 1985; Kooyman et al., 1995), respectively. A more recent proteomic analysis (Bryk and Wisniewski, 2017) attempted to quantify protein copy numbers in erythrocytes, with impressive consistency with the older biochemical measurements (e.g. CD59 and DAF detected in the range of $1\times10^3$ - $3\times10^4$ molecules/erythrocyte). This analysis also reported AChE (~1-$5\times10^3$) and Semaphorin-7a copy numbers (0.9-$2.2\times10^4$). None of the other known erythrocyte GPI-APs had abundance above $0.5\times10^3$. Thus, the combined GPI-APs comprise ~$5\times10^4$ molecules / erythrocyte, which compared to ~$1.5\times10^8$ phospholipids in the outer leaflet suggests them to be a negligible fraction of the membrane lipidome.

Transmembrane proteins comprise ~20% of plasma membrane area (Dupuy and Engelman, 2008; Takamori et al., 2006), which would not be sufficient to counterbalance the area imbalances implied by differential phospholipid abundances of the two PM leaflets we report. Moreover, transmembrane domains are generally asymmetric in their shape, occupying more area in the inner leaflet (Lorent et al., 2020; Sharpe et al., 2010; Takamori et al., 2006), which would exacerbate rather than alleviate the area mismatch due to phospholipid imbalances.

Finally, there are likely lipid components that are not identified in our lipidomics analysis. The most prominent among these are phosphoinositide lipids, whose hydrophilicity and charge present a challenge for quantitative shotgun lipidomics; however, phosphoinositides have been reported to comprise on the order of 1 mol% of cellular lipids (Mucksch et al., 2019). Similarly, lysolipids, cardiolipins, ceramides, etc each comprise <1 mol% each. Other minor components (including potentially yet-unidentified ones) are likely even less abundant.

#### 2. Molecular Dynamics (MD) simulations

##### 2.1 Tolerance for PL imbalance: CG simulations

To examine the effect of cholesterol (Chol) on the tolerance of lipid membranes to phospholipid (PL) imbalance, we built a series of asymmetric membranes with a lipid composition mimicking that of the erythrocyte plasma membrane but with fixed PL imbalance (35% more PLs in the cytoplasmic leaflet) and varying amounts of Chol (ranging between 10 and 50 mol% of all membrane lipids). Table S2 summarizes the initial lipid make-up of the membranes.

*Bilayer construction and simulation.* All bilayers were built with the Martini bilayer builder in CHARMM-GUI (Lee et al., 2019). In addition to the bilayer lipids each system was solvated with water molecules comprising a 40 Å-thick layer above and below the bilayer, charge-neutralized with $Na^+$ ions, and additional $Na^+$ and $Cl^-$ ions were added amounting to a final salt concentration of 150 mM (the number of ions was determined by CHARMM-GUI).

Equilibration of the systems was carried out in Gromacs (Abraham et al., 2015) with the Martini 2.2 force-field (de Jong et al., 2013), following the 6-step protocol provided by CHARMM-GUI. After equilibration all cholesterol molecules were replaced with a recently optimized cholesterol model (Fabian et al., 2023) which circumvents previously reported problems with cholesterol in Martini (Thallmair et al., 2021).

A subsequent production run of 10 $\mu$s was carried out with Gromacs. The simulations were run with a timestep of 20 fs at a temperature of 45°C. Van der Waals interactions were modeled with a Potential-shift-verlet algorithm and a cutoff of 1.1 nm. Long-range electrostatic interactions were modeled with the Martini 2 reaction field electrostatics and a Coulomb cutoff of 1.1 nm. The lincs_iter and lincs_order parameters were set to 1 and 4, respectively, in accordance with the parameterization of the improved Chol model (Fabian et al., 2023). The remaining simulation parameters included the standard set of input parameters for the production run provided by CHARMM-GUI including a semi-isotropic pressure coupling with the Parrinello-Rahman barostat and a reference pressure of 1 bar, and temperature control with velocity rescaling and a time constant for coupling of 1 ps.

*Simulations analysis.* Cholesterol interleaflet distribution was determined from the orientation of the Chol molecules. In particular, the angle between the vector defined by the ROH and R5 beads of Chol was calculated with respect to the bilayer normal (0,0,1) from the last 5 $\mu$s of the trajectories. All molecules with tilt angles less than 90 degrees were counted in the top exoplasmic-like leaflet and all molecules with tilt angles greater than or equal to 90 degrees were counted in the bottom cytoplasmic-like leaflet. Phospholipids did not change their leaflet residence during the simulations.

The maximal phosphate-to-phosphate distance was calculated from the difference between the minimum and maximum z positions of all phosphate (PO4) beads in each frame of the trajectory, and plotted as histograms.

#### 2.2 Effect of PS flip-flop on tension: AA simulations

To evaluate the effect of flipping PS lipids to the opposite leaflet, we performed a series of all-atom (AA) simulations in which the total number of lipids in the bilayer remained fixed and we mimicked PS flipping by generating different starting configurations with respect to PS interleaflet distribution. The effects of Chol flipping were examined in the same way. These simulations were relatively short (not much more than a microsecond) with the goal of equilibrating lipid packing for calculation of the respective leaflet tensions while avoiding any spontaneous translocation of lipids and Chol between leaflets.

*Bilayer construction and simulation*. All bilayer systems were constructed with CHARMM-GUI's Bilayer Builder. Their lipid compositions and system sizes are listed in Table S3. The bilayers were initially equilibrated in NAMD (Phillips et al., 2020) with CHARMM-GUI's 6-step equilibration protocol using the CHARMM36 force-field (Yu et al., 2021). The same software and force field were used for the subsequent production runs which utilized a 2 fs timestep with *rigidbonds* set to all. Van der Waals interactions were truncated at 12 Å and CHARMM's force switching function was applied between 10 and 12 Å with the *vdwforceswitching* option in NAMD. Electrostatic interactions were modeled with the Particle Mesh Ewald (PME) method with a grid spacing of 1 Å. The simulations were run in the NPT ensemble with a semi-isotropic pressure coupling. The temperature and pressure were maintained at 37°C and 1 atm respectively using Langevin thermostat (damping coefficient of 5 $ps^{-1}$) and barostat (period 200 fs and decay 50 fs). The total lengths of these production runs are indicated in Table S3.

*Calculation of leaflet tension.* The tension in each leaflet of the simulated asymmetric bilayers was calculated as described previously (Doktorova and Weinstein, 2018). Briefly, each system was first centered so that the average $z$ position of the terminal methyl carbons of all phospholipids was in the center of the simulation box at $z = 0$ (note that the results did not change significantly when we calculated the average $z$ positions of the terminal methyl carbons of the two leaflets separately and centered the bilayer using their mean instead). The lateral pressure profile of each system was computed with NAMD 2.11 since later versions have a bug preventing the calculation.

The system was divided in $N_\mathrm{s}$ slabs where $N_\mathrm{s}$ was chosen so that the average slab thickness was ~0.8 Å, and the non-Ewald and Ewald (with Ewald grid size in each direction set to 30) contributions to the pressure tensor were calculated separately from the stored atomic coordinates and velocities with the Harasima contour, and summed to obtain the final X ($p_{xx}$) and Y ($p_{yy}$) components of the pressure tensor. The normal Z component of the pressure tensor ($p_N$) in each slab was estimated as the constant $p_N = L_N^{-1}\int \frac{1}{2}\left(p_{xx}(z)+p_{yy}(z)\right)\mathrm{d}z$ where $L_N$ is the length of the simulation box along the bilayer normal. The pressure profile was then computed as $p(z) = \frac{1}{2}\left(p_{xx}(z)+p_{yy}(z)\right) - p_N$ and the tension in each leaflet was obtained by integrating $p(z)$ from $-\infty$ to 0 (bottom leaflet) or from 0 to $+\infty$ (top leaflet) and multiplying the result by $(-1)$. Errors on the tension were estimated with a bootstrapping approach in which a dataset with the same total number of frames was reconstructed by picking frames from the original dataset at random and allowing repetitions. The pressure profile and leaflet tensions were then obtained from the resampled data. This procedure was repeated multiple times to obtain the error (i.e. standard deviation) of the leaflet tension.

#### 2.3 Cholesterol distribution in asymmetric membranes: CG/AA simulations

To examine the effects of phospholipid imbalance and tension on cholesterol distribution in asymmetric membranes, we performed a series of simulations first in coarse-grained (CG) and then in all-atom (AA) representations.

*Coarse-grained simulations*. Initially, 8 asymmetric bilayers with one saturated leaflet (composed of 100 DPPC lipids) and one unsaturated leaflet (composed of a varying number of DAPC lipids), each containing 30 mol% cholesterol (Chol), were constructed with the Martini maker in CHARMM-GUI. Table S4 lists the starting configurations of all bilayers. The systems were equilibrated with CHARMM-GUI's 6-step protocol and simulated at 40°C for 10 $\mu$s with Gromacs. The standard set of parameters for the production run are the same ones as listed above in Section 3.2 and included a timestep of 20 fs, 1.1 nm cutoffs for Coulomb and Van der Waals interactions, and Martini 2 reaction field electrostatics. While the lincs_iter and lincs_order parameters were set to 1 and 4 respectively, as above, the simulations were run with the old Martini 2.2 version of cholesterol since the newer model was not available at the time. For two of the systems, (A.5 and A.10), we later reran the simulations with the new Chol model(Fabian et al., 2023) and verified that there were no observable differences in their equilibrium Chol interleaflet distributions.

To count the number of lipids and Chol in each leaflet, the bilayer was first centered so that the bilayer midplane was in the center of the simulation box in every frame. Then, the DAPC and DPPC lipids whose phosphate (PO4) beads were below/above the bilayer center were counted as belonging to the bottom/top leaflets respectively. Chol was treated the same way except that its headgroup OH bead was used instead. Chol distribution equilibrated within the first few $\mu$s in all systems and all reported quantities are averaged over the last 5 $\mu$s of the trajectories. None of the phospholipids changed its leaflet residence during the simulations.

*All-atom simulations*. The equilibrated lipid distributions from the CG simulations were used to construct all-atom bilayers with CHARMM-GUI's Membrane Builder. Table S4 lists the respective lipid leaflet compositions. The bilayers were hydrated with more than 40 waters per lipid and no added salt. They were simulated with NAMD 2.12 first following CHARMM-GUI's 6-step equilibration protocol, then a production run utilizing a 2 fs timestep with *rigidbonds* set to all. Van der Waals interactions were truncated at 12 Å and CHARMM's force switching function was applied between 10 and 12 Å with the *vdwforceswitching* option in NAMD. Electrostatic interactions were modeled with the Particle Mesh Ewald (PME) method with a grid spacing of 1 Å. The simulations were run in the NPT ensemble with a semi-isotropic pressure coupling. The temperature and pressure were maintained at 40°C and 1 atm respectively using Langevin thermostat (damping coefficient of 5 $ps^{-1}$) and barostat (period 200 fs and decay 50 fs). The total lengths of these production runs are indicated in Table S4.

*Calculation of leaflet tension.* The tension in each leaflet of the simulated asymmetric bilayers was calculated from the AA trajectories as described in Section 3.2.

#### 2.4 PM models: AA simulations

To examine the effects of PL imbalance on cholesterol interleaflet distribution and bilayer properties we built 3 asymmetric models of the plasma membrane. Each had ~40 mol% overall cholesterol and the same leaflet phospholipid compositions modeled after the lipidomics results, but differed in the relative total abundances of PLs in their two leaflets (see Table S5 for the exact numbers and types of lipids in each leaflet). The bilayers were constructed and simulated as described below.

*Bilayer construction.* The "Equal" bilayer (asymmetric lipid composition, but matched leaflet areas) was built by first simulating two symmetric bilayers, one with the inner leaflet composition and one outer leaflet composition (previously published in (Lorent et al., 2020)). After the areas of each were relaxed, an asymmetric bilayer was constructed by taking one leaflet from each simulation, placing them tail to tail, and resolvating with the VMD plugins "solvate" and "ionize." The final system contained ~69,000 water molecules (~53 waters per lipid) and sodium and chloride ions for a final salt concentration of 156 mM. The system was then relaxed using the 6-step CHARMM-GUI protocol described below.

The other two PM models with larger PL imbalances were built with CHARMM-GUI with the initial numbers of PLs and Chol in each leaflet as indicated in Table S5. At the time of construction some lipids were not available in CHARMM-GUI and the bilayers were first constructed with similar available lipids whose structure was subsequently modified. These lipids were as follows: DGPE (20:1,20:1 PE) in place of OAPE; PAPC in place of PAPS; and DEPE (22:1,22:1 PE) in place of PDPE and PLQS. After the structural modifications the systems were hydrated with ~25,000 water molecules (~80.5 waters per lipid) and sodium and chloride ions for a final salt concentration of 150 mM. The hydration and ionization were done with the solvate plugin in VMD.

Additional symmetric bilayers with the lipid compositions of the exo (exo-sym), cyto (cyto-sym) leaflet or overall bilayer (scramble) of the Cyto+ model were built in a similar way. Their lipid compositions were as follows: LSM/NSM/PSM/PAPC/PLPC/SOPC/Chol 7/8/10/3/11/6/55 mol% in exo-sym, OAPE/PAPS/PDPE/PLPC/PLQS/POPC/Chol 6/22/13/15/15/6/23 mol% in exo-cyto and LSM/NSM/PSM/OAPE/PAPC/PAPS/PDPE/PLPC/PLQS/POPC/SOPC /Chol 4/4/6/3/2/9/6/13/6/3/3/41 mol% in scramble.

*Simulation protocol.* All systems were first equilibrated with CHARMM-GUI's 6-step equilibration protocol in NAMD (an updated list of dihedral angle constraints was generated and used instead of the one supplied by CHARMM-GUI after the structural modifications of the lipids described above). The systems were then transferred to Anton2 for long production runs as detailed in Table S6. The Anton-specific system set-up and parameters were as described in (Lorent et al., 2020). The final snapshots of the simulations were then used to start subsequent NAMD production runs with simulation lengths outlined in Table S6. The NAMD runs utilized the same simulation parameters as those described in the All-atom simulations in Section 3.3.

*Analysis.* Prior to analysis the system was centered so that the bilayer was in the middle of the simulation box and the average (x,y,z) of the bilayer midplane was at (0,0,0). The last 1.2 $\mu$s of the simulations (with output frequency of 120 ps) were used for all analysis except for lipid diffusion which was calculated over the trajectory lengths indicated in Table S6 with output frequency of 20 ps.

*Lipid area, order and density.* Individual lipid areas were calculated with the Area per Lipid tool in MEMBPLUGIN (Guixa-Gonzalez et al., 2014) which performs Voronoi analysis with user-specified sets of atoms defining the position of each type of lipid in the bilayer. Here the corresponding definitions consisted of the three glycerol backbone carbon atoms for all phospholipids and the oxygen atom for cholesterol.

The acyl chain order parameters were calculated with custom Tcl and MATLAB scripts with the formula:

$$S_{\mathrm{CD}}(C_i) = \langle \frac{1}{2}(3\cos^2\beta - 1)\rangle$$

where $\beta$ is the angle between a CH bond at carbon $C_i$ and the bilayer normal (the $z$ dimension of the simulation box) and $\langle\cdot\rangle$ denotes ensemble average over all CH bonds, lipids and trajectory frames.

The 3-dimensional bilayer density was calculated with the VolMap plugin in VMD at a resolution of 1 Å in a 25 x 25 x 40 Å box centered in the middle of the simulation box. The reported density was the average over the analyzed trajectory frames.

*Differential stress and first moments.* The differential stress (tension/compression) in the leaflets was calculated from the lateral pressure profile of the bilayer as described in Section 3.2. The first moments were obtained by integrating the product $zp(z)$ over each leaflet. i.e. $0 \rightarrow +\infty$ or $0 \rightarrow -\infty$ for the top and bottom leaflets, respectively.

*Water permeability.* To analyze the permeability of a membrane we first identified all water molecules that at some point in time were found 10 Å or more below the instantaneous average z position of the lipid phosphate groups in either leaflet. We then recorded the time evolution of the z coordinates of each of these water molecules throughout the trajectory frames. This data was then used to count the number of times water molecules entered the bilayer from one side, reached within 10 Å of the bilayer midplane, i.e. with $-10 \leq z \leq 10$ Å, and either moved through the opposite leaflet and exited from the other side (full transitions) or turned back and exited from the same side (partial transitions). The number of full and partial transitions was normalized by the lateral area of the bilayer and the total simulation time of the analyzed portion of the trajectory.

*Defects.* Defects in the bilayer, defined broadly as inhomogeneities on the surfaces of the two leaflets, were calculated with two different algorithms. The first one utilized the PACKMEM tool as detailed in (Gautier et al., 2018) which performs a 2-dimensional analysis of the defects in each leaflet. The output included the numbers of shallow and deep defects defined as above or below 1 Å under the glycerol backbone surface of the lipids, and the mean and standard deviation of the average defect sizes calculated from block averaging.

The second algorithm analyzed defects in 3D as detailed in (Tripathy et al., 2020). The reference atoms were the same as in (Tripathy et al., 2020), that is C2 for non-sphingomyelin and C2S for sphingomyelin phospholipids and O3 for cholesterol in CHARMM36 notation. The output contained the distributions of both defect sizes and their depths, and the defect size constant was obtained from the former by taking the inverse of the slope of the semi-log relationship (size vs. log of distribution) in the size range 150–250 Å$^2$.

*Calculation of diffusion coefficients*. To analyze the lipid and Chol diffusion from each trajectory, we first took the raw coordinates of the lipid phosphate atoms (P in CHARMM36 notation) or Chol's oxygen atom (O3 in CHARMM36 notation) from every trajectory frame output at a frequency of 20 ps, and unwrapped them (that is, removed any jumps in the coordinates between frames due to wrapping around the periodic boundaries). We then used the time evolution of these atoms' $(x, y)$ positions to calculate the 2-dimensional mean squared displacement, $msd(t)$, and average it over all lipids (or all Chol molecules) in each leaflet. A small region of the $msd(t)$ function was then fit to $f(t) = 4Dt$ where $D$ is the diffusion coefficient. This small region was chosen with a heuristic approach where consecutive 100-ns-long blocks of the $msd(t)$ function were fit to $g(t) = ax^b$ and the block with the largest $b$ parameter was the one used for obtaining $D$. Representative plots of the data and fits is shown in Figure S14. Errors on the diffusion coefficients for each leaflet were calculated with a bootstrapping approach in which a dataset with the same total number of lipids was constructed by picking lipids at random, then the respective $msd(t)$ profiles were averaged and analyzed as above. The procedure was repeated multiple times and the standard deviation of the resulting diffusion coefficients was reported.

#### 2.5 Calibration of APL and Di4 lifetime: AA simulations

To calibrate the Di4 lifetime obtained from GUV experiments (see section 4.4) to the area per lipid calculated from MD simulations, we performed a series of AA simulations of symmetric bilayers with the same lipid compositions as those of the experimentally measured liposomes. All bilayers were constructed, equilibrated and simulated as outlined in section 2.2, except that the temperature of the simulations was set to 25°C to match the experimental conditions. Each bilayer had 100 lipids per leaflet (including Chol) and the reported area per lipid was calculated by dividing the equilibrated lateral box size by the number of lipids in the leaflet (i.e. 100). Note that the data for the DOPC bilayers with 0, 10, 20, 30, 40 and 50 mol% Chol was taken from (Chakraborty et al., 2020). The lengths of the new simulations analyzed here were as follows: 710, 704, 970, 935, 1034 and 1175 ns for POPC with 0, 10, 20, 30, 40 and 50 mol% Chol respectively, 570 ns for DOPC/DOPS 80/20 and 1400 ns for DPPC/Chol 70/30.

#### 3. In vitro experiments with liposomes

##### 3.1 Cholesterol and membrane tolerance for PL imbalance: LUVs

*Sample preparation.* Three types of liposomes (POPC with 0, 20 and 40% Chol) were prepared as follows. Lipids were mixed in organic solvent at the appropriate concentrations then dried first under gas nitrogen until the solvent evaporated, then in a vacuum oven at ambient temperature overnight. The dry films were warmed to 40°C, then pre-warmed water was added to a final lipid concentration of 24 mM. Hydration was done for one hour at 40°C with intermittent vortexing. The resulting multilamellar vesicles were frozen with liquid nitrogen, then thawed at 40°C for 5 min for a total of 5 freeze/thaw cycles. The samples were extruded for 31 total passes through a 100 nm pore size filter with a manual Avanti extruder either at room temperature (without Chol) or at elevated temperature of ~50°C (with Chol). The resulting large unilamellar vesicles (LUVs) were stored at room temperature and subsequent measurements were done within one week of preparation.

A concentrated stock solution of (2-Hydroxypropyl)-$\alpha$-cyclodextrin (HP$\alpha$CD, Sigma Aldrich) was prepared by weighing 14.75 g of HP$\alpha$CD in a beaker, then adding ~15 mL water and incubating at room temperature without agitation for 1–2 days until visually dissolved and uniform. The solution was then transferred to a 25 mL volumetric flask, brought up to 25 mL with water for a nominal concentration of 500 mM, and mixed by slowly inverting the flask multiple times. The concentrated stock solution contained a small amount of large micron-sized aggregates as determined by DLS. To remove these aggregates the solution was transferred to a pre-washed centricon filter device (Ultra-15, 100,000 Da molecular weight cutoff; Amicon; MilliporeSigma, Burlington, MA), and spun at 3500 x g until ~1 mL was left in the retentate. The filtrate which contained particles of sizes between 1 and 5 nm was transferred to a falcon tube and used for subsequent measurements.

POPC extraction from the liposomes was achieved by incubating LUVs with HP$\alpha$CD as follows. First, varying amounts of the HP$\alpha$CD solution were brought up to 1 mL with water and pipette-mixed until no strands of HP$\alpha$CD were visible in the solution. Then, 7 $\mu$L of the LUVs were added for a final lipid concentration of 170 $\mu$M and pipette-mixed again until the solution looked uniform. Equilibration of POPC between the liposomes and HP$\alpha$CD was very rapid and the samples were incubated at room temperature without further agitation for anywhere between a few minutes and an hour prior to measurement.

*Light scattering.* Light scattering was measured on an Anton Paar Litesizer 100 with disposable cuvettes. Immediately prior to the measurement the 1 mL-sample was pipette-mixed and the cuvette was capped. Each sample was measured at 22°C three times in a repetition series using automatic modes for filter, focus and quality control (i.e. number of runs). The raw light scattering data from each repetition was recorded and the data was used to obtain the mean and standard deviation for that sample.

*GC/MS.* To quantify and compare the amounts of extracted POPC lipids, after the light scattering measurement each sample was first spun through a pre-washed centricon filter device (Ultra-15, 100,000 Da molecular weight cutoff; Amicon; MilliporeSigma, Burlington, MA) at 3500 x g for 10–20 minutes until the retentate reached dead stop volume. The filtrate was transferred to a glass culture tube and 2 $\mu$l of DMPC (25.27 mg/mL in chloroform) were added with a glass Hamilton syringe as a standard. The sample (total volume ~ 1 mL) was lyophilized by first storing it at -80°C overnight, then dipping in liquid nitrogen immediately prior to placing in a lyophilizer overnight. The lyophilized powder was subjected to acid-catalyzed methanolysis to extract the fatty acid methyl esters for analysis with gas chromatography-mass spectrometry (GC/MS) as done previously (Doktorova et al., 2018). Briefly, 1 mL of 1 M methanolic HCl was added to the powder and vortexed until completely dissolved, then flashed with argon and capped with a Teflon-lined cap. The sample was incubated at 85°C for 1 hr and allowed to cool for a few minutes. Then, 1 mL water and 1 mL hexane were added one after the other with vigorous vortexing after each addition to extract the FAMEs. The sample was spun for 5 min at 400 x g to break the resulting emulsion and the upper phase (containing hexane) was transferred to a GC autosampler vial and brought up to about 1 mL with hexane. FAME analysis was done on an Agilent 5890A gas chromatograph (Santa Clara, CA) with a 5975C mass-sensitive detector operating in electron-impact mode and an HP-5MS capillary column (30 m x 0.25 mm, 0.25-mm film thickness). A preprogrammed column temperature routine was performed as described in detail in (Doktorova et al., 2018). Total ion chromatogram peaks were assigned and integrated with the GC/MSD ChemStation Enhanced Data Analysis software. The respective peak areas of the FAMEs corresponding to the myristoyl chain of DMPC and the palmitoyl chain of POPC were recorded and used to calculate the normalized amount of POPC in the sample.

#### 3.2 DHE self-quenching in liposomes

*Vesicle preparation.* Vesicles were prepared following the rapid solvent exchange (RSE) protocol (Buboltz and Feigenson, 1999). Briefly, lipid mixtures (each totaling 20 nmol) without and with DHE were prepared in chloroform in a glass culture tube. Prior to mixing, the mass of the tube ($m_0$) was recorded, 25 $\mu$L chloroform were added to it and lipid stock solutions were diluted appropriately so that the total volume of the 20 nmol lipid mixture in the tube was less than 100 $\mu$L. Following that, 400 $\mu$L pure water were added to the tube and the sample was subjected to the RSE protocol by applying vacuum while vortexing for 90 seconds, followed by short purging with argon. Immediately after that, the mass of the tube ($m_1$) was recorded again and 200 $\mu$L of the RSE vesicle suspension were taken out and mixed with 1.8 mL of pure water. The 2 mL sample was transferred to a 1 cm path length Quartz cuvette and measured on a spectrofluorometer as detailed below. Due to potential slight differences in the total volume of the RSE vesicle suspension (due to small variations in the applied vacuum and argon purging), the deviation of the mass of the sample, $m_{\text{sample}} = m_1 - m_0$, from the mass of 400 uL pure water measured in similar way was used to normalize the sample volume (and thus lipid and DHE concentration) across all samples.

*Data collection.* The fluorescence intensity was measured on a Horiba Fluorolog 3 spectrofluorometer by exciting the sample at 327.8 nm and collecting the emission in the range 340–640 nm. The excitation and emission slits were set to 2.5 and 3 nm, respectively and the integration time was 0.1 s. The 2 mL sample was continuously stirred during the measurement via a flea stir bar placed in the cuvette.

*Data analysis.* To compare the DHE fluorescence across samples, the full spectrum of each sample was first normalized by subtracting the spectrum of the respective lipid-only (without DHE) sample. The normalized spectra were then integrated to obtain total DHE fluorescence, and the value was corrected for the volume difference measured during sample preparation. Integration of the DHE spectra, as opposed to using the fluorescence at a fixed wavelength, was necessary due to changes in the shape of the spectrum as a result of the formation of various DHE configurational states at higher DHE concentrations (McIntosh et al., 2003) (Figure S15).

##### 3.3 FRET experiments with DHE and Di4: LUVs

*LUV preparation.* Aqueous lipid dispersions at 25 mg/mL were prepared by first mixing appropriate volumes of lipid stocks in organic solvent with a glass Hamilton syringe. The solvent was evaporated with an inert gas stream followed by high vacuum overnight. The dry lipid film was hydrated with ultrapure water above the main transition temperature of the highest-melting lipid in the sample for at least 1 h with intermittent vortex mixing. The resulting multilamellar vesicle (MLV) suspension was subjected to at least 5 freeze/thaw cycles using a −80 °C freezer, then extruded through a polycarbonate filter using a handheld mini-extruder (Avanti Polar Lipids) by passing the suspension through the filter 31 times. Unsaturated lipids were extruded at room temperature, and multicomponent mixtures were extruded at 45°C. For DHE-containing vesicles, DHE was incorporated into the lipid mixture in organic solvent (prior to drying and hydration). DHE titrations were performed by replacing cholesterol for DHE and keeping total sterol concentration constant.

*Data Collection.* Measurements were done on a PTI fluorimeter. To minimize inner filter effect (IFE), measurements were taken using a 10×2mm pathlength cuvette oriented with 2mm pathlength facing detector and minimal sample volume: 1 $\mu$L of 2.5 mg/mL LUVs was diluted into 165 $\mu$L of Ringer's solution, yielding ~15 µg/mL LUVs in the cuvette. Di4 (stock dissolved in ethanol) was first diluted 10x in Ringer's solution, then 1.5 µL from the diluted solution were added to the LUV suspension and pipette mixed (3 strikes with 100 µL volume). IFE was negligible due to minimal absorbance measured at the DHE and Di4 absorbance ranges. Excitation and emission slits were open to 5 nm and the gain was optimized for increased signal. All measurements were taken at a scan rate of 0.1 s/nm to minimize photobleaching. Samples were checked for risk of photobleaching via time-based emission scans. DHE emission spectra was obtained by excitation at 310 nm and emission between 340–500 nm at 37°C. Measurements with Di4 were taken immediately after addition with each sample of LUVs discarded after each measurement and the cuvette thoroughly washed.

##### 3.4 Calibration of APL and Di4 lifetime: GUVs

*GUV preparation.* Giant unilamellar vesicles (GUVs) were prepared by electroformation as described previously (Lorent et al., 2020). Briefly, 1 µl of the lipid solution (1:2 methanol/chloroform at 2.5 mg/mL) was applied to Pt electrodes of an electroformation chamber.

The solution was dried in vacuum for 3 h, then rehydrated in 0.4 M sucrose. Electroformation was performed at 500 Hz and 2.5 V for 2 h at 50 °C and left overnight in heat block to return to room temperature naturally.

*Data Collection.* Measurements of Di4 lifetime of GUVs were performed as previously described (Lorent et al., 2020). Briefly, GUVs were stained with Di4 by adding Di4 dissolved in a mixture of ethanol (10%) and 0.4 M sucrose (90%) to the sample at a final concentration of 250 nM (the amount of ethanol in the GUV suspension did not exceed 0.1% v/v). The sample was incubated for 5 min at room temperature, following which 10 µl of the stained GUVs were diluted into 200 µl of 0.4 M glucose solution (containing 0.5% agarose to prevent vesicle movement during FLIM). The samples were imaged at room temperature on a Leica SP8 FLIM confocal microscope with excitation at 485nm with an 80MHz pulsed laser, and the lifetime collected in an emission window of 550–800nm. The data was then fit using a 2-component reconvolution fit, and the intensity-weighted average lifetime was reported.

#### 4. In vivo experiments with erythrocytes

##### 4.1. DHE exchange in erythrocytes

*Erythrocyte isolation.* Erythrocytes were isolated from healthy donors with informed consent via the Institutional Biosafety Committee of the University of Texas Health Science Center. In short, 90$\mu$L of whole blood was obtained by finger prick method and mixed with 10$\mu$L acid citrate dextrose solution. The mixture was then layered atop equi-volume of purchased polymorph prep solution and spun at 2,000 x g for 5 minutes at room temperature. All solution except the erythrocyte pellet was aspirated, and the erythrocyte pellet was resuspended and washed three times in Ringer's solution (125 nM NaCl; 5 mM KCl; 3 mM CaCl2; 1 mM MgSO4; 30 mM HEPES; 5 mM glucose; pH 7.4).

*Lipid extraction from erythrocytes for DHE quantification*. Lipids were extracted using a two-step extraction protocol (Ejsing et al., 2009). First, a 5x volume of 10:1 chloroform:methanol (v:v) solution was added to cells followed by nutation at room temperature for 30 minutes. The sample was centrifuged at 2350 rcf for 15 minutes and the organic (bottom) phase was transferred into a separate tube. The procedure was repeated with 3:2 chloroform:methanol, with the resulting organic phase added to the organic phase previously set aside. Finally, the sample was dried under $N_2$ and reconstituted in 100% ethanol.

*Total sterol quantification in PMs.* To identify DHE-for-cholesterol exchange conditions that did not affect total sterol levels in RBCs (see Fig S7A), the Amplex Red assay was performed (according to manufacturer instructions; Invitrogen) to determine total sterol abundance (i.e. cholesterol and DHE combined). We validated that the assay is approximately equally sensitive to DHE and cholesterol by measuring a standard curve for DHE and cholesterol separately. Each reading was normalized to cell count. Technical triplicates were measured for each sample. Values shown in Figure S7A are normalized to untreated cells and are the average and standard deviations from at least two independent biological replicates.

*DHE:MβCD complexation.* DHE was first solubilized in ethanol as follows: 5 mg DHE (Sigma E2634) were dissolved in 628 $\mu$L of 100% ethanol to give a final concentration of 20 mM. The sample was sonicated in a bath sonicator to completely solubilize any aggregates.

Next, 42.4 mg of M$\beta$CD were dissolved in 200 $\mu$L of Ringer's solution to yield final concentration of 160 mM M$\beta$CD and vortexed until clear.

To complex DHE and M$\beta$CD, 100 $\mu$L of the DHE solution was dried under $N_2$ for ~30 min and the film was dissolved with 100 $\mu$L of the M$\beta$CD solution for a MβCD to DHE molar ratio of 8:1. The M$\beta$CD solution was clear in the absence of DHE and cloudy in the presence of DHE. Full solubilization of DHE in M$\beta$CD could not be achieved with sonication or overnight incubation at 37°C with a thermomixer. Insoluble DHE aggregates were spun out with a brief spin.

*DHE exchange for cholesterol in erythrocytes.* $3.5\times10^8$ cells/mL were spun down and suspended in a mixture of the DHE/M$\beta$CD solution and free M$\beta$CD in Ringer's solution for final concentrations of 50 $\mu$M DHE and 1.2 mM M$\beta$CD, unless otherwise specified, and nutated for 10 minutes at room temperature. The sample was washed 3 times with Ringer's solution with 2,000-rcf spins for 3 minutes at room temperature. The supernatant from the first spin was saved to test for hemolysis.

*Monitoring erythrocyte morphology.* Cells were imaged by light microscopy with an epifluorescence microscope with 40x objective to assess maintenance or loss of discoid shape.

*Quantification of DHE exchanged for cholesterol in PMs.* DHE quantification was performed by measuring DHE emission spectra of lipid extracts in 100% ethanol of erythrocytes labeled with DHE by subtracting emission spectra of lipid extracts from unlabeled cells (no DHE). DHE content was calculated by direct comparison to a calibration curve of increasing DHE concentration in ethanol. Total sterol was measured with the amplex red assay as described above. The mol% DHE of total PM lipids, $\chi_{\mathrm{DHE}}^{\mathrm{PM}}$, was calculated from the amount of DHE relative to total sterol, $\chi_{\mathrm{DHE}}^{\mathrm{sterol}}$, and the mol% sterol of total PM lipids, $\chi_{\mathrm{sterol}}^{\mathrm{PM}}$, as:

$$\chi_{\mathrm{DHE}}^{\mathrm{PM}} = \chi_{\mathrm{DHE}}^{\mathrm{sterol}}\, \chi_{\mathrm{sterol}}^{\mathrm{PM}}.$$

#### 4.2. DHE quenching in erythrocytes

*Di4 addition.* Di4 was diluted 10-fold to 100 μg/mL in Ringer's buffer before adding to cells ($1.5\times10^7$ cells/mL) at room temperature at the indicated Di4 concentrations. Di4 addition was directly followed by measurement of DHE emission for quantification of quenching via FRET.

*Förster distance calculations.* The Förster distance $R_0$ for energy transfer between DHE donor and Di4 acceptor was calculated as (Lakowicz, 2006):

$$R_0 = \left(\frac{9000\kappa^2\Phi_D \ln(10)}{128\pi^5 N_A n^4} J(\lambda)\right)^{1/6} \approx 0.211\left(\kappa^2 n^{-4}\Phi_D J(\lambda)\right)^{\frac{1}{6}} \,[\text{Å}],$$

where $N_A$ is Avogadro's number, $\kappa^2$ is the unitless dipole orientation factor (2/3 in the limit of isotropic motion and dynamical averaging), $n$ is the refractive index of the medium (taken to be 1.42 for a lipid bilayer), $\Phi_D$ is the donor quantum yield (0.4 for DHE (Schroeder et al., 1987)), and $J(\lambda)$ is the spectral overlap integral:

$$J(\lambda) = \frac{\int_0^\infty F_D(\lambda)\varepsilon_A(\lambda)\lambda^4 d\lambda}{\int_0^\infty F_D(\lambda)d\lambda}.$$

In the calculation of $J$, $F_D(\lambda)$ is the DHE emission spectrum, $\varepsilon_A(\lambda)$ is the Di4 excitation spectrum normalized to a peak absorbance of $\varepsilon_A$ = 10,110 $M^{-1}cm^{-1}$, and wavelengths are in nm. Excitation and emission spectra were obtained from fluorescence spectroscopic measurements described below. Based on these calculations and assumptions R0 is 28.3 Å or 29.5 Å depending on which Di4 excitation spectrum is used.

*Fluorescence detection of DHE and Di4 via spectrofluorometer*. Measurements were taken on a PTI fluorimeter as described in Section 4.3. Cells were measured at $1.5\times10^7$ cells/mL and the temperature was controlled to 37°C. DHE emission spectra was obtained by excitation at 310 nm and emission from 340–500 nm with 2 nm step size and 0.1 s/nm integration time. Di4 emission spectra was obtained by excitation at 488 nm and emission from 550–600 nm with 2 nm step size and 0.1 s/nm integration time.

*Fluorescence measurements of DHE distribution*. Measurements were always taken for control (untreated) erythrocytes and subtracted. This was necessary since erythrocytes cause high scattering effects due to their shape and hemoglobin content. For counterpart measurements, cell concentration was checked via cell count. DHE quenching was monitored by measuring the decrease in emission at 385 nm after adding increasing amounts of Di4 until a plateau was reached. The relative amount of quenched DHE fluorescence at this plateau was used to estimate the fraction of DHE residing in the outer leaflet of the membrane.

It is important to emphasize that some fluorescence effects are not fully accounted for in this calculation. Specifically, there is the likelihood of some quenching of inner leaflet DHE from outer leaflet Di4, effects of membrane environment/hydration on DHE and Di4 spectral properties, and cell-to-cell variation in DHE and/or Di4 labeling. Fully characterizing and controlling for these effects would be necessary for quantitatively accurate measurements of DHE transbilayer distribution. However, definitive quantification of cholesterol asymmetry is not the central focus of this manuscript, rather that polar lipid imbalances and preferential lipid-lipid interactions are key determinants of this asymmetry. Our DHE measurements strongly support enrichment of sterol in the outer PM leaflet, consistent with simulations and other experiments (see Fig 4).

##### 4.3. Scrambling of erythrocytes

DHE-labeled cells were suspended in 100 $\mu$M phorbol-12-myristate acetate (PMA) for 10 minutes on nutate at room temperature, unless otherwise specified. Di4 was then immediately added to the suspension and cells were measured for DHE distribution as described in the previous section.

4.4. Permeability measurements

*Data collection.* Equal numbers of erythrocytes (~$3.4\times10^8$ cells/mL), were treated with or without 100 µM PMA for 10 minutes. $2\times10^6$ cells/mL were resuspended in Ringer's solution to a quartz 2 mL cuvette. Just prior to reading on the spectrofluorometer, 1 µg/mL fluorescein diacetate (stock 1 mg/mL in acetone) was added to the cells and mixed by pipetting. The fluorescence intensity was read at 450 nm excitation and 510 nm emission over a period of 1 min (1 read/second) with 5 nm excitation and emission slits. In each of the samples measured, the slope of the fluorescence increase over time remained constant over the 60 s, consistent with a constant flux (Q) (Figure S16).

*Calculation.* The permeability coefficient ($P$) of a membrane to a particular substrate is then defined by Fick's Law as:

$$Q = PA(\mathrm{C_{out}} - \mathrm{C_{in}})$$

where $P$ is permeability, $A$ is the area of the membrane, and $\mathrm{C_{out}}$ and $\mathrm{C_{in}}$ are the concentrations of the solute outside and inside the cells, respectively. In our experiments, $\mathrm{C_{in}} = 0$ because FDA is quickly converted into fluorescein upon entering the cell. With equal $A$ (due to same number of cells and cellular membranes) and $\mathrm{C_{out}}$ across samples, permeability can be directly compared by normalizing to control samples to evaluate changes in scrambled PMs.

#### 5. Experiments with nucleated cells

5.1. Chol extraction and PM scrambling/permeability

*Cell culture.* RBL-23T cells were purchased from ATCC and grown in EMEM media with 10% fetal bovine serum (FBS) and 1% penicillin. In all experiments, confluent cells at passage between 10 and 16 were split the day before imaging in glass-bottom plates at a 1:4 dilution.

*Reagents.* A CaR buffer containing 145 mM NaCl, 5 mM KCl, 10 mM HEPES at pH 7.4 was prepared, filtered and stored at 4°C until use. A stock solution of 70–90 mM M$\beta$CD was prepared in CaR buffer and stored at 4°C until use. A concentrated solution of $CaCl_2$ was also prepared in water, its osmolarity measured with an osmometer (Advanced Instruments) to be 83 mM, and used to add calcium to the CaR buffer when needed. A concentrated (0.5 M) stock solution of EDTA (Thermo Scientific) was used to add EDTA to the calcium-free CaR buffer when needed. Imaging buffer consisted of CaR buffer with either 0.5–0.95 $\mu$M LactC2-mClover (purified via His-tag) and 2 mg/mL Rhodamine-dextran (Sigma-Aldrich T1037, MW 4,400, dissolved in CaR buffer) or 50 $\mu$g/mL propidium iodide (PI, Alfa Aesar J66764, dissolved in water).

*Cholesterol extraction and cell imaging.* Cells were first washed with CaR buffer twice, then incubated with either 2 mM $CaCl_2$ or 2 mM EDTA in the absence or presence of 6 mM M$\beta$CD. For monitoring PS exposure and dextran permeability the buffer also contained LactC2-mClover and Rhodamine-dextran. For PI permeability the buffer contained PI. Cells and buffer were incubated at room temperature (20–22°C) and imaged at the indicated time points (30, 60 and 90 minutes). Confocal fluorescence imaging of cells was performed at 21°C using a Nikon C2+ point scanning system attached to a Nikon Eclipse Ti2-E microscope equipped with a Plan Apo Lambda 60X oil immersion objective and a Bioptech objective cooling collar.

*Data analysis.* Imaging data was analyzed with FIJI. The contour of each cell was first manually traced with a line with Line Width 5. The raw intensity of the green fluorescence (coming from the interaction of LactC2-mClover with the membrane) was recorded and normalized first to the length of the perimeter outlining the cell, then to the background of the image measured in an analogous way by selecting 3 empty regions (with no cells) from different parts of the image. PI permeability was measured by counting PI-positive and PI-negative cells using saponin treatment (dissolved in CaR buffer and added at concentration of 1.6 mg/mL) as a positive control.

##### 5.2. Di4 lifetime measurements in untreated and scrambled 3T3 cells

To determine the Di4 lifetime of the outer leaflet of 3T3 cells, cells were incubated with 1 µg/mL Di4 at 4°C for 8 min in CaR buffer (145 mM NaCl, 5 mM KCl, 10 mM HEPES) supplemented with 2 mM $CaCl_2$ and 2 mM $MgCl_2$ at pH 8.0. Cells were briefly washed twice in the same buffer at ambient temperature before imaging at room temperature using the same microscopy conditions as described in section 4.4. To prevent any contaminating signal from internalization of the dye, all images were acquired within 20 minutes. To determine the Di4 lifetime of scrambled plasma membranes, 3 µM A23187 (Cayman, #11016) was added to cells in the same CaR buffer for 15 minutes at 37°C. Following this treatment, the cells were washed with CaR buffer, and Di4 incubation and imaging occurred as above.

##### 5.3. Cholesterol and lipid diffusion in 3T3 cells

3T3 cells were maintained in DMEM medium supplemented with 10% FBS. They were seeded on glass-bottom dishes two days before the diffusion experiments. For fluorescent labeling, the fluorescent lipid analogs (Topfluor SM and Topfluor Chol, Avanti Polar Lipids) were first dissolved in DMSO at a final concentration of 1 mg/ml. Before labelling, the cells were washed twice with L15 medium to remove the full media. Later the fluorescent analogs were mixed with L15 medium at a 1:1000 ratio (final concentration of 1 µg/ml). The cells were incubated with this suspension for three to five minutes at room temperature. After that, the cells were washed with L15 twice followed by fluorescence correlation spectroscopy (FCS) measurements. To measure diffusion of scrambled plasma membranes, 3T3 cells were treated with 3 µM A23187 (Cayman, #11016) ionophore for eight to ten minutes at 37°C before lipid labelling. Scrambling was confirmed by AnnexinV-AF647 labelling (ThermoFisher A23204).

FCS was carried out using Zeiss LSM 780 microscope, 40X water immersion objective (numerical aperture 1.2). Briefly, before the measurement, the shape and the size of the focal spot was calibrated using Alexa 488 dye in water. To measure the diffusion on the membrane, the laser spot was focused on the bottom membrane of the cells by maximizing the fluorescence intensity. Then, 3-5 curves were obtained for each spot (five seconds each). To avoid the crosstalk from internalized fluorophores, measurements were done at the bottom of the nucleus. The obtained curves were fit using the freely available FoCuS-point software using 2D and triplet model (Waithe et al., 2016). Although it is possible that there are multiple diffusing populations (inner versus outer leaflet, protein interactions, lipid domains, etc) in these cellular experiments, the obtained FCS curves fit well to a single diffusing component, thus to avoid overfitting and overinterpretation, all FCS curves were fit to a single 'apparent' diffusion coefficient.

#### 5.4. Lipidated protein binding to cell PM

The plasmid SH4-GFP was a kind gift from the Sarah Veatch lab (University of Michigan). Briefly, the N-terminal sequence of Lyn that codes for the myristol- and palmitoyl-acylation sites were inserted into the pEGFP-N1 vector (M**GC**IKSKRKDKDLELKLRILQSTVPRARDPPVAT – EGFP). The tH-KtoQ and tH-KtoA plasmids were generated from the tH-mEmerald plasmid that encodes for the two palmitoylation and one farnesylation sites of tH-Ras linked to an mEmerald fluorescent protein (mEmerald – SGLRSKLNPPDESGPG**C**MS**C**K**C**VLS*). The K (underlined in previous sequence) was mutated to Q or A by site-directed mutagenesis to remove any additional electrostatic effects.

RBL cells at 70-80% confluency were harvested with 0.25% trypsin, and transfected with SH4-EGFP, mEmerald-tH-KtoQ or mEmerald-t-KtoA via electroporation. After 24 hours, cells were washed with PBS buffer, then maintained in Annexin V buffer (150 mM NaCl, 10 mM HEPES, 2 mM $CaCl_2$, pH 7.4, with 100 times diluted Annexin V AF647 from ThermoFisher, #A23204). Ionophore A23187 (Cayman, #11016) diluted in Annexin V buffer was added to the cells with a final concentration of 6 μM. Cells were imaged immediately after the addition of A23187 with a 63X water immersion objective on a Leica SP8 confocal microscope. Images were taken with 10 second intervals for up to 40 minutes.

#### 5.5. Manipulation of plasma membrane lipids in RBL cells

For extraction and loading of phospholipids from/to cells, Hexakis-(2,3,6-tri-O-methyl)-alpha-Cyclodextrin, MaCD (Cyclodextrin-Shop) was used. A stock solution was prepared in CaR buffer by incubation overnight at room temperature at a nominal concentration of 65 mM. Precipitates were removed by light centrifugation and pelleting, and the purity of the final clear solution was verified with dynamic light scattering.

RBL cells at 70-80% confluency were washed three times with CaR buffer and incubated with 6.5 mM (nominal concentration) MaCD in CaR buffer (or CaR buffer alone as parallel control) for 5 min at 37 °C. The supernatant was carefully removed and added to another washed plate of RBL cells and incubated for 5 min at 37 °C to fully saturate MaCD with outer leaflet PLs. The lipid-saturated MaCD-containing supernatant from this second incubation was then removed and spun through a 5 $\mu$m centrifugal filter to remove any contaminating cells, producing a PL-loaded MaCD solution (PL-aCD). This filtered supernatant was then added to a fresh plate of RBL cells for 15 min at 4 °C (after washing 2x in cold CaR buffer). This solution was then removed, and fresh PL-aCD was added to the same plate for another 15 min at 4 °C. For DOPC loading, DOPC (Avanti Polar Lipids) was dissolved to 10 mM in 100% ethanol. This was then diluted to 0.1 mM in CaR buffer and added to cells at room temperature for 15 min.

For sphingomyelinase (SMase) treatment, ~25,000 RBL cells were treated with 0.1 U in Tyrode's buffer (25 mM HEPES, 150 mM NaCl, 5 mM KCl, 5.4 mM glucose, 1 mM CaCl2, 0.4 mM MgCl2, pH 7.2) for 20 min at 37 °C.

*Quantification of PL-loading.* The abundance of phospholipids in the PL-aCD solutions was quantified with a colorimetric inorganic phosphate assay. After incubation of MaCD with cells to extract or load lipids, the supernatants from several samples were spun down to remove cell debris (5 min at 200xG) and a fixed volume of the top fraction (500 $\mu$L) from each was transferred to a 13 x 100 mm disposable glass culture tube (Fisher) for replicate measurements. Tubes were left uncapped on a hot plate at 94°C overnight to evaporate the solvent. The next day, tubes were removed from the hot plate and allowed to cool to room temperature. Each experiment contained at least 3 replicates and 3 controls, in addition to a set of 11 samples prepared from an inorganic phosphate standard solution (Phosphate Standard, LabChem) for a calibration curve covering a range of 0-61 nanomoles phosphate. All subsequent steps were performed identically on samples and calibration standards.

First, 200 $\mu$L of 10% (v/v) sulfuric acid (Ricca Chemical) was added to each tube. The tubes were then returned to the hot plate and the temperature was set to 200°C. After 90 min, 20 $\mu$L of 30% $H_2O_2$ (MilliporeSigma Supelco) was added directly to each tube on the hot plate followed by incubation for another 40 min. This step (addition of hydrogen peroxide and subsequent 40 min incubation at 200°C) was repeated until the solution was visibly transparent (typically four times). Tubes were then removed from the hotplate and allowed to cool to room temperature. A color reagent was prepared by mixing 0.5 mL 5% (w/v) ammonium molybdate (Thermo Scientific Chemicals) in H2O with 4.5 mL H2O and 0.1 g ascorbic acid (Spectrum Chemical), followed by vortexing until fully mixed. To each tube, 500 $\mu$L of color reagent was added followed immediately by vortexing before moving on to the next tube. After all tubes received color reagent, 500 $\mu$L of water was added to each tube followed again by immediate vortexing. The tubes were then moved to a 45°C water bath and incubated for 20 min. After removing the tubes from the water bath, absorbance was measured immediately at 820 nm on a UV/vis spectrometer (Mettler Toledo). The absorbance from each sample was converted to nanomoles phosphate using the calibration curve (Fig. S17).

*Estimation of percent loaded PL.* The measurements from the phosphate assay revealed that the transfer of PL-aCD from donor to acceptor cells added 4 nmol of PLs to ~3e5 cells, which yields ~8.0e$^9$ PL molecules/cell. From estimates of the typical surface area of mammalian cells (5,000 um$^2$) (Jaiswal et al., 2019) and the area/lipid in the PM outer leaflet (~40 Å$^2$/lipid, Fig 4C), we approximate a typical PM outer leaflet contains ~1.25e$^{10}$ lipids. According to these rough estimates, the delivered PLs represent ~40% of the PM outer leaflet after loading.

*Quantification of cholesterol.* Following incubation of cells with PL-MaCD, cells were scraped into 2% SDS, 50 mM Tris, 5 mM EDTA, pH 7.4 and sonicated for 3x 10s. An Amplex Red assay was performed (according to manufacturer instructions; Invitrogen) to determine total sterol abundance. A commercial BCA assay (Pierce) was performed to normalize the measured cholesterol amounts to total protein.

5.6. Imaging of cholesterol-binding proteins, C-laurdan, and lipid droplets

*EGFP-GRAMH, EGFP-GRAMD1a, or EGFP-GRAMD1b imagin*g. RBL cells were transfected with EGFP-GRAMH, EGFP-GRAMD1A, or EGFP-GRAMD1B (kind gift from Yasunori Saheki lab, Nanyang Technological University, Singapore) 48 h prior to the experiment via electroporation to give a final confluency of 40-50%. GRAMD1A and GRAMD1B were co-transfected with a PM-resident synthetic protein (trLAT) (Diaz-Rohrer et al., 2014). For EGFP-GRAMH, EGFP-GRAMD1A, or EGFP-GRAMD1B imaging, after PL-MaCD, DOPC, or SMase incubation, the cells were washed in CaR buffer and imaged at room temperature with a 63X water immersion objective on a Leica SP8 confocal microscope. 10-15 cells were quantified for each condition per experiment.

*C-laurdan spectral imaging.* C-laurdan imaging was performed as previously described (Owen et al., 2011; Sezgin et al., 2014). Briefly, after PL-MaCD incubation, the PL-MaCD supernatant was removed, cells washed 2x with PBS, and incubated with 5 µg/mL C-Laurdan for 20 min at RT. Finally, the cells were washed with PBS and then imaged at room temperature with a 63X water immersion objective on a Leica SP8 confocal microscope with spectral imaging with excitation at 405 nm. The emission was collected as two images: 420–460 nm and 470–510 nm. MATLAB (MathWorks, Natick, MA) was used to calculate the two-dimensional (2D) GP map, where GP for each pixel was calculated as previously described (Sezgin et al., 2015). Briefly, each image was background subtracted and thresholded to keep only pixels with intensities greater than 3 standard deviations of the background value in both channels. The GP image was calculated for each pixel with the following equation (where G is the G-factor and $I_x$ is intensity at wavelength $x$):

$$GP = \frac{\sum_{420}^{460} I_x - G\sum_{470}^{510} I_x}{\sum_{420}^{460} I_x + G\sum_{470}^{510} I_x}$$

The G-factor was determined before each experiment using a previously published protocol (Owen et al., 2011). GP maps (pixels represented by GP value rather than intensity) were imported into ImageJ. The average GP of the internal membranes was calculated by determining the average GP of all pixels in a mask drawn on each cell just inside of the PM. Histograms of the GP values for individual cells were determined and fit to two Gaussian peaks to isolated lipid droplet values from the other internal membranes. 10-15 cells were quantified for each condition per experiment.

*Lipid droplet quantification.* After PL-MaCD, DOPC, or SMase incubation, the cells were washed 2x with CaR buffer, and incubated in Tyrode's buffer for 45min at 37 °C. After 45 min, 2 $\mu$M Bodipy 493/503 (Thermo Fisher) in Tyrode's buffer was added to the cells and incubated at 37 °C for another 15 min. The cells were then washed in Tyrode's buffer 2x and imaged at room temperature with a 63X water immersion objective on a Leica SP8 confocal microscope with spectral imaging with excitation at 493 nm and emission from 503-603 nm. Z-stacks of images were taken every 1 $\mu$m from just above to just below the cell to capture the entire cellular volume. Using ImageJ, the z-stacks were converted to a maximum projection. ROIs were drawn around the perimeter of the cells. Images were then background subtracted using the mean + 3x SD of background cellular signal. The total pixel intensity and percent area covered by non-zero pixels were calculated on a per-cell basis. 15-25 cells were imaged for each condition per experiment. For ACAT inhibition the cells were treated with 1 ug/mL Sandoz 58-035 (Santa Cruz Technologies) overnight prior to the experiment and throughout the experimental incubations. For a positive control, cells were treated with 200uM cholesterol in complete medium (water soluble, Sigma Aldrich) overnight.

#### Supplemental Figures

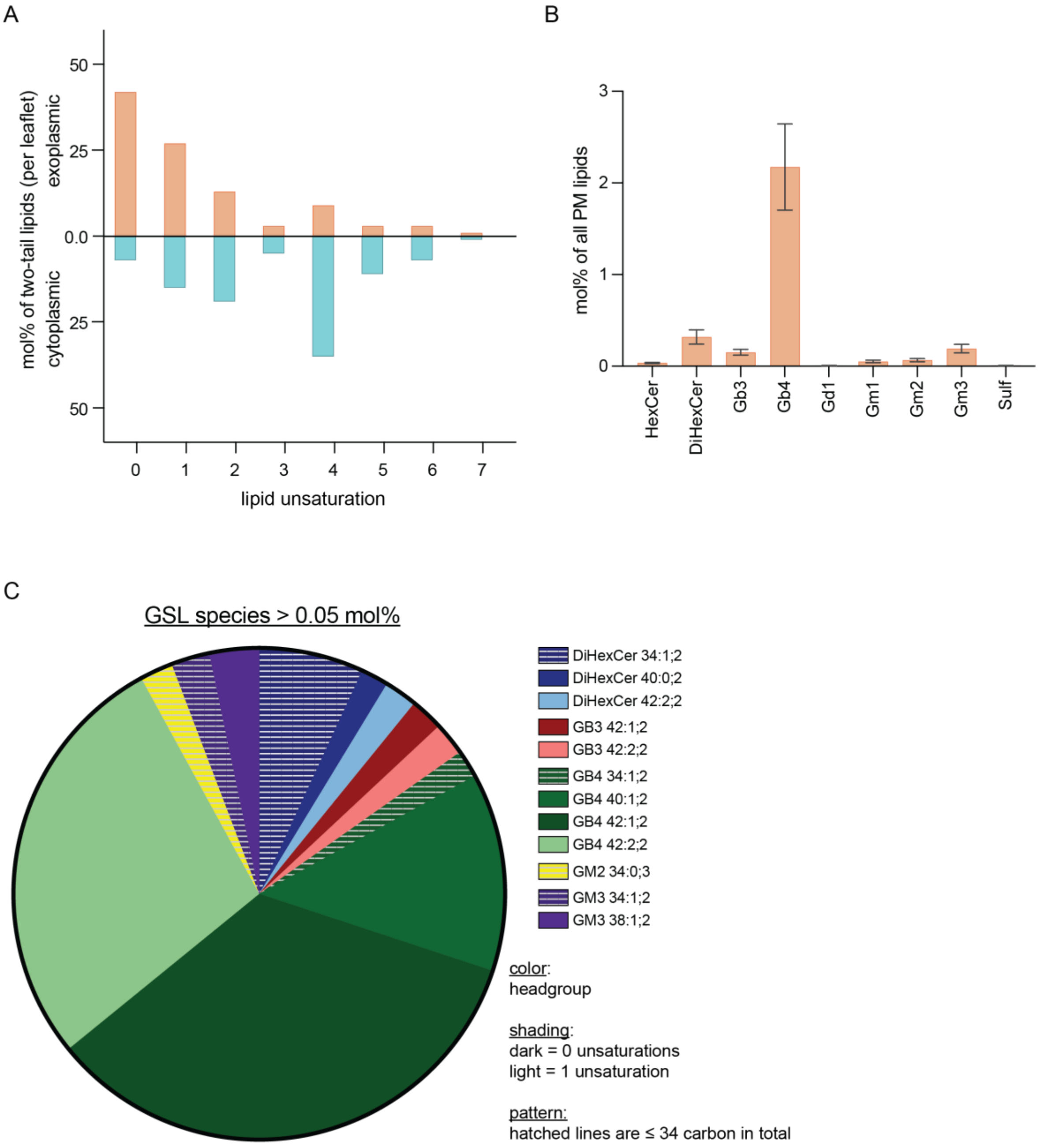


**Fig. S1. Lipidomics results including glycosphingolipids.** (A) Lipid unsaturation distribution for all lipids (including GSLs) in the exoplasmic and cytoplasmic leaflets show the exoplasmic leaflet is much more saturated. (B) Distribution of GSL headgroups as mol % of all erythrocyte lipids. (C) Abundance of all GSL species >0.05 mol% of erythrocyte lipids as a fraction of all GSLs. Color represents the GSL headgroup, shading represents the total unsaturation of the two acyl chains (not including the double bond in the ceramide backbone). Hatched bars represent shorter GSLs, with ≤34 carbons in the two acyl chains combined.

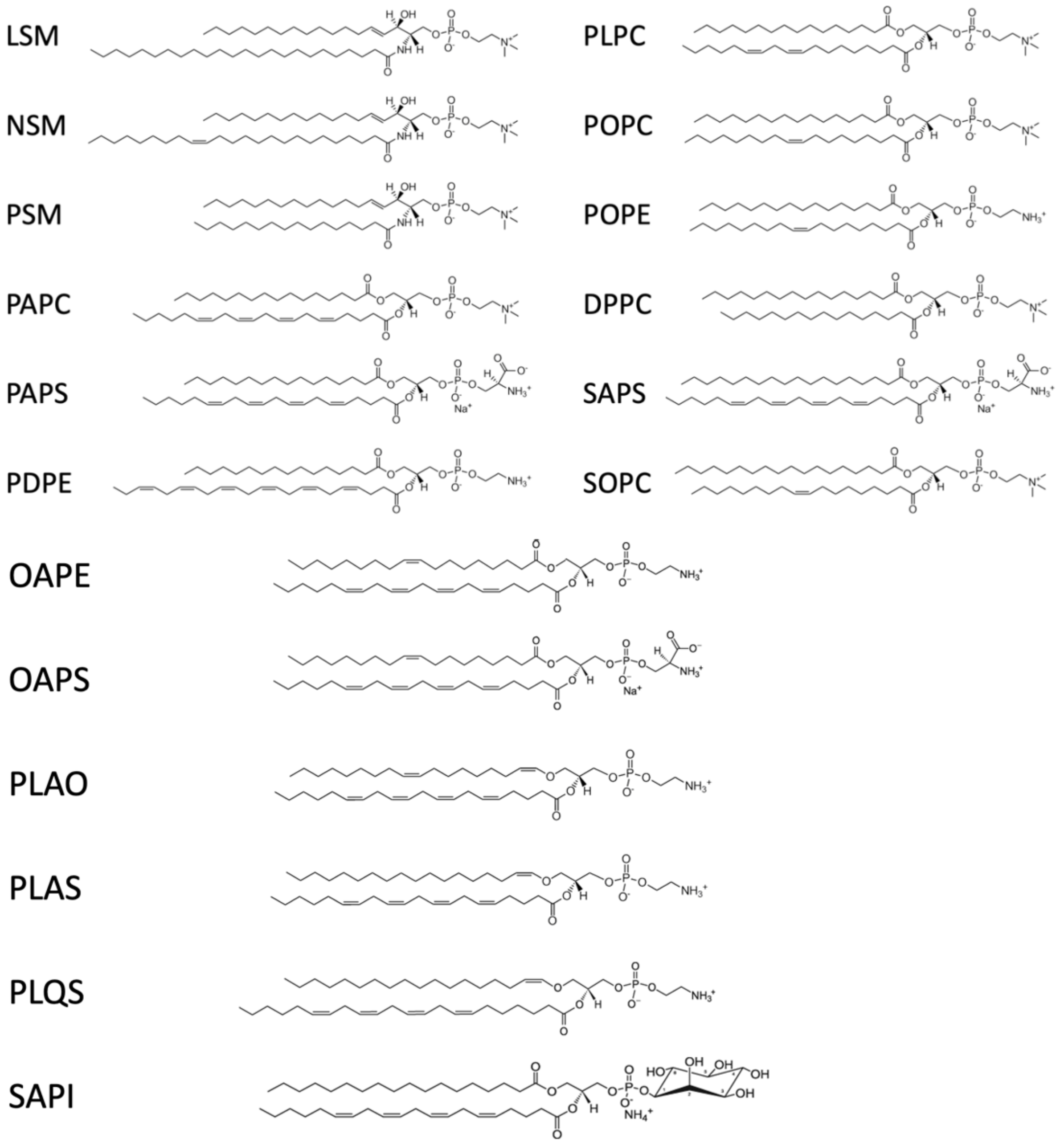


**Fig. S2. Names and structures of lipids used in the all-atom PM model simulations.**

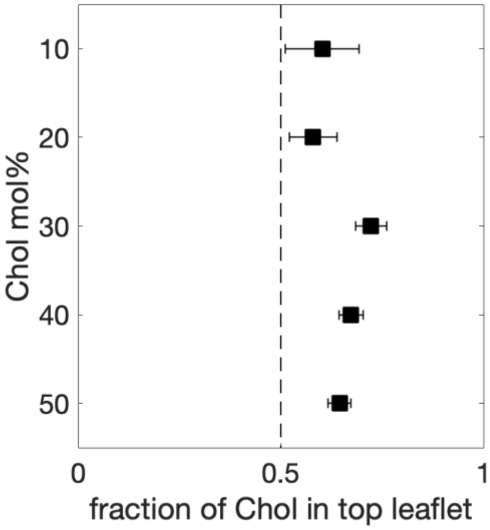


**Fig. S3. Equilibrated cholesterol distribution in coarse-grained PM models with fixed PL imbalance and varying total cholesterol abundance**. In all simulations (each 10$\mu$s-long), cholesterol accumulates in the 'exoplasmic' top sphingomyelin-rich leaflet. Specific leaflet lipid compositions and abundances are listed in Table S2.

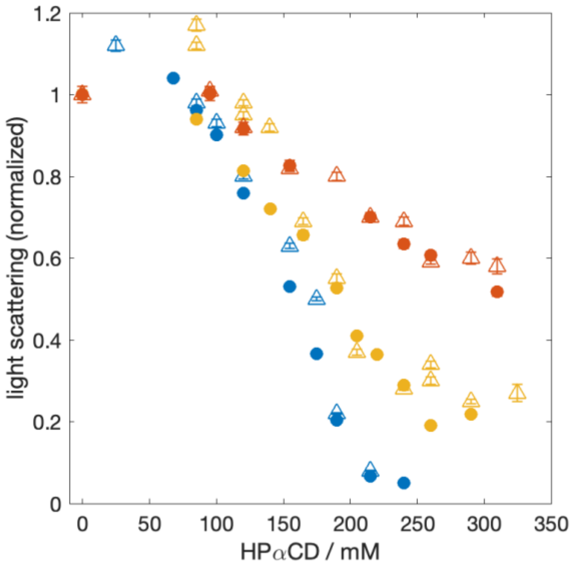


**Fig. S4. Light scattering of liposomes with increasing interleaflet PL imbalance induced by HP$\alpha$CD extraction.** Dynamic light scattering measured for extruded 100-nm liposomes composed of POPC with 0 (blue), 20 (yellow) and 40 (red) mol% cholesterol and incubated with increasing concentrations of hydroxy-propyl-alpha cyclodextrin (HP$\alpha$CD). HP$\alpha$CD extracts POPC lipids (but not cholesterol) from the outer leaflet of the membranes. A decrease in light scattering indicates destruction of the liposomes due to increased phospholipid imbalance. Cholesterol apparently increases tolerance for PL imbalance. Circles and triangles indicate independent replicate experiments.

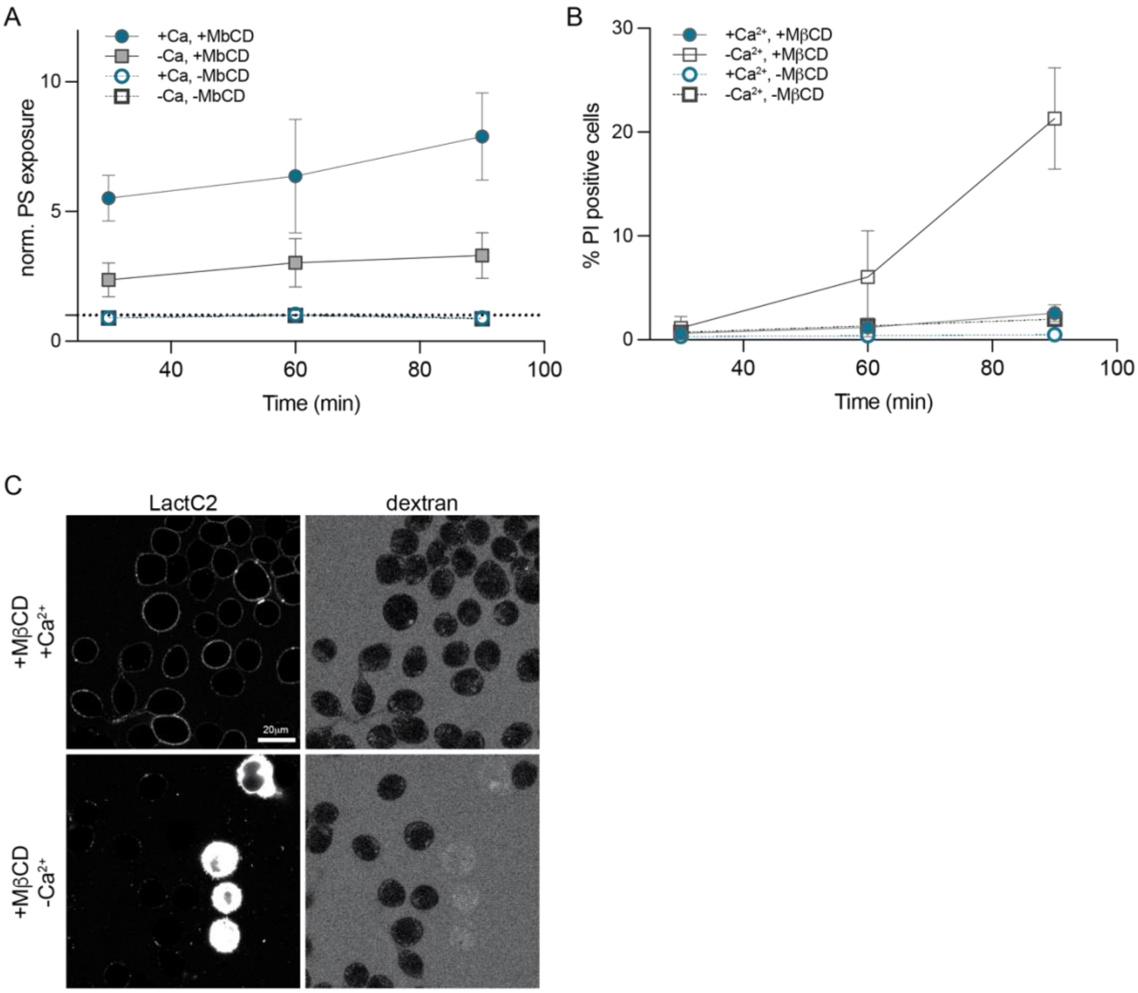


**Fig. S5. Plasma membrane scrambling and permeability of M$\beta$CD-treated cells.** (A) Time course of PS exposure on the surface of RBL cells treated with M$\beta$CD either in the presence or absence of calcium. PS exposure measured via binding of PS-marker LactC2-mClover and normalized to the respective –M$\beta$CD conditions. (B) Time course of membrane integrity in RBL cells treated with M$\beta$CD in the presence or absence of calcium. Membrane integrity was measured by percentage of cells with propidium iodide (PI) stained nuclei. PI is not membrane permeable and therefore only stains cells with disrupted PM integrity. Saponin treatment was used as a positive control, leading to 100% of cells become PI-positive. (C) Membrane disruption can also be measured by leakage of fluorescent dextran into the cytoplasm. Images show RBL cells treated with M$\beta$CD either in the presence (top) or absence (bottom) of calcium for 90 minutes at 23$^o$C. The former results in PS exposure and preserved membrane integrity as evidenced by impermeability to dextran. M$\beta$CD treatment in the absence of calcium suppresses PS exposure and compromises membrane integrity, with some cells becoming permeable both to dextran and LactC2 (which has a similar size).

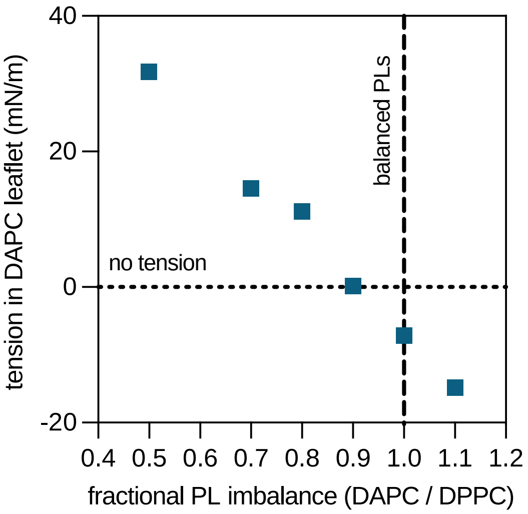


**Fig. S6. Leaflet tension in asymmetric DAPC/DPPC membranes with cholesterol and varying PL imbalance.** Asymmetric bilayers with one fully saturated DPPC leaflet opposing a highly unsaturated DAPC leaflet with varying relative number of lipids. Cholesterol re-distributes due to both its preference for saturated lipids and the PL imbalances between leaflets, as evidenced from long coarse-grained simulations (Figure 3B in main text). Fully atomistic trajectories of the equilibrated bilayer compositions (Table S4) were used to calculate the tension in all membrane leaflets. As the relative PL abundance in the DAPC leaflet (i.e. number of DAPC / number of DPPC) decreases, the tension in the DAPC leaflet increases.

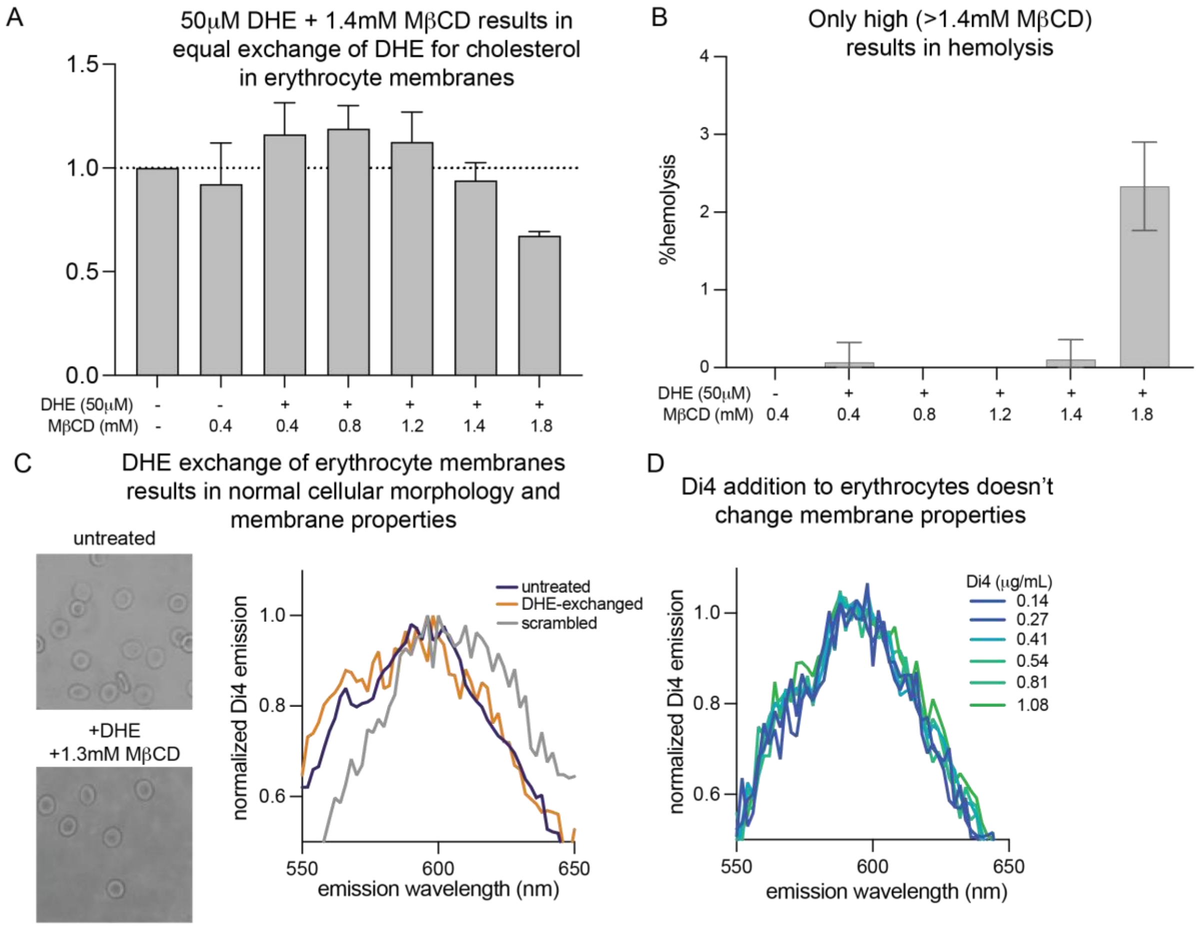


**Fig. S7. Optimization of DHE exchange conditions in erythrocytes.** (A) Incubation of 50 $\mu$M DHE with 1.2-1.4 mM M$\beta$CD yields a 1-to-1 exchange of DHE for cholesterol and maintains integrity of PM sterol composition as determined by Amplex Red of erythrocyte membranes after exchange. (B) M$\beta$CD concentrations greater than 1.4 mM result in erythrocyte hemolysis. (C) Erythrocytes with DHE exchanged for cholesterol at optimized conditions (1.3 mM MBCD + 50 $\mu$M DHE) do not exhibit altered cellular morphology (left) or lipid packing as measured by lack of shift in Di4 emission spectra (right). Grey line shows erythrocytes scrambled by treatment with 100 mM PMA for 10 minutes to illustrate a perturbation in membrane properties. (D) Addition of Di4 to erythrocytes does not change membrane properties as indicated by lack of shift in Di4 emission spectra.

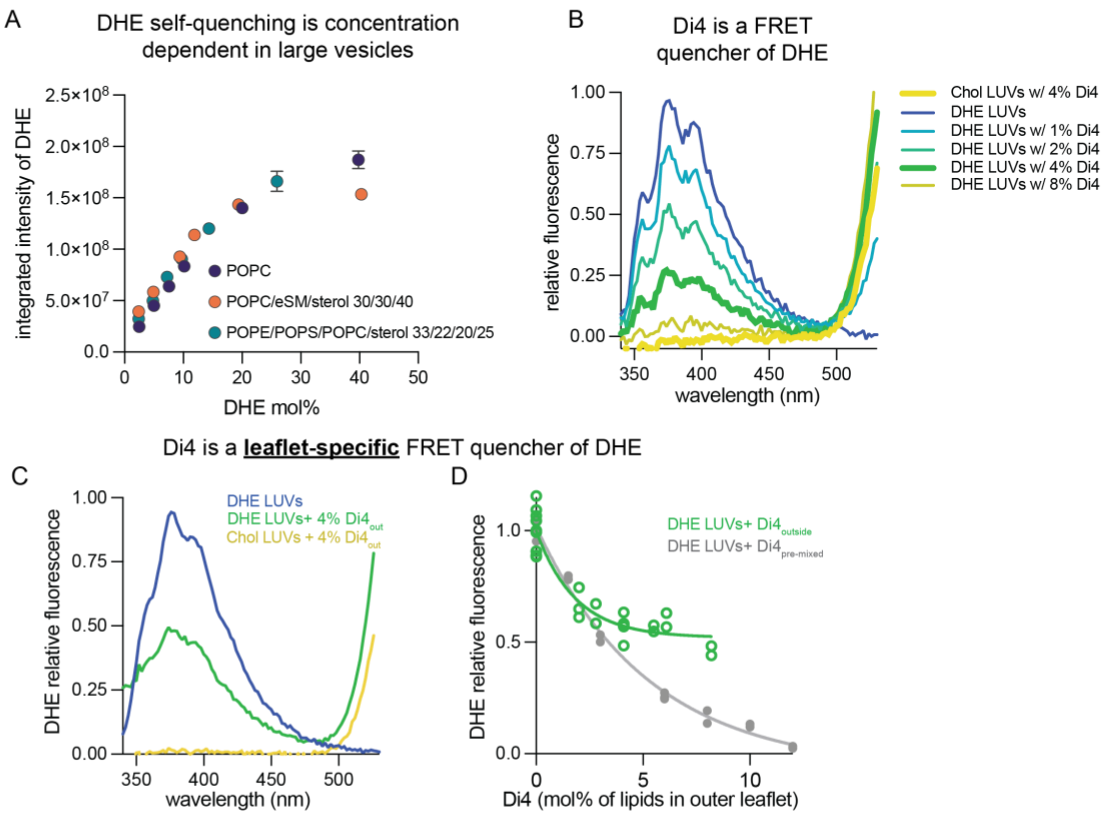


**Fig. S8. Quenching of DHE.** (A) DHE fluorescence in large liposomes with increasing amounts of DHE. The liposomes were prepared with the indicated lipid compositions where the added DHE either replaced part of the lipid (for POPC-only) or the sterol (for the multi-component mixtures) in the membrane. All lipid and DHE were mixed in organic solvent and prepared via rapid solvent exchange as detailed in Section 3.2. Since increasing amounts of DHE result in changes in the shape of the emission spectra, the integrated intensity of the full spectra is used for comparison between the samples. A clear linear regime is observed up to ~10-15 mol% DHE followed by a deviation from it indicating DHE self-quenching at higher concentrations. This behavior is independent of lipid composition as it is observed in mixtures with a wide range of fluidity. (B) Effect of Di4 on DHE fluorescence in extruded large unilamellar vesicles composed of 40% sterol (including 2.5% DHE), 5% POPG, and 55% (DOPC+Di4). Controls with Chol (instead of DHE) and DHE with no Di4 show the range of DHE fluorescence between 350 nm and 500 nm. Increasing amounts of Di4 (premixed with the lipids) reduce DHE fluorescence until it is barely detectable at Di4:DHE of ~3:1. (C) Externally added Di4 at 8 mol% of outer leaflet lipids quenches roughly half of the 2.5 mol% DHE symmetrically distributed in the LUVs composed of 40% sterol, 5% POPG and 55% DOPC. (D) Increasing amounts of Di4 added externally to LUVs progressively reduces DHE fluorescence measured at 385 nm, plateauing at ~50%. The data demonstrates that when added from the outside Di4 preferentially quenches outer leaflet DHE.

<u>Discussion of previous measurements of DHE asymmetry:</u> Cholesterol asymmetry has proven difficult to define because sterols rapidly flip-flop between leaflets (Bennett et al., 2009; Bruckner et al., 2009; Steck and Lange, 2012; Steck et al., 2002). This rapid equilibration presents a unique difficulty in trying to measure cholesterol's residence in a single leaflet, since each cholesterol

molecule samples both leaflets frequently (Lange et al., 1981). Unsurprisingly, previous reports have diverged dramatically: a report in the 1970s claimed ~3-fold enrichment in the outer leaflet (Fisher, 1976), contradicted 15 years later by claims of 3-fold enrichment in the cytoplasmic leaflet (Schroeder et al., 1991). More recently, evidence for cytoplasmic (Igbavboa et al., 1996; Mondal et al., 2009; Solanko et al., 2018), exoplasmic (Bittman and Rottem, 1976; Hale and Schroeder, 1982; Ingolfsson et al., 2014), or neither (Lange and Slayton, 1982) leaflet being cholesterol enriched has been presented. In recent years, two high-profile papers again presented conflicting claims of either strong inner PM leaflet enrichment (Courtney et al., 2018) or almost exclusive enrichment in the outer leaflet (Liu et al., 2017).

Methods relying on DHE quenching, similar to our approach, have also been used previously to infer cholesterol asymmetry. In both yeast and mammalian cells, quantifying DHE fluorescence intensity by imaging cells before and after addition of external DHE quenchers was used to conclude that most DHE (and by inference, most cholesterol) resided in the cytoplasmic PM leaflet (Mondal et al., 2009; Solanko et al., 2018). The yeast experiments were cleverly constructed to allow essentially all (native) ergosterol to be replaced by fluorescent DHE. However, one potential issue with this approach is the strong self-quenching of DHE at high concentrations, which has been previously reported (Schroeder et al., 1987) and which we reproduce here (Fig S8A). Such self-quenching is likely to affect conclusions based on external quenching in cell PMs and this problem would be especially acute if sterol is asymmetrically distributed. An extreme, illustrative case would be one leaflet with high DHE concentration leading to complete self-quenching, while the other leaflet has low, sub-self-quenching concentration. In this case, most (or even all) observable fluorescence could come from the leaflet with *lower* DHE concentration. In our experiments, we took care to avoid the possibility of self-quenching by minimizing the fraction of DHE exchanged into erythrocytes, keeping it below 10% of all PM lipids.

Furthermore, previous experiments often relied on highly reactive oxidizing quenchers like trinitrobenzenesulfonic acid (TNBS), which covalently react with biomolecules and lead to cellular toxicity. We used a bio-orthogonal fluorophore (Di4) as the fluorescence quencher, with the added benefit that Di4 also reports on changes in membrane properties (or lack thereof in our measurements – see Fig S7C-D). However, while our measurements strongly support enrichment of sterol in the outer PM leaflet, they do not provide absolute quantification of molar enrichment between leaflets (see SI Section 4.2). Absolute quantification would require detailed information about how the spectral properties of both the donor (DHE) and acceptor (Di4) depend on various membrane environments, absolute quantifications of their lateral density in erythrocyte membranes, and careful modeling to interpret and frame the results. Such characterization is outside the scope of the present study.

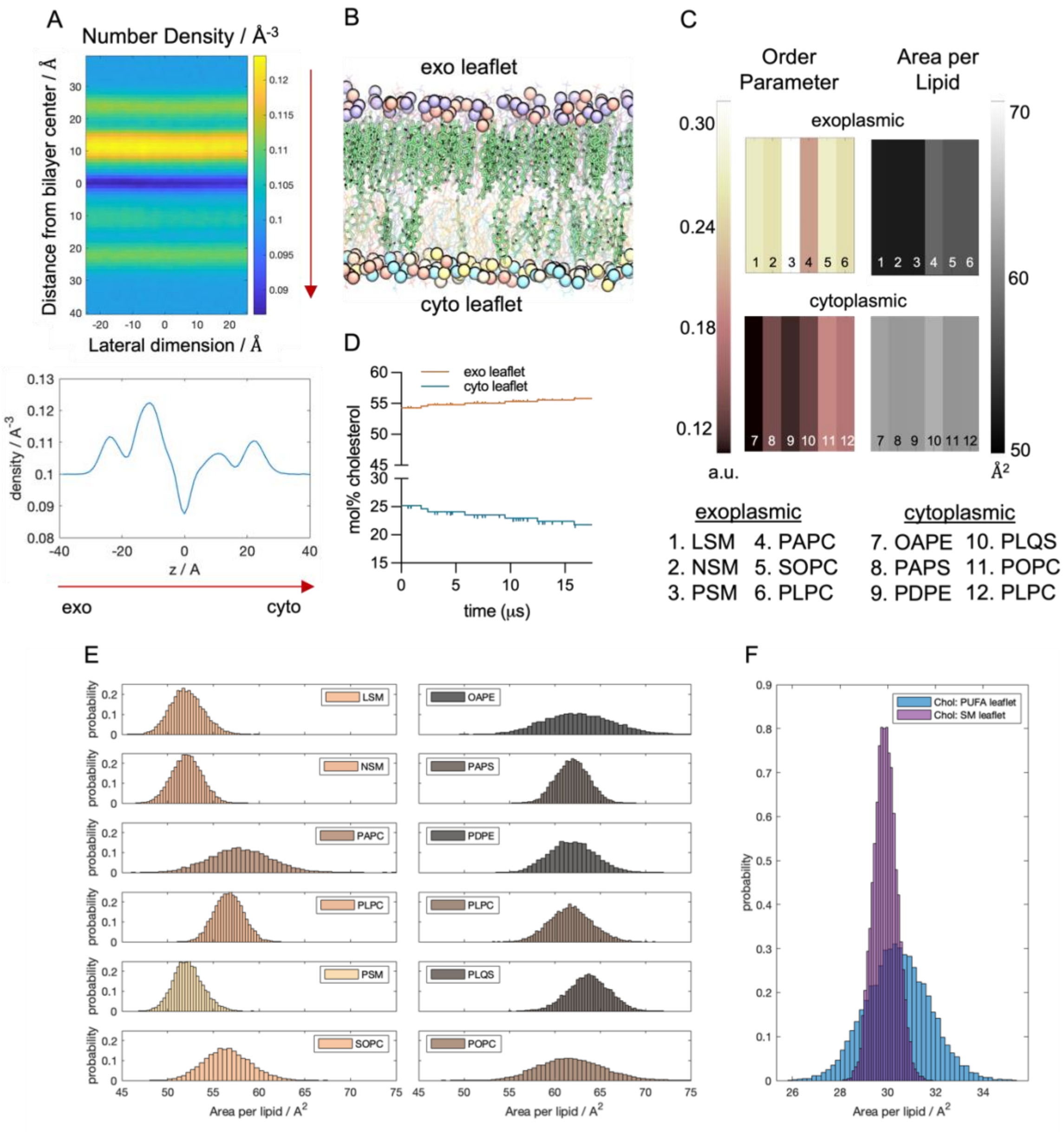


**Fig. S9. Structural properties of Cyto+ model.** (A) (top) Number density of all atoms (including lipids and water) calculated in 3D and projected onto the xz plane. Y-axis of the heatmap shows distance from bilayer center and from top to bottom goes through water region, exoplasmic leaflet, bilayer center, cytoplasmic leaflet and water region, as indicated by the red arrow (as shown in panel (B) for reference). (bottom) Laterally averaged number density profile for more direct quantitative comparison of the density peaks. (B) Snapshot of the equilibrated simulated Cyto+ membrane (as detailed in the caption of Figure 1D in main text). Water atoms not shown. (C) Average order parameter (left) and area per lipid (right) for each type of lipid in the exoplasmic (top) and cytoplasmic (bottom) leaflets. Order parameter was averaged over all carbon atoms on both chains. Area per lipid was calculated from Voronoi tessellation analysis detailed in the SI text. (D) Cholesterol concentration (expressed as mol% of each leaflet) in the Cyto+ membrane shows the time evolution of cholesterol interleaflet distribution. (E) Histograms of individual lipid areas for exoplasmic (left) and cytoplasmic (right) leaflet lipids whose averages are shown in the heatmaps in panel (C). (F) Histograms of the lipid areas of cholesterol (Chol) in the exoplasmic (SM) leaflet and cytoplasmic (PUFA) leaflet calculated from Voronoi analysis in every frame of the trajectory. The analyses in (B–C) and (E–F) are from the last 1.2 $\mu$s of the trajectory.

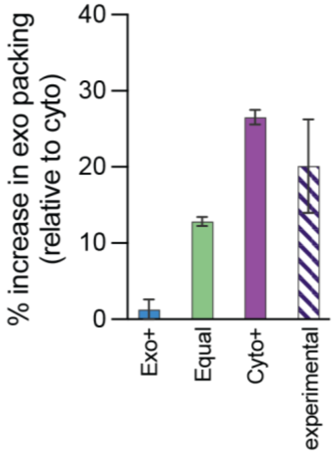


**Fig. S10. Experimental validation of lipid packing in Cyto+ model.** Comparison of the area/lipid difference between exoplasmic and cytoplasmic leaflets in simulations versus experiments (shown as percent increase in the packing: ($APL_{cyto}$ - $APL_{exo}$) / $APL_{cyto}$). Experimental measurements represent APLs inferred from Di4 lifetime measurements in fibroblast 3T3 cells and the calibration curve in Figure 4A in main text. The Cyto+ model is within error of experiment, while the Exo+ and Equal models are significantly different.

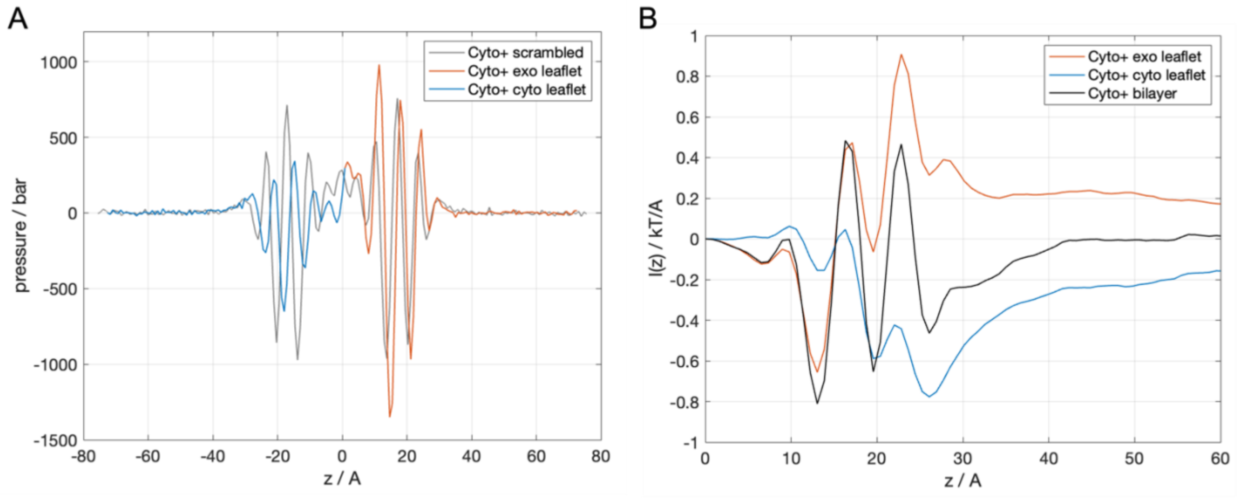


**Fig. S11. Lateral pressure profile and bilayer torque density of Cyto+ membrane.** (A) Lateral pressure profiles of the asymmetric Cyto+ bilayer (with the exo and cyto leaflets colored red and blue, respectively) and its symmetric scrambled counterpart (gray). Scrambling membrane lipids equalizes the lipid compositions of the two leaflets and makes the pressure distribution on the two sides of the bilayer midplane (z=0) the same. (B) First moment of the pressure profile for each leaflet in the Cyto+ membrane and their sum, i.e. for the bilayer. The plateau values far from the membrane (i.e. at large z values) represent the torque densities of either individual leaflets or the whole bilayer. Torque density quantifies the propensity for a membrane to curve, with positive values representing positive curvature and vice versa. Torque density close to zero implies a bilayer with a tendency to remain flat. In the Cyto+ bilayer, although both individual monolayers have a significant torque density, the net bilayer torque density is essentially 0, indicating that differential and curvature stresses in the membrane are balanced and result in preference for flat morphology.

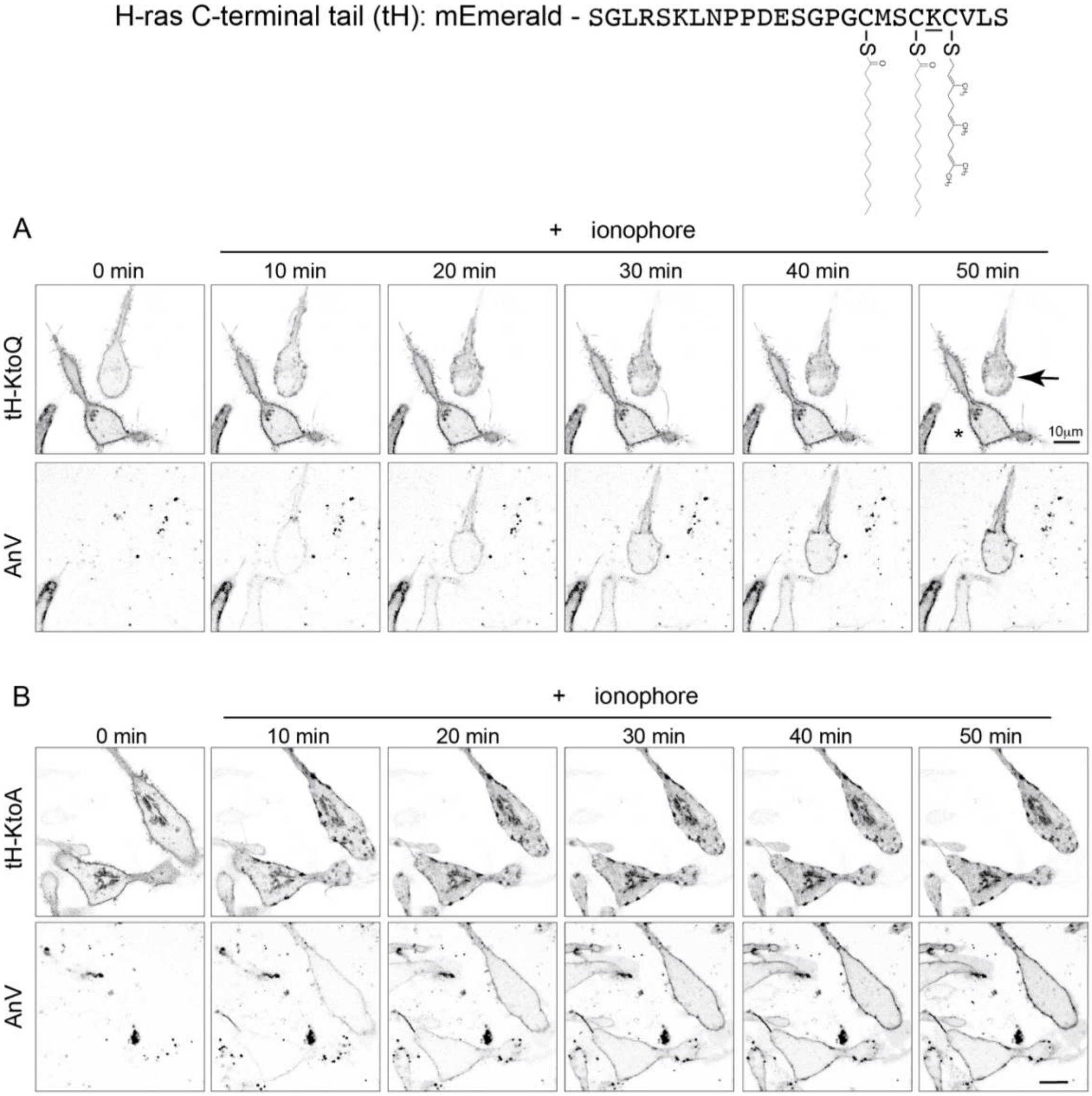


**Fig. S12. Effects of plasma membrane lipid scrambling on the interaction of lipidated peptides with the membrane's cytoplasmic leaflet.** Confocal images of the cellular distribution of two variants of the tH-Ras peptide in RBL cells following membrane scrambling. tH-Ras is composed of the 20 C-terminal amino acids of H-Ras, which contains its farnesylation signal (CAAX) and two sites for palmitoylation – thus this peptide has up to 3 lipid anchors. The peptide is tagged with mEmerald for visualization. To eliminate the possibility of electrostatic interactions, we mutated the Lysine of the peptide which is located between the farnesyl and palmitoyl chains to either glutamine (A) or alanine (B). The sequence of the probe is given, with the acylation sites noted and mutated lysine underlined. Scrambling of PM lipids was initiated with the ionophore A23187 while exposure of PS on the cell surface was monitored with Annexin V (AnV) Alexa Fluor 647. Both variants of the peptide localized largely to the PM at steady-state but redistributed to inner membranes upon release of asymmetry (indicated by AnV binding). The favorable interaction with the asymmetric PM is consistent with hydrophobic defects present in the cytoplasmic leaflet (Figure 5E-F) which promote the insertion of the lipid anchors and disappear after PM scrambling. As a control, in (A) the cell marked with an asterisk does not scramble during this time course, and the peptide remains at the plasma membrane; in contrast, the cell marked with an arrowhead does scramble and the peptide is redistributed to inner membranes.

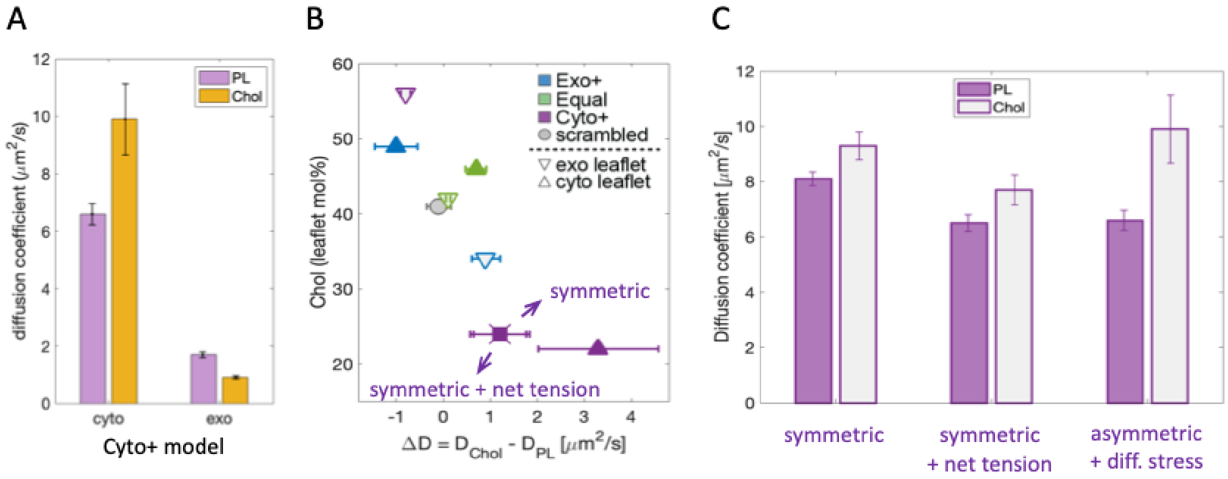


**Fig. S13. Determinants of fast cholesterol diffusion.** (A) Phospholipid (PL) and cholesterol (Chol) diffusion in each leaflet of the Cyto+ membrane. (B) Difference between cholesterol ($D_{Chol}$) and phospholipid ($D_{PL}$) diffusion ΔD in each leaflet of the three PM models and the Cyto+ scrambled membrane is plotted versus cholesterol concentration in the leaflet. A high correlation between the two shows that the cytoplasmic leaflet of the Cyto+ model, which is differentially stressed and has tension of ~8 mN/m, has the largest ΔD. A symmetric membrane with the same lipid composition has a smaller ΔD both in the absence (symmetric) or presence of 8 mN/m net tension (symmetric + net tension). These results indicate that the fast diffusion of cholesterol relative to PLs is related to both low cholesterol concentration in the leaflet and tension arising from differential stress. (C) PL and Chol diffusion coefficients in mixtures having the lipid and cholesterol composition of the cytoplasmic leaflet of the Cyto+ model: symmetric membrane with no differential stress or tension (left), symmetric membrane with applied net tension (middle), and the cyto leaflet of the Cyto+ model which has differential stress of the same magnitude (right). These results show that tension of 8 mN/m reduces PL diffusion to the same extent, irrespective of its source; however, Chol diffusion is only slightly affected by differential stress and to a much greater extent by net bilayer tension. Thus, differential stress contributes to a large ΔD by its differential effects on PL and Chol diffusion.

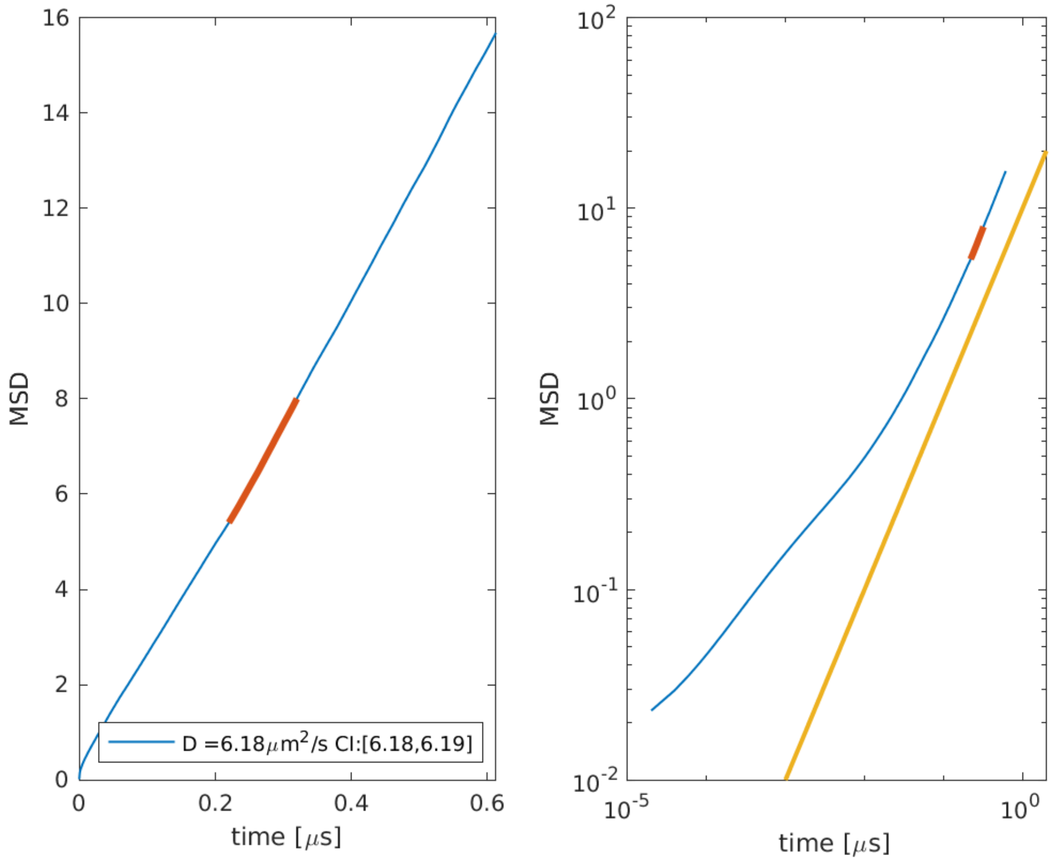


**Fig. S14. Calculation of diffusion coefficients from simulations.** Representative plots showing mean square displacement (MSD) as a function of time in a linear (left) and a log-log (right) scale. Red indicates the 100-ns block from which the diffusion coefficient was calculated (see SI text, section 3.3). Data shown is for the phospholipids in the cytoplasmic leaflet of the Cyto+ model.

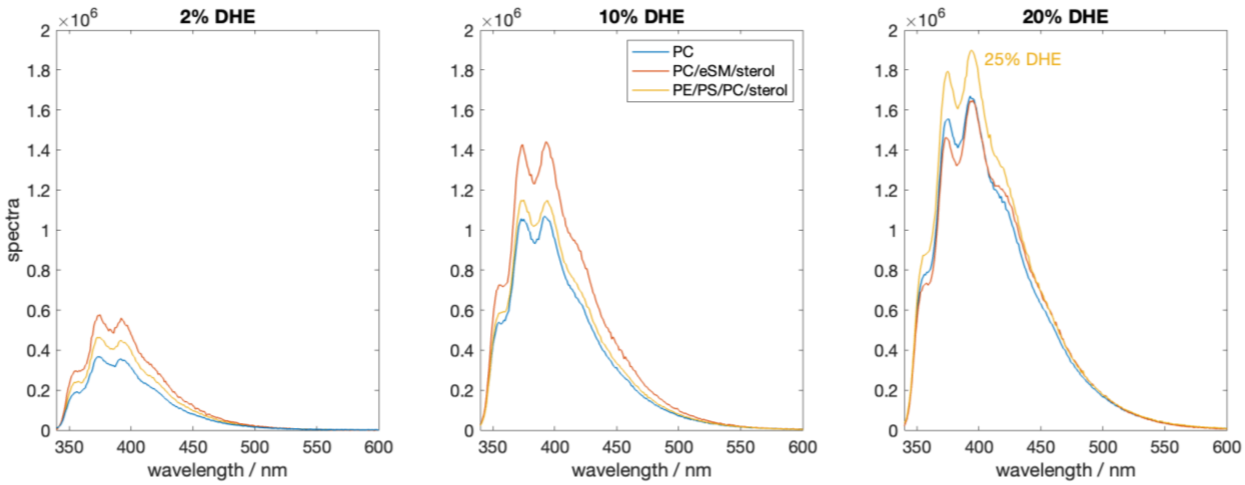


**Fig. S15. Changes in DHE spectra in liposomes at higher DHE concentrations.** Representative spectra of DHE in liposomes (large vesicles prepared by rapid solvent exchange) composed of POPC (PC), egg sphingomyelin (eSM), POPE (PE), POPS (PS) and cholesterol (see Figure S8A for detailed lipid compositions) with increasing amounts of DHE indicated in the title of each plot. The shape of the spectra changes at higher DHE concentrations, making trends with composition or DHE content dependent on the choice of wavelength. This problem can be circumvented by integration of the spectra.

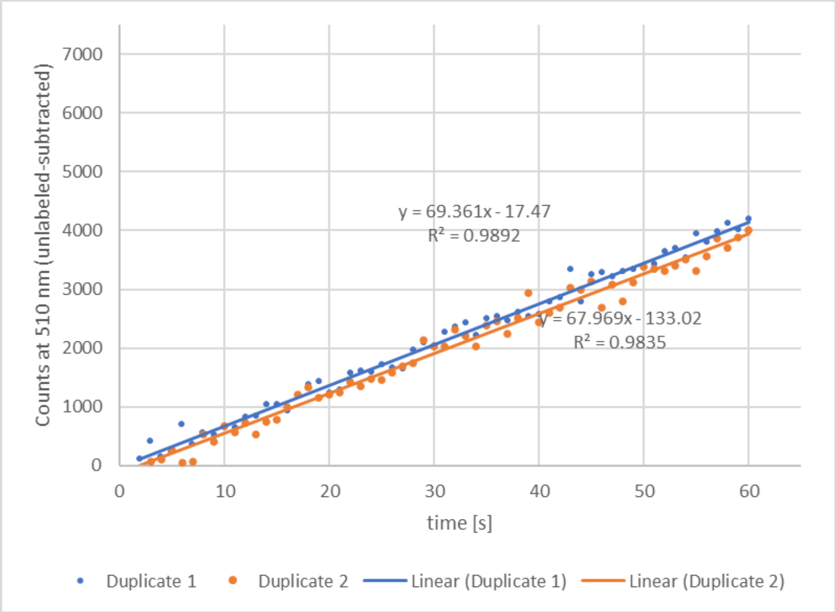


**Fig. S16. Erythrocyte permeability measurements.** Flux of FDA across the PM of intact untreated erythrocytes is measured via the slope of linear increase in fluorescein fluorescence (due to cytoplasmic FDA hydrolysis). Two representative replicates are shown with linear fits.

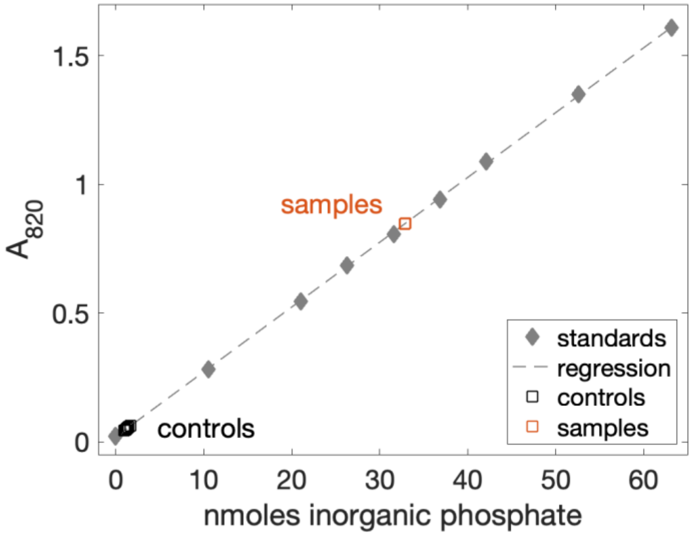


**Fig. S17. Quantification of phosphate.** Calibration curve of phosphate standards (diamonds) used to quantify the nanomoles lipid in a sample (orange squares) relative to controls (black squares). Data shown is for two replicates of MαCD solution after incubation with cells to extract lipids. Corresponding controls were done with the same conditions but in the absence of MαCD.

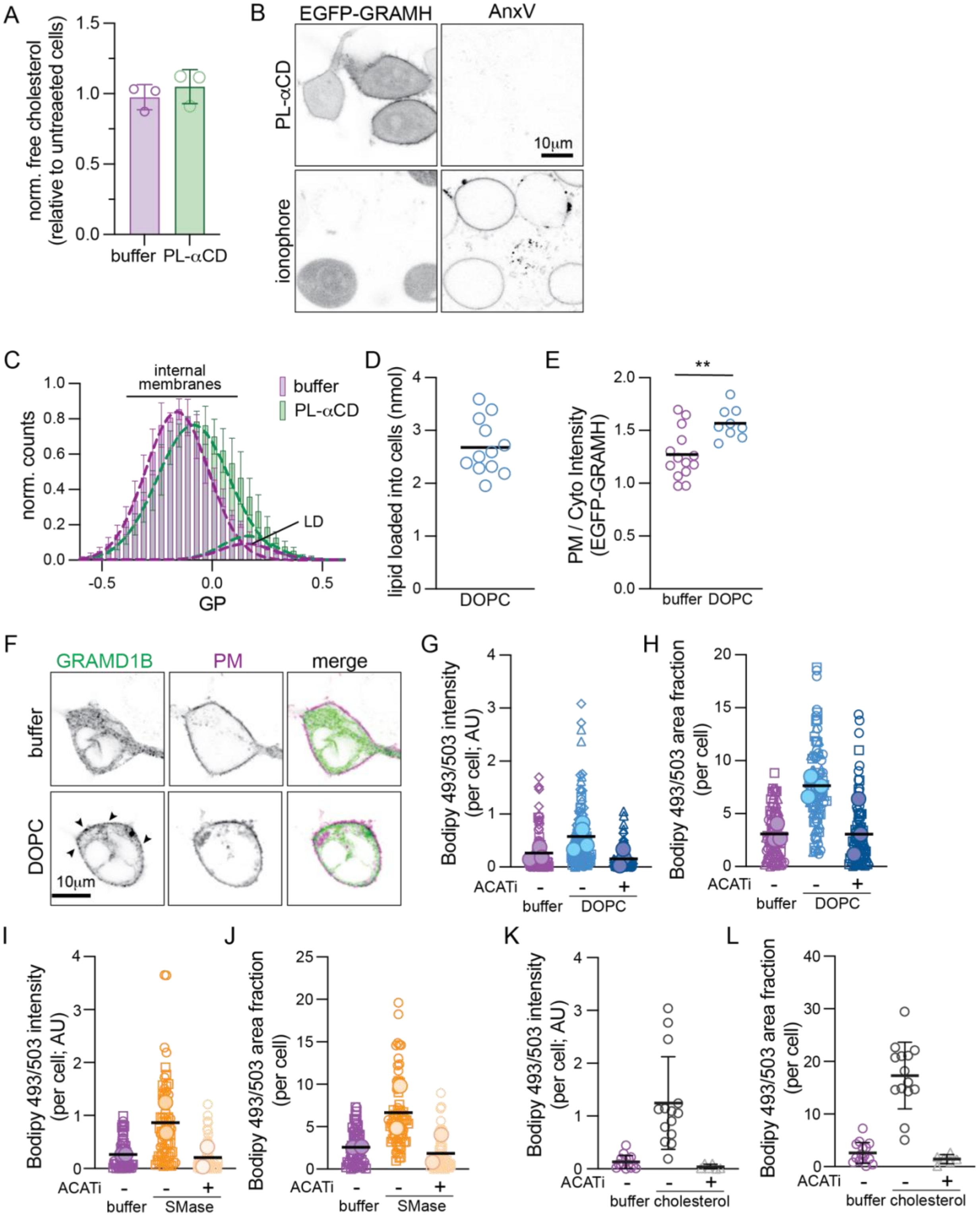


**Fig. S18. Manipulating PM outer leaflet phospholipid abundance induces CE lipid droplets and internal membrane ordering.** (A) Cholesterol abundance in cells treated with PL-aCD relative to protein amount and normalized to untreated cells. Analyzed by AmplexRed assay; data points represent independent experiments. (B) AnnexinV (AnxV) staining of cells treated with PL-aCD or 3 μM A23187 as a positive control. PL-aCD treatment does not induce PS scrambling as indicated by a lack of AnxV external staining. (C) Histograms of C-laurdan GP in cells. To determine the GP of the internal membranes (i.e. non-lipid droplet), a histogram of all pixels excluding the PM was generated (for each experiment), and the data fit to two Gaussian peaks (with one mean fixed at GP > 0.1, representing the very high GP of LDs). PL-aCD treatment results in a higher amplitude for the LD peak (more LD) and a higher mean for the internal membranes (tighter lipid packing). (D) The abundance of DOPC loaded into cells quantified by phosphate assay. (E) EGFP-GRAMH translocation to the PM in DOPC-loaded cells. (F) Imaging of RBL cells expressing full-length EGFP-GRAMD1B and trLAT (PM marker) with and without DOPC treatment. Arrowheads show translocation of GRAMD1B to the PM after DOPC treatment. (G-H) Quantification of LDs in DOPC-treated cells with and without ACAT inhibition (1 ug/mL Sandoz 58-035). (I-J) Quantification of LD in SMase-treated cells with and without ACAT inhibition (1 ug/mL Sandoz 58-035). (K-L) Quantification of LD in cholesterol-loaded cells with and without ACAT inhibition (1 ug/mL Sandoz 58-035). Using maximum projection of volumetric imaging, the total intensity of staining (G, I, K) and area fraction covered by LDs (H, J, L) are reported.

**Table S1. Re-analyzed lipidomics data from literature.** Reported overall and leaflet phospholipid (PL) compositions for the plasma membrane of erythrocytes were used to calculate respective PL abundances in each leaflet. The data excludes lysoPC. Note that the sum of exo and endo lipids as mol% of total lipids is different from the total overall lipids for the cases where the asymmetry of PI was not reported. Last column shows the relative PL abundances in the cyto and exo leaflets with upper and lower bounds calculated assuming that the missing lipids (i.e. lysoPC or those whose asymmetry was not reported) are all in the exo or endo leaflet. Reported quantities are with white background, and calculated ones are shaded in color.

| Ref | Species | Leaflet | Mol% of lipid class | | | | | Mol% of total lipids | | | | | | Relative PL abundance cyto/exo |
|---|---|---|---|---|---|---|---|---|---|---|---|---|---|---|
| | | | SM | PC | PE | PI | PS | SM | PC | PE | PI | PS | Total PLs | |
| 1 | Human | exo- | 85 | 68 | 0 | - | 0 | 21.25 | 19.99 | 0.00 | - | 0.00 | 41.24 | 1.34 [1.23; 1.42] |
| | | cyto- | 15 | 32 | 100 | - | 100 | 3.75 | 9.41 | 27.00 | - | 15.00 | 55.16 | |
| | | overall | | | | | | 23.50 | 29.40 | 27.00 | 3.00 | 15.00 | 97.90 [a] | |
| 2 | Rat | exo- | 100 | 48 | 8 | 0 | | 12.00 | 20.16 | 2.00 | 0.00 | | 34.16 | 1.78 [1.55; 1.93] |
| | | cyto- | 0 | 52 | 92 | 100 | | 0.00 | 21.84 | 23.00 | 16.00 | | 60.84 | |
| | | overall | | | | | | 12.00 | 42.00 | 25.00 | 16.00 | | 95.00 | |
| 3 | Human | exo- | 85 | 68 | 0 | - | 0 | 19.98 | 19.99 | 0.00 | - | 0.00 | 39.97 | 1.37 [1.22; 1.5] |
| | | cyto- | 15 | 38 | 100 | - | 100 | 3.53 | 9.41 | 27.00 | - | 15.00 | 54.93 | |
| | | overall | | | | | | 23.50 | 29.40 | 27.00 | 3.00 | 15.00 | 97.90 [a] | |
| 4 | Mouse | exo- | 85 | 30 | 20 | 42 | 0 | 9.78 | 13.62 | 4.86 | 1.72 | 0.00 | 29.98 | 2.21 [1.95; 2.34] |
| | | cyto- | 15 | 70 | 80 | 58 | 100 | 1.73 | 31.78 | 19.44 | 2.38 | 10.80 | 66.12 | |
| | | overall | | | | | | 11.50 | 45.40 | 24.30 | 4.10 | 10.80 | 96.10 | |
| 5 | Monkey | exo- | 82 | 42 | 5 | 0 | 0 | 12.87 | 16.17 | 1.38 | 0.00 | 0.00 | 30.42 | 2.19 [1.98; 2.29] |
| | | cyto- | 18 | 58 | 95 | 100 | 100 | 2.64 | 22.33 | 26.22 | 5.10 | 10.20 | 66.49 | |
| | | overall | | | | | | 15.50 | 38.50 | 27.60 | 5.10 | 10.20 | 96.90 | |
| 6 | Human | exo- | 89 | 48 | 4 | 0 | 15 | 14.98 | 14.57 | 1.10 | 0.00 | 1.46 | 32.11 | 2.02 [1.84; 2.11] |
| | | cyto- | 11 | 52 | 96 | 100 | 85 | 1.85 | 15.78 | 26.51 | 1.26 | 19.41 | 64.81 | |
| | | overall | | | | | | 16.83 | 30.34 | 27.62 | 1.26 | 20.87 | 96.92 | |

[1] Verkleij et al. (Verkleij et al., 1973), [2] Renooij et al. (Renooij et al., 1974), [3] Zwaal et al. (Zwaal et al., 1975), [4] Rawyler et al. (Rawyler et al., 1985), [5] Van der Schaft et al. (Van der Schaft et al., 1987), [6] Lorent et al. (Lorent et al., 2020)

[a] SM and PC calculated from % degradations (i.e. from reported % SM and % total degraded); PE, PI and PS assumed based on relative compositions of RBCs

**Table S2. Leaflet compositions of CG bilayers used to test cholesterol effect on bilayer tolerance for PL mismatch.** Number of lipids used to construct asymmetric coarse-grained (CG) bilayers mimicking the cell PM and having varying Chol concentration. Each bilayer has 35% more phospholipids in the model cytoplasmic leaflet, and the same starting concentration of Chol in both leaflets. The corresponding lipid names in Martini are DPSM (PSM), PEPC (PLPC), PRPE (PDPE) and PAPS (PAPS). Shown are the numbers of Chol molecules in each leaflet at the beginning of the simulation.

| system | exoplasmic leaflet | | | | cytoplasmic leaflet | | | | |
|---|---|---|---|---|---|---|---|---|---|
| | PSM | PLPC | Chol | total | PDPE | PAPS | PLPC | Chol | total |
| A10 | 45 | 45 | 10 | 100 | 40 | 40 | 41 | 14 | 135 |
| A20 | 40 | 40 | 20 | 100 | 36 | 36 | 36 | 27 | 135 |
| A30 | 35 | 35 | 30 | 100 | 31 | 31 | 32 | 41 | 135 |
| A40 | 30 | 30 | 40 | 100 | 27 | 27 | 27 | 54 | 135 |
| A50 | 25 | 25 | 50 | 100 | 22 | 22 | 23 | 68 | 135 |

**Table S3. All-atom simulations for studying the effect of PS flipping on leaflet tension.** Shown are the numbers of POPC, POPS, and Chol in each leaflet of the simulated bilayers and the effective number and type of *flipped* lipids modeled by the system (POPS is always "flipped" from the top to the bottom leaflet while Chol is "flopped" from the bottom to the top leaflet). The total length of the respective trajectories and the length of the last portion used for analysis are indicated in the last two columns (the latter was chosen based on equilibration of the bilayer area). Each bilayer was hydrated with 45 waters per lipid and charge-neutralized with potassium ions.

| **bilayer** | **# lipids (POPC/POPS/Chol)** | | **# flipped lipids** | | **Simulation length / ns** | |
|---|---|---|---|---|---|---|
| | **top** | **bottom** | **POPS** | **Chol** | **analysis** | **total** |
| **0PS** | 64 / 16 / 20 | 64 / 16 / 20 | 0 | 0 | 460 | 960 |
| **1PS** | 64 / 15 / 20 | 64 / 17 / 20 | 1 | 0 | 650 | 900 |
| **1PS-1Ch** | 64 / 15 / 21 | 64 / 17 / 19 | 1 | 1 | 910 | 1050 |
| **1PS-2Ch** | 64 / 15 / 22 | 64 / 17 / 18 | 1 | 2 | 806 | 806 |
| **5PS** | 64 / 11 / 20 | 64 / 21 / 20 | 5 | 0 | 500 | 750 |
| **5PS-5Ch** | 64 / 11 / 25 | 64 / 21 / 15 | 5 | 5 | 970 | 1020 |
| **5PS-10Ch** | 64 / 11 / 30 | 64 / 21 / 10 | 5 | 10 | 880 | 1013 |
| **5PS-13Ch** | 64 / 11 / 33 | 64 / 21 / 7 | 5 | 13 | 806 | 806 |

**Table S4. Coarse-grained and all-atom simulations for investigating the determinants of cholesterol distribution in lipid bilayers.** Name of each system, initial numbers of lipids in each leaflet in the CG simulations, and the equilibrated numbers of lipids from the 10-$\mu$s-long trajectories. The latter were used to construct corresponding AA bilayers whose leaflet compositions did not change in the course of the production runs. Both the total length of the respective trajectories and the length of the last portion used for analysis are indicated for each simulation. Bilayers A.12 and A.13 did not tolerate the lipid number mismatch between the leaflets and exhibited an altered morphology as shown in Figure 2A in the main text. They were not used for subsequent simulation and analysis in AA representation.

| **bilayer** | **start CG (DPPC/DAPC/Chol)** | | **end CG / start AA (DPPC/DAPC/Chol)** | | **AA simulation length / ns analysis / total** |
|---|---|---|---|---|---|
| | **top** | **bottom** | **top** | **bottom** | |
| A.5 | 70 / 0 / 30 | 0 / 35 / 15 | 70 / 0 / 29 | 0 / 35 / 16 | 400 / 1260 |
| A.7 | 70 / 0 / 30 | 0 / 49 / 21 | 70 / 0 / 37 | 0 / 49 / 14 | 392 / 1130 |
| A.8 | 70 / 0 / 30 | 0 / 56 / 24 | 70 / 0 / 43 | 0 / 56 / 11 | 400 / 1102 |
| A.9 | 70 / 0 / 30 | 0 / 63 / 27 | 70 / 0 / 47 | 0 / 63 / 10 | 400 / 1060 |
| A.10 | 70 / 0 / 30 | 0 / 70 / 30 | 70 / 0 / 52 | 0 / 70 / 8 | 400 / 1040 |
| A.11 | 70 / 0 / 30 | 0 / 77 / 33 | 70 / 0 / 56 | 0 / 77 / 7 | 400 / 1077 |
| A.12 | 70 / 0 / 30 | 0 / 84 / 36 | - | - | - |
| A.13 | 70 / 0 / 30 | 0 / 91 / 39 | - | - | - |

**Table S5. Phospholipid and cholesterol compositions of simulated PM models.** The compositions of three simulated PM models (Cyto+, Equal, Exo+), with types and numbers of lipids in each leaflet, as well as the corresponding number and leaflet mol% of cholesterol at the beginning and end of the simulations. Figure S2 shows the structures and full names of all lipids.

<table>
<tr><th rowspan="2">model</th><th colspan="2">exoplasmic leaflet</th><th colspan="2">cytoplasmic leaflet</th><th colspan="2">cholesterol exo/cyto number<br>(leaflet mol%)</th></tr>
<tr><th>lipid</th><th>number</th><th>lipid</th><th>number</th><th>beginning</th><th>end</th></tr>
<tr><td>Cyto+</td><td>LSM<br>NSM<br>PSM<br>PAPC<br>PLPC<br>SOPC</td><td>12<br>14<br>18<br>6<br>20<br>10</td><td>OAPE<br>PAPS<br>PDPE<br>PLQS<br>PLPC<br>POPC</td><td>8<br>30<br>18<br>20<br>20<br>8</td><td>95/35<br>(54/25)</td><td>101/29<br>(56/22)</td></tr>
<tr><td>total</td><td></td><td>80</td><td></td><td>104</td><td colspan="2">130</td></tr>
<tr><td>Equal</td><td>LSM<br>NSM<br>PSM<br>PAPC<br>PLAS<br>PLPC<br>SAPS<br>SOPC</td><td>54<br>63<br>81<br>36<br>18<br>99<br>9<br>45</td><td>DPPC<br>PSM<br>OAPE<br>OAPS<br>PAPS<br>PDPE<br>PLAO<br>PLPC<br>PLQS<br>POPC<br>POPE<br>SAPI</td><td>16<br>8<br>25<br>8<br>82<br>49<br>16<br>57<br>57<br>25<br>16<br>16</td><td>267/261<br>(40/41)</td><td>294/234<br>(42/38)</td></tr>
<tr><td>total</td><td></td><td>405</td><td></td><td>375</td><td colspan="2">528</td></tr>
<tr><td>Exo+</td><td>LSM<br>NSM<br>PSM<br>PAPC<br>PLPC<br>SOPC</td><td>16<br>18<br>22<br>8<br>28<br>12</td><td>OAPE<br>PAPS<br>PDPE<br>PLQS<br>PLPC<br>POPC</td><td>6<br>22<br>14<br>16<br>16<br>6</td><td>35/95<br>(25/54)</td><td>54/76<br>(34/49)</td></tr>
<tr><td>total</td><td></td><td>104</td><td></td><td>80</td><td colspan="2">130</td></tr>
</table>

**Table S6. Trajectory lengths of fully atomistic PM models.** Production runs of the PM models and their symmetric counterparts were simulated first on the Anton supercomputer with the Desmond software, then with NAMD. Short NAMD trajectories with Langevin damping coefficient $\gamma = 5$ / ps were followed by longer ones with $\gamma$ set to 1 / ps; last 1.2 $\mu$s (or 1 $\mu$s for Cyto+ cyto-sym) were used for most subsequent analyses. Last column shows the total length of the last portion of the NAMD trajectories with $\gamma = 1$ / ps used for calculation of lipid and cholesterol diffusion coefficients.

| bilayer | initial anton run / $\mu$s | subsequent NAMD run / $\mu$s | | diffusion analysis / $\mu$s |
|---|---|---|---|---|
| | | $\gamma = 5$ / ps | $\gamma = 1$ / ps | |
| **Cyto+** | 15 | 0.4 | 2 | 2 |
| **Equal** | 25 | - | 1.51 | 1.5 |
| **Exo+** | 22.8 | - | 3.15 | 2.7 |
| **Cyto+ exo-sym** | 3 | - | 3 | 2.5 |
| **Cyto+ cyto-sym** | 3 | - | 1 | 1 |
| **Cyto+ scramble** | 5 | - | 1.4 | 1.24 |
